# High-resolution influenza mapping of a city reveals socioeconomic determinants of transmission within and between urban quarters

**DOI:** 10.1101/2020.04.03.023135

**Authors:** Adrian Egli, Nina Goldman, Nicola F. Müller, Myrta Brunner, Daniel Wüthrich, Sarah Tschudin-Sutter, Emma Hodcroft, Richard Neher, Claudia Saalfrank, James Hadfield, Trevor Bedford, Mohammedyaseen Syedbasha, Thomas Vogel, Noémie Augustin, Jan Bauer, Nadine Sailer, Nadezhda Amar-Sliwa, Daniela Lang, Helena M.B. Seth-Smith, Annette Blaich, Yvonne Hollenstein, Olivier Dubuis, Michael Nägele, Andreas Buser, Christian H. Nickel, Nicole Ritz, Andreas Zeller, Tanja Stadler, Manuel Battegay, Rita Schneider-Sliwa

## Abstract

With two-thirds of the global population projected to be living in urban areas by 2050, understanding the transmission patterns of viral pathogens within cities is crucial for effective prevention strategies. Here, in unprecedented spatial resolution, we analysed the socioeconomic determinants of influenza transmission in a European city. We combined geographical and epidemiological data with whole genome sequencing of influenza viruses at the scale of urban quarters and statistical blocks, the smallest geographic subdivisions within a city. We observed annually re-occurring geographic clusters of influenza incidences, mainly associated with net income, and independent of population density and living space. Vaccination against influenza was also mainly associated with household income and was linked to the likelihood of influenza-like illness within an urban quarter. Transmissions patterns within and between quarters were complex. High-resolution city-level epidemiological studies combined with social science surveys such as this will be essential for understanding seasonal and pandemic transmission chains and delivering tailored public health information and vaccination programs at the municipal level.

## Introduction

Transmission of influenza is influenced by extensive but poorly understood interactions between various viral, host and environmental factors^1-3^. Influenza may serve as a model for pandemic threats including the most recent COVID-19 pandemic. Whole viral genome sequencing has enabled reconstruction of phylogenetic relatedness at high resolution. Using these approaches, the interactions and dynamics of influenza transmission events have been described across a range of scales: globally^4-6^, across continents^1,7^, in university campuses^8^, or within households^9-12^. With two-thirds of the global population projected to be living in urban areas by 2050, understanding the transmission patterns of influenza within cities is crucial for effective prevention strategies and may help to prepare for pandemic threats. Previous work identified cities as containing critical chains of transmission outside of peak climatic conditions (Dalziel, B. D. et al Science 2019), but the resolution to look at these critical intra-city transmission chains in detail has until now been lacking. Very few studies have explored transmission events and dynamics of influenza viruses at the scale of a city^13-17^. Cities are heterogeneous with remarkably different neighbourhoods based on the socioeconomic position of the individuals living there. Consequently, a high spatial resolution of the urban space and built environment is crucial to fully understanding the impact of factors linked to health and disease^18,19^.

In this study we combine epidemiologic, geographical and demographical factors at the unprecedented resolution of urban quarters (i.e. neighbourhood/area) and statistical blocks (i.e. city block/street) levels, the smallest statistical enumeration areas within a city. Basel, Switzerland, serving as our model city, we explored the local patterns of influenza distribution and transmission from 2013 to 2018. Basel has urban quarters that differ substantially in socioeconomic indicators and housing structure. By using information at the level of statistical blocks, we were able to construct a detailed picture of influenza transmission within the city. We visualized kernel density estimates of influenza cases (reflecting the clustering of cases), a fundamental data smoothing method where inferences about the population are made, together with various population-based factors on maps with high resolution. Cases were corrected for population density and socioeconomic factors such as education levels, income and available living space were then analysed. Furthermore, we complemented the data with a detailed personal survey distributed to 30,000 households during the 2015/2016 flu season, to further explore urban quarter-specific aspects of self-reported influenza-like illness and association to socioeconomic factors. Finally, we described the details of influenza transmission during the 2016/2017 season using whole genome sequencing of all collected influenza viruses, covering more than 650 isolates, combining the data, resulting in an unprecedented spatiotemporal resolution. The details on the study design for this project have previously been published^20^.

## Results

### Reoccurring influenza patterns within a city

The City of Basel has 19 urban quarters and a stable mean population of 175,350 inhabitants during the 5-year study period (+/- 1,737 inhabitants) (**Figure S1A**). A total of 1,078 statistical blocks were identified within the city for the subsequent analysis. First, we collected epidemiologic and geographic information of 1,715 PCR-confirmed influenza cases over five consecutive influenza seasons from 2013/2014 to 2017/2018 and identified areas with a high burden of influenza infections (**Figure S1B-C**). Influenza viruses isolated in Switzerland were similar to viruses isolated across Europe in the study period (**Table S1**). On a weekly basis, we linked each individual case of PCR-confirmed influenza to the patient’s place of residence and anonymized the data at the scale of statistical blocks. We determined both the kernel density estimates for absolute influenza cases across all urban quarters and observed regional clusters of influenza cases within the city across all five seasons (**Figure 1A**) with similar spatial patterns for each individual season (**Figure S1D-H**).

**Figure 1:**
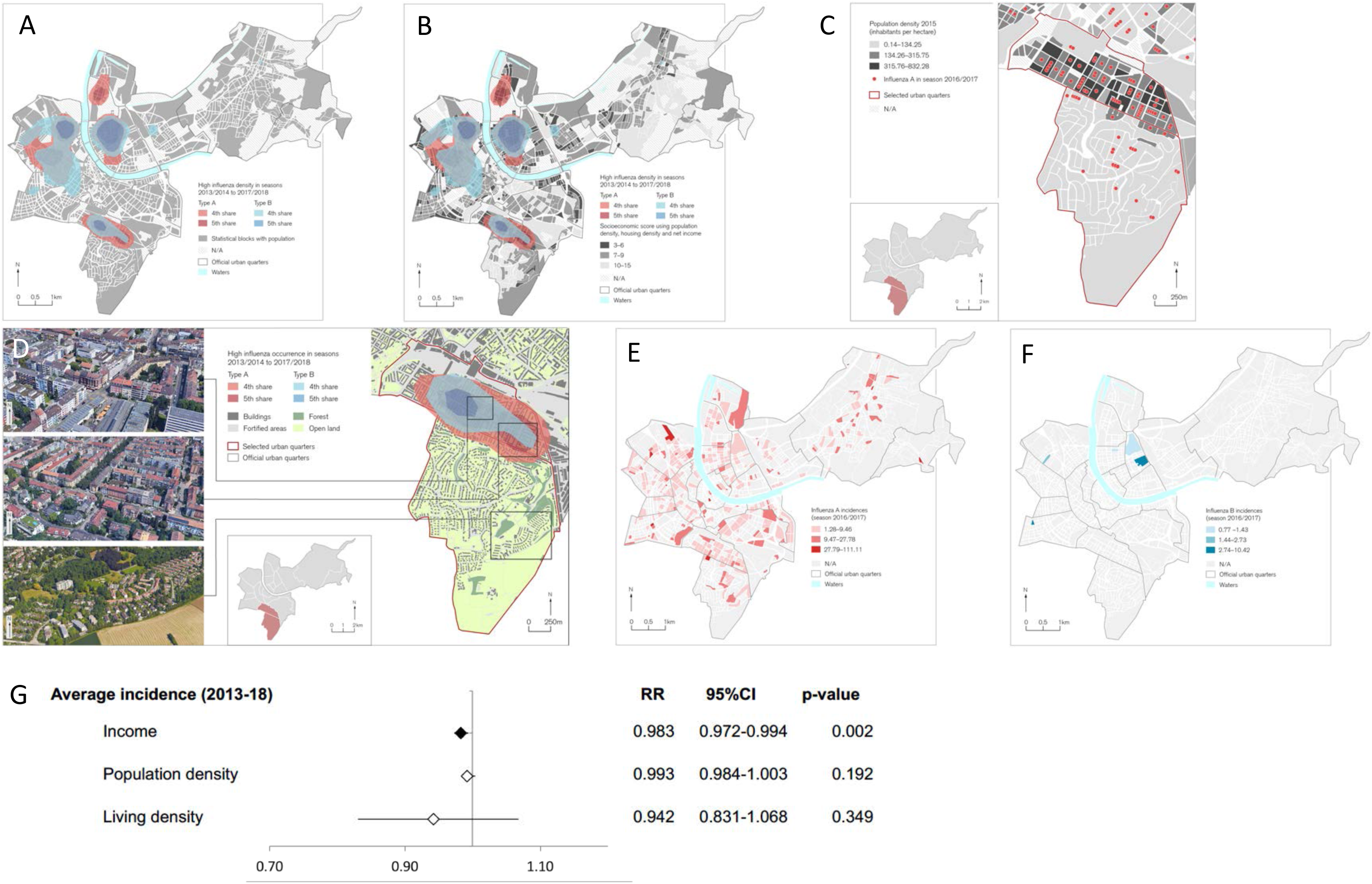
Influenza cases in Basel 2013-2018. The graph shows the distribution of PCR confirmed influenza cases in the City of Basel at the resolution of statistical enumeration blocks. **(A)** Geographical clustering of influenza infections. A kernel density estimation across all five influenza seasons from 2013/2014 to 2017/2018 was used to analyse the influenza case distribution across the city. The 4^th^ and 5^th^ share of the kernels were visualized. The red clouds show Influenza A cases and the blue clouds Influenza B cases. **(B)** As **A**, with underlying socioeconomic score for each housing block visualized, ranging from 3 to 15. The score consists of three individual factors: population density per ha, living space per person in m^2^ and net income in CHF. The natural break of the socioeconomic scores were used and visualized with three different shades of grey. Clusters of influenza correlate with lower socioeconomic scores. **(C)** Case load across representative examples of urban quarters (Bruderholz as less densely populated and Gundeldingen as more densely populated). Each red dot represents an Influenza A case. In order to not show the actual address of a patient, we set the dots in the middle of the statistical block. **(D)** We combined Google Earth satellite images with maps showing the kernel density estimates of influenza cases to indicate building density. Red clouds show Influenza A cases and blue Influenza B cases, correlating with building density, reflecting population density. **(E)** Influenza A virus incidence rates of season 2016/2017 per 1000 inhabitants of each statistical housing block. **(F)** Influenza B virus incidence rates of season 2016/2017 per 1000 inhabitants of each statistical housing block. **(G)** Multivariable Poisson regression analysis to explore the association of influenza virus incidence (per 1000 inhabitants corrected for the population of each housing block) and each socioeconomic factor. Average influenza incidences for the seasons 2013/2014 to 2017/2018 are shown. Net income relative risk per 1 CHF; populations density per person per ha; living density per person in m^2^.

We further explored the association between influenza occurrence and socioeconomic factors as well as the built city environment. We linked socioeconomic scores to each statistical block reflecting (i) the population density (inhabitants per hectare (ha)), (ii) the living space (per capita in m^2^), and (iii) the net income (median in CHF). Each of these three key factors was translated to “socioeconomic points” ranging from one to five, which were added to generate a total socioeconomic score for each housing block: a score of three reflecting the lowest and a score of fifteen the highest possible socioeconomic value^20^ (**Figure S2A-D**). The median socioeconomic scores of urban quarters ranged from three to ten. We observed that city areas with a high socioeconomic score had fewer influenza cases in comparison to areas with lower socioeconomic scores (**Figure 1B**). **Figure 1C** provides a representative example of the effect of population density during the 2016/2017 influenza season comparing the urban quarters Gundeldingen (GU, high population density with high influenza burden) with Bruderholz (BR, low population density with low influenza burden). These two urban quarters are located next to each other but are notably different. The urban quarter with small detached residential buildings (single family homes) showed a mean socioeconomic score of 9.5 (+/- 1.71 standard deviation) while the other urban quarter with large (multifamily) residential buildings shows a mean socioeconomic score of 5.6 (+/- 1.87 standard deviation). The housing structures of particular urban quarters may serve as a surrogate marker for the population density and available living space in a particular area and thereby also be an indicator of influenza burden (**Figure 1D**).

To account for this potential ecologic fallacy, we corrected the influenza incidence rates per 1,000 inhabitants for each statistical block. We still observed similar dense influenza case patterns at the level of statistical blocks across urban quarters during the examined influenza seasons (**Figure 1E-F, S3A-H, Supplementary file 1**). Besides visualizations of influenza case distributions, we further explored associations between socioeconomic factors and the influenza incidence (corrected for the population density within each statistical block). The analysis included all five seasons and data from all 1,484 statistical blocks for every year. We used a multivariable Poisson regression and observed that net income of the statistical block (per 1,000 CHF) was the strongest predictor for influenza incidence at the level of statistical blocks, independent of population density and living space in most influenza seasons (**Figure 1G, S4**).

### Socioeconomic factors determinate herd immunity

To further explore the interrelationship between socioeconomic factors and rates of influenza-like illness, we conducted a detailed survey across ten urban quarters for the 2015/2016 season (**Figure S1A**). The survey included 54 questions, addressing (i) influenza-like illness and vaccination, (ii) aspects of the urban environment, (iii) access to health care information, (iv) health related data, and (v) the place of residence at the level of statistical enumeration blocks. The English version of the survey is attached as supplementary file (**Supplementary File 2**).

The share of self-reported influenza-like illness cases fulfilling all three defining criteria for each of the selected urban quarters ranged from 3.4% to 7.0% (n=358; median 4.5%) during the influenza season 2015/2016 (**Figure 2A; S5A**). The number of influenza-like illness and PCR-confirmed cases (71 influenza A and 44 influenza B) did not correlate in that particular season across the explored statistical blocks responding to the survey (r=0.0144, p=0.7411; **Figure S5B**). However, the reported frequency of influenza-like illness corresponds to previously reported attack rates of three to five percent^21-23^. Other respiratory viruses likely contribute to cases matching the non-specific influenza-like illness definition. In our analysis, 20-25% of the influenza PCRs performed on patients presenting an influenza-like illness are subsequently confirmed by PCR-based diagnostics as indicated for the 2017/2018 influenza season (**Figure S5C**).

**Figure 2:**
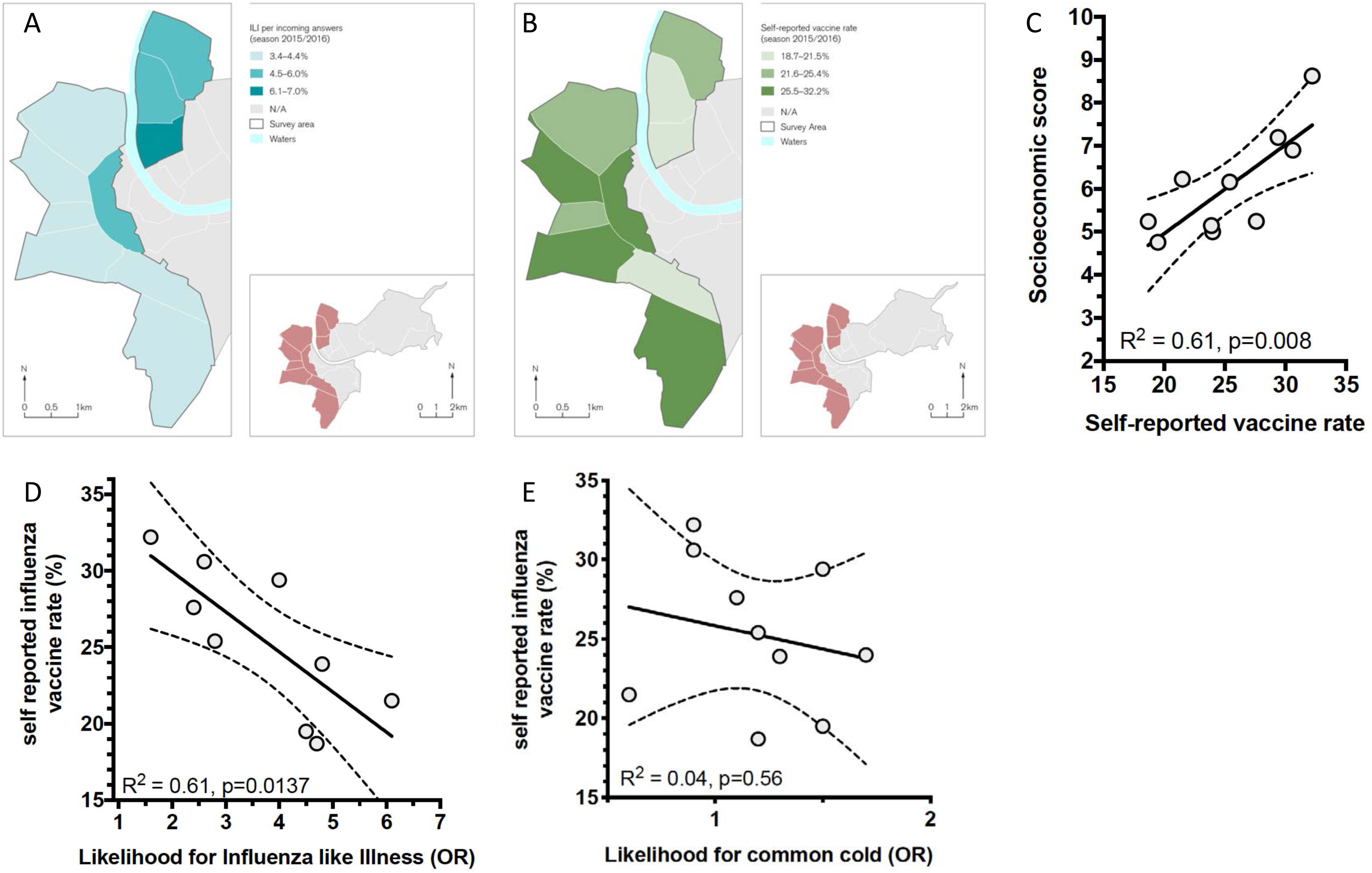
Survey on self-reported Influenza-like illness and vaccination over season 2015/2016. **(A)** Distribution of self-reported influenza-like illness cases according to the WHO definition^41^ over the 10 selected urban quarters. The number of Influenza-like illness cases per incoming answers is expressed as percentage value for each quarter. The blue colour range is according to natural breaks and ranges from 3.4% to 7%. **(B)** Self-reported vaccine rates (on a per person basis / all received surveys for the specific urban quarter). Natural breaks between urban quarters are shown in different shades of green, ranging from 18.7% to 32.2%. **(C)** Comparison of socioeconomic score with self-reported vaccine rates using logistic regression shows a correlation at the levels of urban quarter in the aggregate. **(D)** Self-reported vaccine rates against influenza and the likelihood for ILI, using logistic regression, shows an inverse correlation. **(E)** Self-reported vaccine rates against influenza and the likelihood of common cold, using logistic regression, shows no correlation.

Next, we performed regression models with self-reported influenza-like illness as a binary endpoint for each individual. The factors associated with increased relative risks for self-reported influenza-like illness in stepwise forward and backward selection multivariable analysis were: ≥3 people per household and daily use of public transport. Of note, a total of 22.8% (1859 of 8149) were from households with ≥3 people. Of which 81.3% (1511/1859) were couples or single parents with children and 8.8% (720/8149) had children under the age of 7 years. The factors associated with decreased relative risks were: vaccination against influenza, age more than 65 years, and daily physical activity (**Table 1; Table S2**). The most protective variable against influenza-like illness was vaccination. In a multilevel multivariable model, vaccination remained the most important factor even when correcting for the urban quarter an individual lived in (RR 0.4, 95% CI 0.24 – 0.67). Of note, people with and without self-reported influenza vaccination showed similar odds ratios for symptoms of common cold (respiratory symptoms not fulfilling the influenza-like illness case definition, such as running nose and sore throat; OR 0.98, 95% CI 0.78 - 1.24).

**Table 1.**
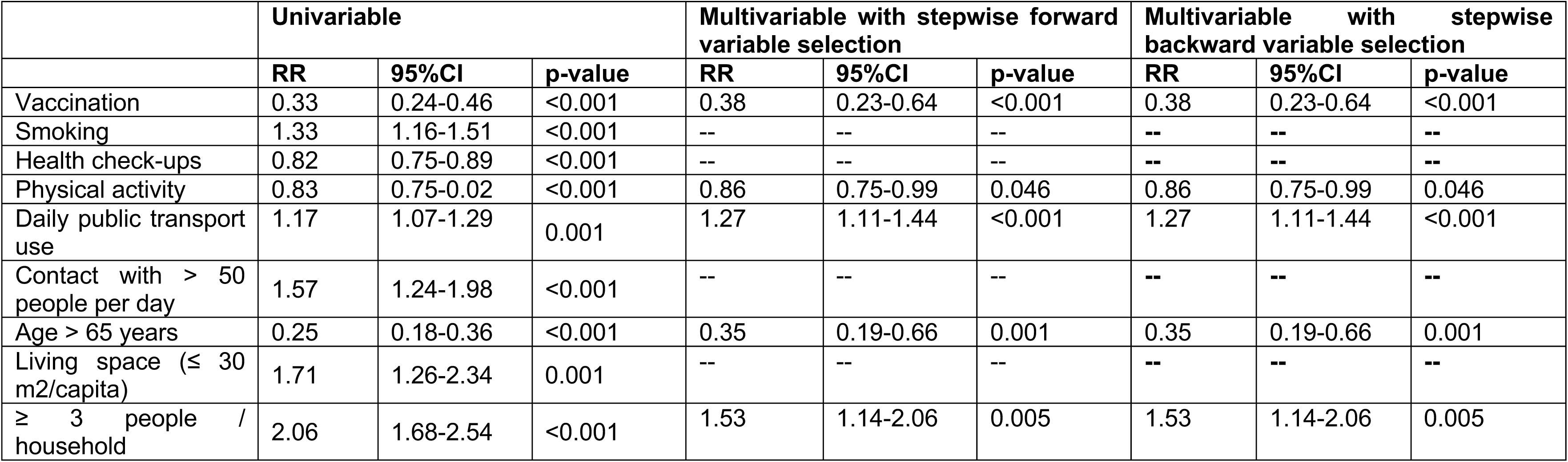
Factors associated with self-reported influenza-like illness. Forward and backward selection multivariable regression models for factors, which are associated with influenza-like illness. All variables with univariable associations to influenza-like illness are shown in **Table S1**. RR, relative risks.

The self-reported influenza vaccination rates were heterogeneously distributed at the level of urban quarters (**Figure 2B; S5D**). The median vaccine rate was 25.8% (IQR 18.7% - 32.2%) throughout the city, which is in the range of the previously reported averages in Switzerland ^24^. We performed regression models with self-reported vaccination against influenza as binary endpoint for each individual. The factors consistently associated with an increased likelihood for self-reported influenza vaccination were people belonging to a risk group of chronic disease and health care workers. Low household income (below 6000 CHF per month) was associated with a significant lower likelihood of self-reported vaccination (**Table 2; Table S3**). When adjusting the multivariable model to specific urban quarters, low income was no longer correlated with vaccination (p=n.s.), which confirms the previously mentioned profound difference of income between urban quarters (**Figure S2C**).

**Table 2.**
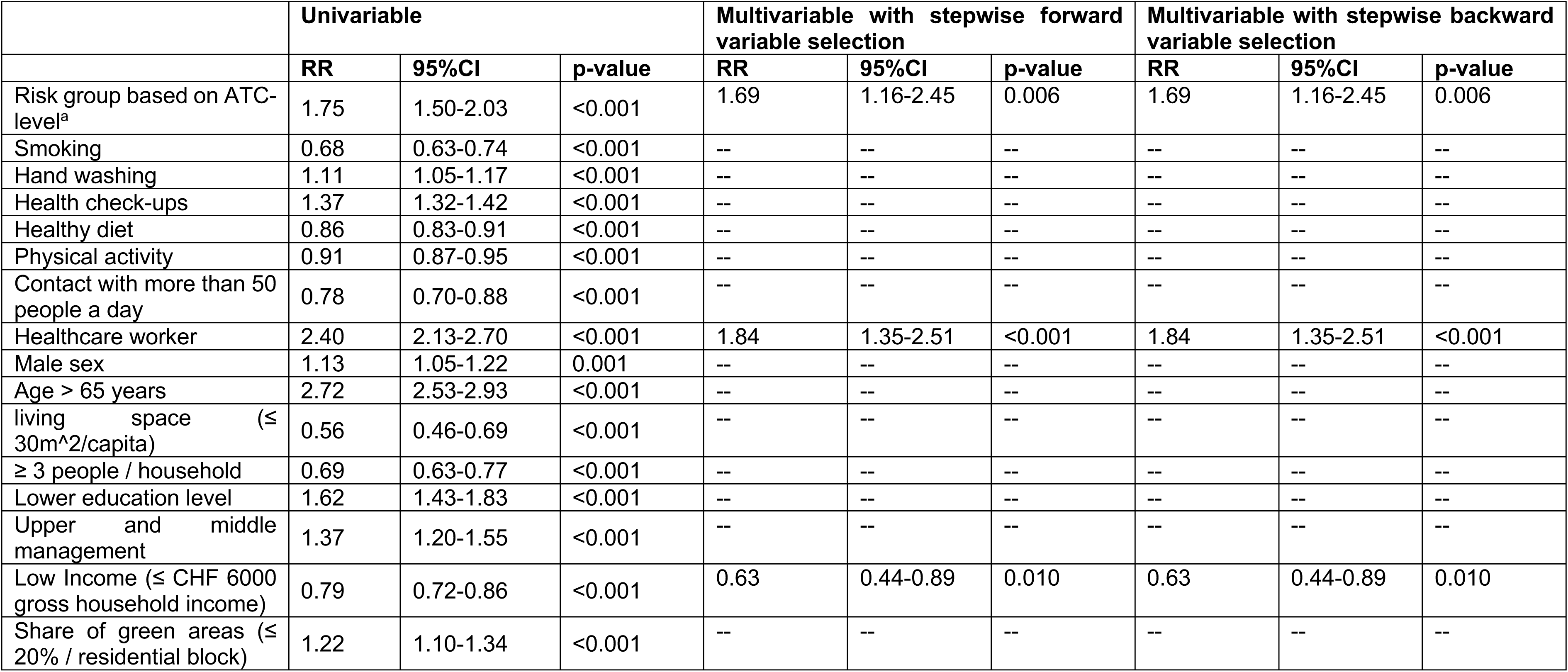
Factors associated with self-reported vaccination against influenza. Forward and backward selection multivariable regression models for factors which are associated with vaccination against influenza. Data on univariate associations with vaccination can be found in **Table S2**. RR, relative risks. ^a^, Anatomical Therapeutic Chemical (ATC) according to WHO definition.

|  | Univariable |  |  | Multivariable with stepwise forward variable selection |  |  | Multivariable with stepwise backward variable selection |  |  |
| --- | --- | --- | --- | --- | --- | --- | --- | --- | --- |
|  | RR | 95%CI | p-value | RR | 95%CI | p-value | RR | 95%CI | p-value |
| Risk group based on ATC-level<sup>a</sup> | 1.75 | 1.50-2.03 | <0.001 | 1.69 | 1.16-2.45 | 0.006 | 1.69 | 1.16-2.45 | 0.006 |
| Smoking | 0.68 | 0.63-0.74 | <0.001 | -- | -- | -- | -- | -- | -- |
| Hand washing | 1.11 | 1.05-1.17 | <0.001 | -- | -- | -- | -- | -- | -- |
| Health check-ups | 1.37 | 1.32-1.42 | <0.001 | -- | -- | -- | -- | -- | -- |
| Healthy diet | 0.86 | 0.83-0.91 | <0.001 | -- | -- | -- | -- | -- | -- |
| Physical activity | 0.91 | 0.87-0.95 | <0.001 | -- | -- | -- | -- | -- | -- |
| Contact with more than 50 people a day | 0.78 | 0.70-0.88 | <0.001 | -- | -- | -- | -- | -- | -- |
| Healthcare worker | 2.40 | 2.13-2.70 | <0.001 | 1.84 | 1.35-2.51 | <0.001 | 1.84 | 1.35-2.51 | <0.001 |
| Male sex | 1.13 | 1.05-1.22 | 0.001 | -- | -- | -- | -- | -- | -- |
| Age > 65 years | 2.72 | 2.53-2.93 | <0.001 | -- | -- | -- | -- | -- | -- |
| living space (≤ 30m<sup>2</sup>/capita) | 0.56 | 0.46-0.69 | <0.001 | -- | -- | -- | -- | -- | -- |
| ≥ 3 people / household | 0.69 | 0.63-0.77 | <0.001 | -- | -- | -- | -- | -- | -- |
| Lower education level | 1.62 | 1.43-1.83 | <0.001 | -- | -- | -- | -- | -- | -- |
| Upper and middle management | 1.37 | 1.20-1.55 | <0.001 | -- | -- | -- | -- | -- | -- |
| Low Income (≤ CHF 6000 gross household income) | 0.79 | 0.72-0.86 | <0.001 | 0.63 | 0.44-0.89 | 0.010 | 0.63 | 0.44-0.89 | 0.010 |
| Share of green areas (≤ 20% / residential block) | 1.22 | 1.10-1.34 | <0.001 | -- | -- | -- | -- | -- | -- |

The survey results confirm that vaccination against influenza has a protective effect against influenza-like illness. The odds of acquiring influenza-like illness during the 2015/2016 season were 3-times less in the group of vaccinated people (0.33 OR, 95% 0.24-0.46) in comparison to non-vaccinated people. This correlates to reported vaccine effectiveness rates of 44% in children and 78% in adults against influenza for the 2015/2016 season^25,26^ (**Table S4**). The median self-reported vaccination rates between urban quarters showed a direct correlation with the socioeconomic score of the respective urban quarter (R^2^=0.607, p=0.0079; **Figure 2C**) – the higher the socioeconomic score, the higher the vaccine rate. Income showed the highest correlation with vaccination rates, followed by living space and population density, respectively (R^2^ 0.622, p=0.0067; R^2^ 0.618, p=0.007; R^2^ 0.532, p=0.0167). Self-reported vaccination rates may serve as a surrogate marker for herd immunity^27,28^. Similarly, at the level of an urban quarter, herd immunity strongly affected the risk of acquiring influenza-like illness during the 2015/2016 season. Urban quarters with a high self-reported vaccination rate showed a significantly lower likelihood of influenza-like illness of the surveyed population compared to urban quarters with a low vaccine rate (R^2^=0.61, p=0.01; **Figure 2D**). However, no protective effect against common cold was observed (**Figure 2E**, p=0.56). Therefore, unvaccinated people living in the Matthaeus quarter (MA) show an 8-times higher probability of contracting an influenza-like illness than those vaccinated while in the Bruderholz quarter (BR) unvaccinated people only show a 1.2-fold higher risk. We again observed the association of self-reported influenza-like illness and vaccination rates (in % of returned survey) at the level of individual statistical blocks (r=-0.1182, p=0.007). However, in years with a low vaccine effectiveness (**Table S4**) this association may be weaker.

### Humoral immunity of healthy donors is variable across urban quarters

In order to monitor antibody titres over time in a healthy population across the city, we recruited 214 healthy blood donors living in Basel before the 2016/2017 influenza season. We determined their hemagglutination inhibition titres against the circulating virus, H3N2 (Influenza A/Hong Kong/4801/2014). We also quantified antibody titres against all other vaccine strains (Influenza A/California/7/2009 H1N1 pdm09; Influenza B/Brisbane/60/2008; and Influenza B/Phuket/3073/2013). Previous antigen exposure to other strains may affect the response rates in the general population (^29,30^**; Table S1**). Before the 2016/2017 influenza season, we observed that across all urban quarters a median of 21% (IQR 17-28.5%) had seroprotective antibody levels (defined as hemagglutination inhibition titres equal or more than 1:40^31^) (**Figure 3A**). Again, urban quarters with lower socioeconomic scores also showed low seroprotection rates (e.g. Matthaeus, Breite, Kleinhueningen and Klybeck) (**Figure S6A**). Urban quarters with higher socioeconomic scores showed a median seroprotection rate of 26.1%, whereas those with lower socioeconomic scores showed a median seroprotection rate of 14.6% (p=0.05). Blood donors with influenza vaccination showed significant higher H3N2 specific HI titers in comparison to people who were not vaccinated (p<0.0001; **Figure S6B**). Similar to the survey, in this cohort net income was associated with the vaccination status. Blood donors who were influenza vaccinated had a median higher net income per statistical block when compared to non-vaccinated blood donors (median CHF 54,144 vs. CHF 48,898, p=0.047; **Figure S6C**), whereas population density did not differ (p=0.39).

**Figure 3:**
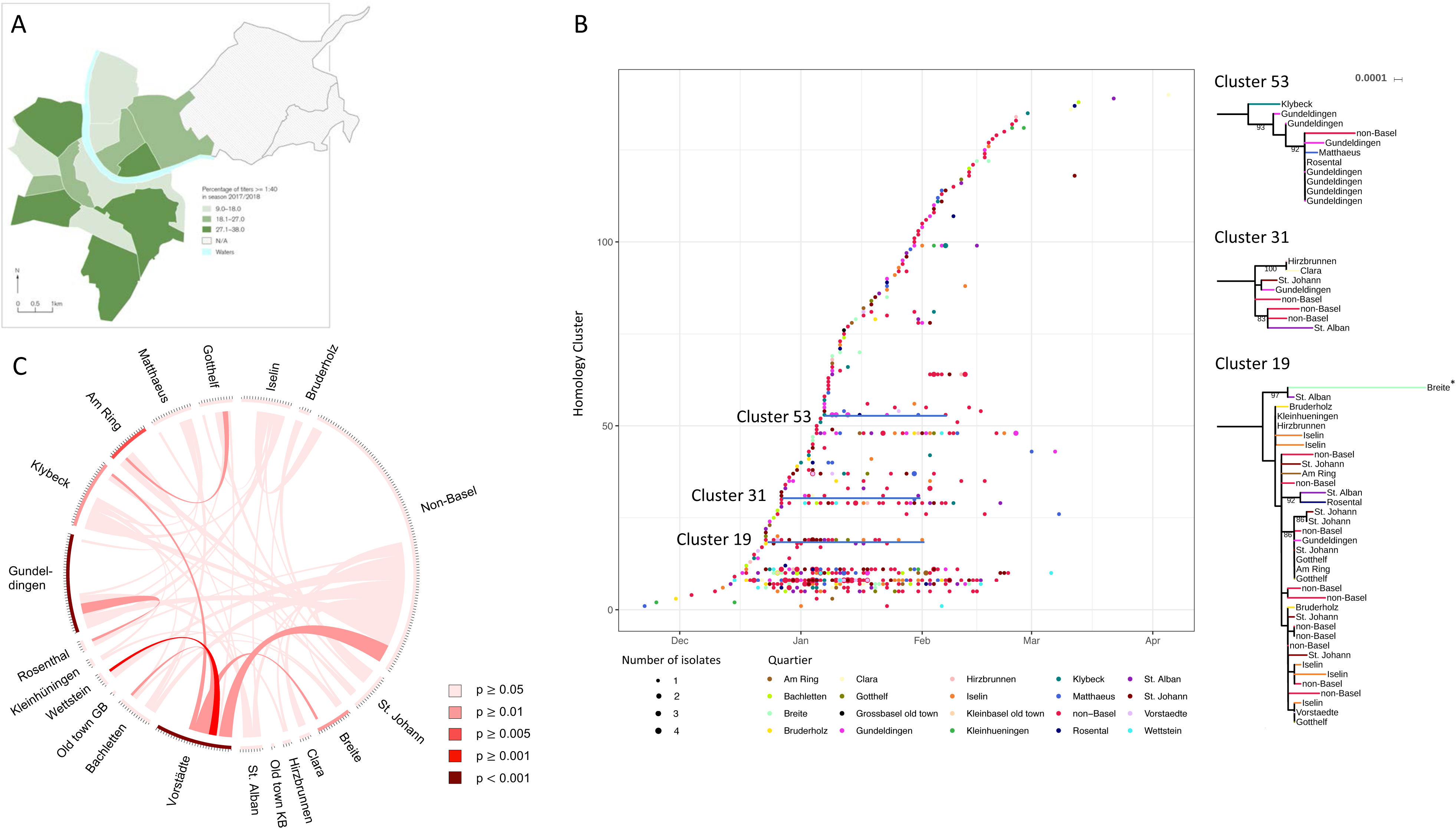
Influenza antibody titres and whole genome sequencing of influenza viruses to explore transmission between urban quarter levels. **(A)** Pre-seasonal antibody titres against H3N2 from healthy blood donors in the 2016/2017 influenza season measured using hemagglutination inhibition assay. Distribution of seroprotection (defined as titre >1:40) are visualized as percentages using natural breaks. **(B)** Cluster analysis of each influenza case genome during the 2016/2017 season. Each dot represents a sequenced influenza virus isolate. Colour reflects the host address by specific urban quarter. Each horizontal line corresponds to a local transmission cluster of influenza viruses. Three representative clusters 19, 31 and 53 are detailed on the right using maximum likelihood trees indicating the complexity of transmission clusters across time and urban quarters. **(C)** Exchange of influenza viruses throughout the city. Red lines connecting two quarters indicates isolates that are identical within the same transmission cluster, with the line width and colour intensity indicating the number of isolates in that cluster shared by the connected quarters. Therefore, they indicate the level of transmissions between two quarters. When correcting for multiple comparison, only the transmission within the same urban quarters remained statistically significant (p<0.05), which is indicated as a coloured bar at the basis of the circle. The darker the colour the more significant the association.

### Geographic patterns of influenza transmission within a city

Next, we studied the impact of geographic and socioeconomic factors on influenza transmission using molecular epidemiological techniques. During the 2016/2017 season, we screened patients presenting an influenza-like illness using an influenza-specific PCR. We included 663 influenza virus positive samples recruited from 12 different study sites across the city (^32^; **Figure S7A**). The largest cohort within a single city, to the best of our knowledge. First, we determined antibody titres against the circulating H3N2 virus and other vaccine viruses in the patients, who having presented within days of feeling sick still had high viral loads meaning that antibody levels could be assumed to be close to baseline. Patients with PCR-confirmed influenza showed significant lower antibody titres against influenza A H3N2 in comparison to patients without influenza, highlighting the importance of protective antibodies (**Figure S7B** and **S7C**). Of note, antibody titres against viruses not circulating (e.g. H1N1) where not significantly different between PCR positive and negative patients.

A total of 663/858 viral genomes passed our duplicate and quality control filter (methods and^33^) and were included in subsequent analyses. 427/663 (64.4%) of isolates were from people living in Basel and the remaining 236/633 (37.3%) isolates were people living outside of Basel, mainly in surrounding villages within a 20km range but occasionally (n=6/633) from further away (e.g. Columbia, Turkey or Italy).

First, we compared the hemagglutinin (HA) gene sequence from our strains to a global sample of ∼1400 influenza virus HA genes from the GISAID data repository of the 2016/2017 seasons. Using the integrated visualization of a phylogenetic tree and a map implemented in Nextstrain, we investigated the geographic distribution of influenza virus diversity at the levels of cities, urban quarters, and statistical blocks (https://nextstrain-dev.herokuapp.com/community/appliedmicrobiologyresearch/Influenza-2016-2017/h3n2/ha; links to visualization of all viral segments are provided in Table S5). This visualization can be interactively explored using the narrative functions of Nextstrain. The diversity of European influenza isolates was fully represented by the isolates collected from patients living in Basel and the surrounding area. In addition, also at the level of urban quarters and statistical blocks, we observed the same high intermixture and diversity. Although the socioeconomic scores differ between urban quarters and statistical blocks, we did not observe an association with the phylogenetic tree structure (https://nextstrain-dev.herokuapp.com/community/narratives/appliedmicrobiologyresearch/Influenza-2016-2017/baselFlu).

Next, we further explored viral transmission clusters in more detail. Based on previously published mutation rates^34,35^, we assumed that influenza viruses accumulate an average of 10 single nucleotide polymorphisms during a single influenza season. Viral strains with a genetic relatedness within this range were assigned as being in the same transmission cluster. A sequence, which is more than 10 single nucleotide polymorphisms different from any other sequence was ignored, so that a transmission cluster contains at least 2 viral strains. The 54 transmission clusters identified (**Figure 3B**) incorporated 547/663 influenza strains, each comprising a minimum of two isolates, a median of three isolates, ranging from 2 to 111 within and outside of the city (**Figure S7D**). Only 116 (17.5%) cases could not be linked to another individual. Most of the transmission clusters contained samples from more than one urban quarter, and only a few clusters were predominantly located within a single urban quarter (see clusters 19, 31 and 53, **Figure 3B;** https://nextstrain-dev.herokuapp.com/community/narratives/appliedmicrobiologyresearch/Influenza-2016-2017/baselFluClusters). Removing isolates from outside the city and focusing only on transmission events within city, we observed that 368/427 (76%) influenza strains belong to 43 of the overall 54 clusters across this single influenza season.

Next, we connected all identical influenza isolates between different urban quarters. Using permutation tests, we observed a generally high exchange rate and complex transmission dynamic between the different urban quarters. Some urban quarters showed significant connections to other urban quarters. For example, isolates from Vorstaedte (VO) and Wettstein (WE) were more identical in comparison to other urban quarters (p<0.005, **Figure 3C**). Transmission events within the same urban quarter were explored. Interestingly, two urban quarters – Gundeldingen (GU) and Vorstaedte (VO) – showed influenza isolates that were significantly more related to other isolates from within the same urban quarter than to isolates from other quarters or outside of Basel (p<0.001, **Figure 3C**). These two urban quarters show a low socioeconomic score and lower pre-seasonal seroprotection rate. Phylogenetic cluster size did not correlate with any socioeconomic factors (p=n.s.).

## Conclusions

Each year influenza infects millions of people around the globe^36,37^. Historically, human pandemics have had devastating outcomes, and preventive measures should have high priority. Our results reflect an unprecedented large and dense datasets of PCR-confirmed influenza cases, the largest city-wide survey on influenza-like illness performed to date, and influenza genome sequences, all mapped to the statistical block level. We observed annually re-occurring geographic clusters of influenza cases and incidences. These influenza hot spots were mainly associated with net income, and independent of population density and living space. Vaccination against influenza was heterogeneously distributed between urban quarters and dependent mainly on household income. The rate of vaccination was linked to the likelihood of contracting an influenza-like illness within a specific urban quarter. Finally, transmission events between quarters were highly complex, but for two urban quarters particularly high levels of transmission events within the same urban quarter could be observed.

Our study has certain limitations. First, in the survey, a more research-interested population may have replied to the questionnaire. In particular, 37.9% of males replied to the survey, whereas in a census 48.2% were males. However, the overall reported vaccination are in line with regular interviews conducted by the Federal Office of Public Health of Switzerland, which report vaccination rates of >30% in people with 64 years of age and older^24^. Also, the self-reported influenza-like illness rates corresponds to estimated and published attack rates^38,39^. For influenza diagnostic using PCR confirmation, we cannot exclude some recruitment bias, that particular people with a specific demographic background are more likely to be included at a tertiary healthcare centre. In order to compensate for such a bias, we also included data from other study sites including a children’s hospital and family doctors and a private diagnostic laboratory which also covers family doctors and other non-University hospitals in the region (**Figure S7A**). Further, ecologic fallacy may link certain socioeconomic factors, therefore we adjusted our analysis for population density. However, when correcting for population densities, statistical associations remained stable. Although, the influenza sampling for the sequencing part of the study was reasonably thorough, certainly not all influenza cases could be captured as many patients with influenza will not seek healthcare. Thus, some transmission links will always remain unknown. For more detailed analyses, future studies would need to include hundreds of samples with high resolution epidemiological data across multiple influenza seasons to study the changing patterns of transmission over multiple years. In addition, the transmission events across urban quarters are highly complex, driven by mobility of people living within a city and commuters, place of working, and changing antibody titres of exposed patients in each year. Although we have asked where a person works, we were not able to collect sufficient data for this question – only 24.6% provided details and people working in the same statistical block were very rare (<5%).

Our findings provide important insights demonstrating influenza transmission patterns in a city serving as a model for studying the dynamics of seasonal flu transmission and evolution within a city. These results should be repeated in cities of different sizes and complexities around the world to allow public health services to be tailored most effectively. It will be interesting to see whether further factors concerning influenza transmission can be identified in other cities as well as their role in public health measures. Importantly, since vaccination rates were strongly dependent on income and linked to influenza incidences, providing better access to vaccination for low income households would likely have a substantial impact on influenza transmission. The knowledge gained with our study can help to tailor public health measures such as urban vaccination programs to urban influenza hotspots and not just to selected target groups (e.g. high-risk populations with chronic illness). National vaccination recommendations often only include a selected population, e.g. health care workers, patients with comorbidities or particular age groups. Our results suggest that such strategies may entirely miss the populations where most transmission occurs such as elderly, chronically ill, and children. Finally, this large-scale interdisciplinary study may serve as a blueprint for investigating seasonal and pandemic viruses on the smallest scale within a given geographical context such as a city. It may form the basis to develop more effective and targeted counter-measures at the most relevant public health levels e.g. at the urban quarter, or even smaller urban subdivisions and social milieus reflected therein. This would allow us to address more and account for a greater variety of population segments and help to identify potential drivers of transmission.

## Methods and Material

### Data and sample collection

The overall study design has been previously published ^20^. Briefly, the study had retrospective and prospective parts. The retrospective study part consisted of an analysis of the numbers of PCR-confirmed influenza cases over the course of five years from 2013 to 2018. The prospective study part consisted of (i) a household survey focusing on influenza-like illness and vaccination against influenza during the 2015/2016 season; (ii) an analysis of influenza viruses collected during the 2016/2017 season using whole genome sequencing of viral genomic sequences to determine genetic relatedness, clusters and putative transmission events; and (iii) measurement of influenza-specific antibody titres against all vaccinated and circulated strains during the 2016/2017 season from healthy individuals, allowing us to monitor herd immunity across urban quarters. Particular patient groups may more likely present at a hospital and receive PCR-based diagnostics – to minimize this potential bias, we also integrated data from a large private diagnostic laboratory, which includes mainly non-hospitalized patients. Study teams at the University Hospital Basel and the University Children Hospital collected data and recruited patients for this study. In addition, a network of 24 paediatricians and family doctors also helped in recruiting patients (see acknowledgments). Viollier, a private laboratory, providing its services to a large number of private practices within the City of Basel, also provided samples and data.

#### Ethics and dissemination

The study is registered (clinicaltrials.gov; NCT03010007 on 22^nd^ December 2016) and approved by the regional ethics committee as an observational study (EKNZ project ID 2015–363 and 2016-01735).

#### PCR-confirmed cases

Influenza specific PCRs were performed on nasopharyngeal swabs as part of the routine diagnostic workup. For (semiquantitative) PCR detection all laboratories used either the Xpert Flu/RSV or the Xpress Flu/RSV (Axonlab, Switzerland). For each PCR-confirmed case the exact place of residency was transposed to statistical blocks in order to anonymize the data using a geoinformation system (ArcGIS by ESRI, Switzerland) and to visualize the findings on maps. A statistical block is usually bordered from all sides by roads. In individual cases, the boundary is by zone plan categories (e.g. railway areas, forest, green zone, agricultural zone, etc. ^40^). For each statistical block demographic data was obtained from the Statistics Office of the Canton Basel-City: net income in CHF, population per ha, and living space in m^2^ per capita. These socioeconomic factors contributed to an overall socioeconomic score for every statistical block ranging from minimum three to maximum fifteen.

#### Survey on ILI and vaccination

We selected urban quarters for the survey a priori, based on identified diversity in their socioeconomic structure (**Figure S2A-D**). We translated the questionnaire into the six most common spoken languages in the selected urban quarters in order to reach a maximum population. The survey was distributed within one week after the end of the influenza season. Of the 30,000 questionnaires distributed in 10 urban quarters, 8,149 (27.1%) were returned and fulfilled the quality criteria for subsequent analysis. Returned questionnaires were quality controlled and the data was entered into a database. Questionnaires were excluded when no information of influenza-like illness or vaccination status was reported. Influenza-like illness was defined according by the WHO as a combination of self-reported fever, coughing, and illness of less than ten days ^41,42^. The questionnaires allowed the option of self-identifying one’s place of residence within a statistical block within the urban quarter.

Relative risks for influenza-like illness and influenza vaccination were estimated by uni- and multivariable Poisson regression with robust error variance. To deal with possible confounding, all variables found to differ significantly in univariable analyses between participants with and without ILI and with and without influenza vaccination, respectively, were included in the multivariable generalized linear models. A Bonferroni-adjusted p-value threshold was applied to select variables for inclusion into the multivariable models. Poisson regression models using stepwise forward and backward selection were applied to identify variables independently associated with both primary outcomes.

To account for socioeconomic differences related to each urban quarter and potentially influencing both the risk for influenza-like illness acquisition and vaccination uptake, analyses regarding individual risk factors were complemented by multivariable, multilevel mixed-effects generalized linear models. The Pearson and deviance goodness-of-fit tests were performed to assess the fit of the data to a Poisson distribution in the final regression models. Analyses were performed using Stata statistical software, version 15.1 (Stata Corp, College Station, Texas, USA).

#### Sample collection

At 12 study sites we recruited paediatric and adult patients fulfilling the following criteria: cough, fever, and sudden disease onset – these patients with influenza-like illness were further evaluated using the previously described influenza specific PCR.

### Whole genome sequencing and data analysis

Details regarding the sequencing have been previously described^33^. Briefly, samples from PCR-confirmed cases had the whole genome amplified using PCR. From the resulting amplicons we created libraries using the Nextera XT protocol and sequenced them on a MiSeq platform (Illumina, San Diego, California) with 300bp paired end reads at 48-plex.

#### Quality control and genome assembly

We collected all data in the 2016/17 influenza season as previously described^20^. In total, we sequenced 857 samples. All genomes are available at NCBI with accession numbers MN299375 - MN304713. A full list of all used influenza genome for this publication (including GISAID strains) is available at: https://github.com/appliedmicrobiologyresearch/Influenza-2016-2017/blob/master/data_information/accession_numbers.tsv. If a patient was sampled more than once, we only included the first isolate, resulting in 755 remaining samples. If not otherwise indicated default settings of the bioinformatic tools were used. Raw Illumina reads were trimmed with Trimmomatic 0.36^43^. Alignment of paired-end reads used bowtie 2.2.3^44^ with strain A/New York/18/2014 as a reference (Fludb.org segment accession numbers: PB2, KT837257; NP, KT837224; PA, KT837236; NA, KT837237; PB1, KT837241; M1/M2, KT837260; HA, KT837206; NS1/2, KT837196). The aligned reads were sorted by using samtools 1.2^45^. Variants were called and filtered by using LoFreq 2.1.2^46^. Variant calling was done for sites with a coverage of at least 100. Sites with a coverage of less than 100 were assumed to be unknown and were denoted as N, that is any possible nucleotide. Sequences that showed a read depth of 100 for at least 80% of the positions in at least four segments were used for the analysis. Non double infections with two strains were noted. Using these parameters, we continued our analysis using 663 samples. The consensus sequences from these strains were deposited in GenBank (numbers will be available upon acceptance of the manuscript). We aligned the consensus sequences using muscle v3.8.31^47^ and used the concatenated alignment of all segments to calculated a maximum likelihood tree^8,48^ using RaxML (^49^; version 8.2.8, options: -x 1522 -f a -m GTRGAMMA -p 1522 - # 100) to depict the relationship between the isolates.

#### Identification of transmission clusters

In order to trace local influenza virus lineages, we grouped the different influenza sequences into clusters using a maximum distance of 10 SNPs using an in-house python script (https://github.com/appliedmicrobiologyresearch/Influenza-2016-2017). In a recent publication the average evolutionary rate for influenza was determined to be 2.5×10^−3^ nucleotide substitutions per site per year^50^. With a genomes size of 13,588 bp this equates to 34 mutations per year. Therefore, 10 SNPs can be estimated to corresponds to 3.5 months, the length of a typical influenza season in Basel.

#### Transmission within urban quarters

In order to determine direct transmission between patients, we use the same in-house python script and clustered the influenza strains that have no detected variants in-between. With this we found that 139 of 663 isolates (all full genomes) are connected in 52 clusters in which the isolates are identical (0 SNPs). To identify quartiers that show increase transmission within or to other quartiers, we performed a permutation test (10,000 repetitions) in which the connections between quarters are compared to samples to which the quartiers are randomly assigned using an in-house python script (https://github.com/appliedmicrobiologyresearch/Influenza-2016-2017). The distribution of clusters over time and in the different quarters were visualized using ggplot2^51^. The phylogenetic trees were visualized using iTOL^52^. The transmission of isolates between different quarters was visualized using Circos^53^.

#### Visualisation in Nextstrain

The visualization at https://nextstrain-dev.herokuapp.com/community/appliedmicrobiologyresearch/Influenza-2016-2017/h3n2/ha and https://nextstrain-dev.herokuapp.com/community/narratives/appliedmicrobiologyresearch/Influenza-2016-2017/baselFlu was produced using the Nextstrain toolchain^54^. Specifically, influenza virus A/H3N2 sequences were downloaded from GISAID^55^, filtered down to ∼1400 sequences from 2016-07-01 to 2017-06-30, augmented with influenza reference viruses, and combined with sequence data from this study. The resulting sequences were then aligned using mafft^56^, a phylogenetic tree was built using IQ-tree^57^, which was subsequently turned into a time scaled tree using treetime^58^ in analogy to the analysis workflow used for the weekly updated influenza phylogenies at nextstrain.org/flu. The visualization is implemented through the Nextstrain community feature.

The analysis was repeated for each segment of the influenza genome, as well as for a concatenation of all segment. While the latter is not expected to yield trees that faithfully reflect the relationship of all viruses and segments to each other, such an analysis of concatenated genomes are nevertheless useful to resolve the relationship of very similar genomes.

The WGS dataset of all sequenced strains were visualized in the Nextstrain tool with demographic and socioeconomic metadata at spatiotemporal resolution – **Table S5** provides all links for data access on nextstrain. We also used the narrative function of Nextstrain for specific highlighting of aspects, namely the phylogenic tree structure across cities, urban quarters and statistical blocks: https://nextstrain-dev.herokuapp.com/community/narratives/appliedmicrobiologyresearch/Influenza-2016-2017/baselFlu and for specific transmission clusters: https://nextstrain-dev.herokuapp.com/community/narratives/appliedmicrobiologyresearch/Influenza-2016-2017/baselFluClusters). All used files can be downloaded at https://github.com/appliedmicrobiologyresearch/FluBasel.

## Acknowledgement

We thank the family doctors and pediatricians helping to recruit the patients for this study: Praxisgemeinschaft Dornacherstrasse (Dr. Burger, Dr. Eggenschwiler, Dr. Wyss, Dr. Gessler, Dr. Nonnenmacher), Praxis Bündnerhof (Dr. Müller, Dr. Peters, Dr. Hantke), Praxisgemeinschaft Banderet-Malè (Dr. Banderet-Uglioni, Dr. Malè), Hammerpraxis (Prof. Zeller), Praxis Schneider/von Hornstein (Dr. Schneider, Dr. von Hornstein), Davidsbodenpraxis (Dr. Amacher, Dr. Hug, Dr. Voelin, Isay, Dr. Pizzagalli, Dr. Navarini), Praxis Türkoglu/Bär (Dr. Türkoglu, Dr. Bär), and Praxis Gordon / Walker (Dr. Gorden, Dr. Walker). We also thank the study nurse team of the Clinical trial unit (Silke Purschke and Karin Wild) for their excellent organization and coordination of the patient recruitment. We thank Magdalena Schneider, Rosamaria Vesco, Christine Kiessling, Elisabeth Schultheiss, and Clarisse Straub for excellent technical assistance of genome sequencing. We also thank Dr. Vladimira Hinic (University Hospital Basel) and Prof. Emma Slack (ETH Zurich) for critically reading the manuscript. We thank Andrew Jermy for excellent feedback and critical review on the manuscript (Germinate, UK).

We also acknowledge the contributing colleagues and centers to GISAID (see https://github.com/appliedmicrobiologyresearch/Influenza-2016-2017/blob/master/data_information/acknowledgement_table.tsv for a full list).

## Author contributions

Study design: AE, MaB, TS, RSS

Data capture: AE, CS, NG, MB, MS, AB, YH, JB, TV, NAS

Strain collection: AE, OD, MN

Whole genome sequencing: DL, DW, HSS

Hemagglutionation inhinition assay: MS, DL

Generation of maps: MB, NA, NS

Nextstrain visualization: RN, EH

Patient recruitment: NR, CHN, AZ, AB

Bioinformatic analysis: DW, EH, RN, NFM, JH, TB, HSS

Statistical analysis: STS

Writing of first draft manuscript: AE, NG

Reviewed the manuscript: MaB, TS, RSS, MB, NFM, DW, RN, HSS, AB, STS

## Data availability statement

All sequencing data (raw reads) will be made available at NCBI. As well tables with anonymized PCR-confirmed cases and anonymized survey results will be made available in a public data repository.

## Code availability statement

All codes used to process the viral genomic data will be made available on github.

## Supplementary material

**Supplementary file 1.** Animation of cumulative PCR-confirmed influenza cases during the Influenza 2017/2018 seasons. Influenza A and B are shown in the same colour. Based on the incidence rates of Figure S4.

**Supplementary file 2.** Questionnaire, English version

**Supplementary file 3.** Submitted paper “Characterising the epidemic spread of Influenza A/H3N2 within a city through phylogenetics” by Müller NF, Wüthrich D, Goldman N et al.

## Supplement methods and results

### Calculations for tables 1 and 2 and table S3

Y serves as placeholder for the outcome variable and x as placeholder for the examined variable. The calculations are performed a modified Poisson regression approach to prospective studies with binary data^59^.

#### Univariable model

glm y x, fam (poisson) link(log)nolog eform robust glm: generalized linear model

fam: family (describes the distribution; here we choose a poisson distribution to estimate relative risks)

link: link function

log: exponentiated coefficients are incidence-rate ratios eform: report exponentiated coefficients

nolog: suppresses the iteration log display robust: robust variance estimator

#### Multivariable model

##### Stepwise forward selection

stepwise, pe(0.05): glm y x1 x2 x3 x(…), fam (poisson) link(log)nolog eform robust

pe: begin with empty model

(0.05): specification of the significance level

All other abbreviations see above

##### Stepwise backward selection

stepwise, pr(0.05): glm y x1 x2 x3 x(…), fam (poisson) link(log)nolog eform robust

Pr: begin with full model

All other abbreviations see above

#### Multilevel model (mixed effects generalized linear model)

meglm y x1 x2 x3 x(…), II urban quarter:, fam (poisson) link(log)nolog eform

Group variable : urban quarter

### Phylodynamic model for transmission

In addition to the previously described clustering model based on the number of single nucleotide polymorphisms, we have also applied a clustering method based on a phylogenetic approach (see attached supplementary publication by Mueller N, Wuethrich D et al.).

## Methods

### Initial clustering based on nucleotide differences

We combined the Basel sequences with a global sample of sequences (downloaded on 17th July 2018) from https://www.gisaid.org which had been sampled between 1st January 2016 and 31st December 2017 for which at least the segments HA, NA and MP were available. We then calculated the average nucleotide difference between any of the sequences and the sequences from Basel. To split the dataset into manageable pieces, we first grouped any two sequences from Basel together if they were within an average nucleotide difference of 0.0025 per position. When the full genome for two sequences was available, this would correspond to about 32 different positions on the full genome. For an average clock rate of 0.005 mutations per site per year, 32 differences correspond to a pairwise phylogenetic distance of about 0.5 years. Sets of sequences from Basel were only split into two groups if the two closest related sequences of each group exceeded this criterium. Based on this initial grouping, we added global sequences to each cluster if they were at maximum 0.0025/2 mutations away from any of the sequences from Basel. Factor 2 was used to reduce the number of global sequences in each of these initial clusters.

### Phylogenetic trees of initial clusters

We next estimated rates of evolution for each segment using the SRD06 model^60^ and a strict clock model, from 200 full genome sequences sampled in California, New York and Europe between 2010 and 2015, using Beast 2.5^61^. Apart from this, these sequences were not otherwise used in the analysis, and as such are an independent dataset. We allowed each segment to have its own phylogeny in order to avoid the possibility of a reassortment biasing the estimates of evolutionary rates. Each of the segments, as well as the first two and third codon position was allowed to have its own rate scaler. We ran ten independent Markov Chain Monte Carlo (MCMC) chains each for 10^8^ iterations and then combined them after a burn-in of 10%. These estimated evolutionary rates are long-term rates of influenza A/H3N2.

We next reconstructed the phylogenetic trees of all initial clusters by using the full genomes of all samples from the initial clusters. We fixed the evolutionary rates to be equal to the mean evolutionary rates as estimated using the methods above. As a population model, we used a constant coalescent model with an estimated effective population size that was shared among all initial clusters. We then estimated a distribution of phylogenies for each initial cluster, assuming that all segments share the same phylogeny. As estimated in the previous analysis, reassortment will not bias evolutionary rates.

### Local cluster identification

To identify sets of sequences from Basel that were likely to have been transmitted locally, we reconstructed the geographic origins of lineages that were introduced into Basel. Therefore, we used the phylogenetic tree distributions for each initial cluster to reconstruct the ancestral states using parsimony. We made some modifications to the standard algorithm for parsimony ancestral state reconstruction to reflect our prior assumption, that Basel is unlikely to act as a relevant source of influenza on a global scale. If any descendent was not in Basel, the node was classified as “not Basel”. Since the influenza season covers only a few months, we additionally assumed that lineages are unlikely to persist in Basel for more than 5 weeks (0.1 years) without being sampled. To reflect this assumption, we classified internal nodes that are more than 0.1 years from a sample from Basel to be either outside of Basel, or to be in an unknown location. We then defined sequences to be in the same local cluster if all their ancestors are inferred to be in Basel. Local clusters are obtained for each iteration of the MCMC. The exact workflow, including BEAST2 input files can be found at https://github.com/nicfel/Flu-Basel/LocalClusters. While alternative model-based approaches exist to reconstruct locations of internal nodes^62^, these approaches themselves make strict assumptions that are violated when studying the spread of diseases on a city scale. Also, it is further unknown how well they perform when migration between individual locations is very strong. From the grouping of sequences into local clusters as described above, sequences were classified into different local clusters over the course of the MCMC.

## Results

Using clusters obtained from a phylodynamic approach, we obtain a similar clustering pattern meaning the pattern is robust towards how we define clusters. In particular, using the phylodynamic method, we identified 96 clusters of at least two highly similar viral genomes, representing influenza transmission, which together incorporated 534/663 influenza strains. We determined local transmission clusters by identifying groups of local isolates that are phylogenetically more closely related to each other than to any isolate from the global collection from GISAID (www.gisaid.org). Some of the clusters included isolates from both within and outside of the city, whereas other clusters were focused mainly within the city. Most of the clusters contained samples from different urban quarters, and only a few clusters were predominantly located within a single urban quarter (see Cluster A to C, **Figure S8**). Within the city, we observed 323/427 (76%) of influenza strains belong to 69 of the overall 96 clusters across this single influenza season. The clusters comprised a median of 3 isolates, ranging from 2 to 42 within and outside of the city.

---

**Figure S1:**
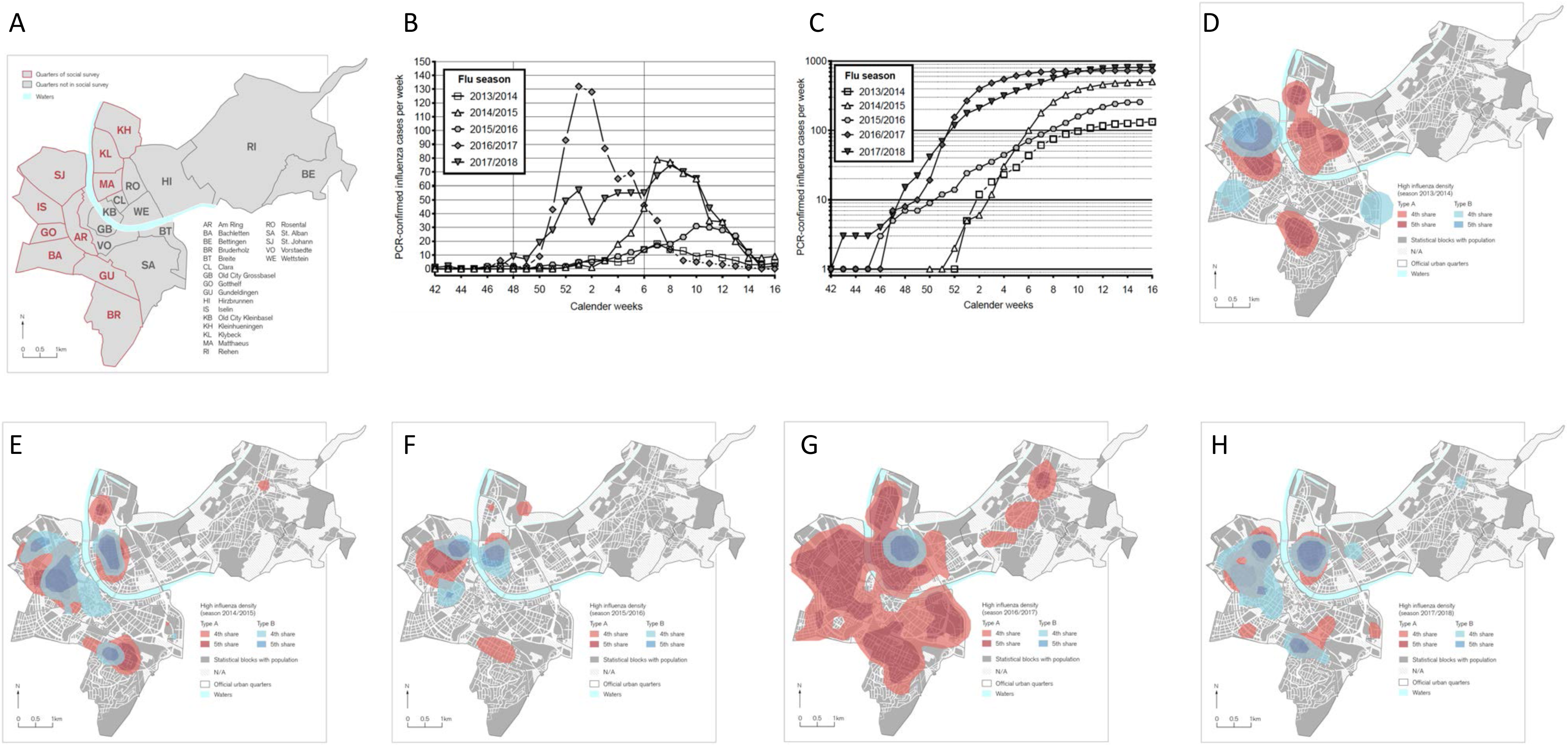
Influenza cases and distribution in the City of Basel per season from 2013/2014 to 2017/2018. **(A)** Map of urban quarters showing the study area. Urban quarters included into the household survey are marked in red. **(B)** Absolute numbers of PCR-confirmed influenza for each season. **(C)** Cumulative influenza cases for each season. **(D-H)** Influenza cases per statistical block shown as kernel density estimates for each influenza season.

**Figure S2:**
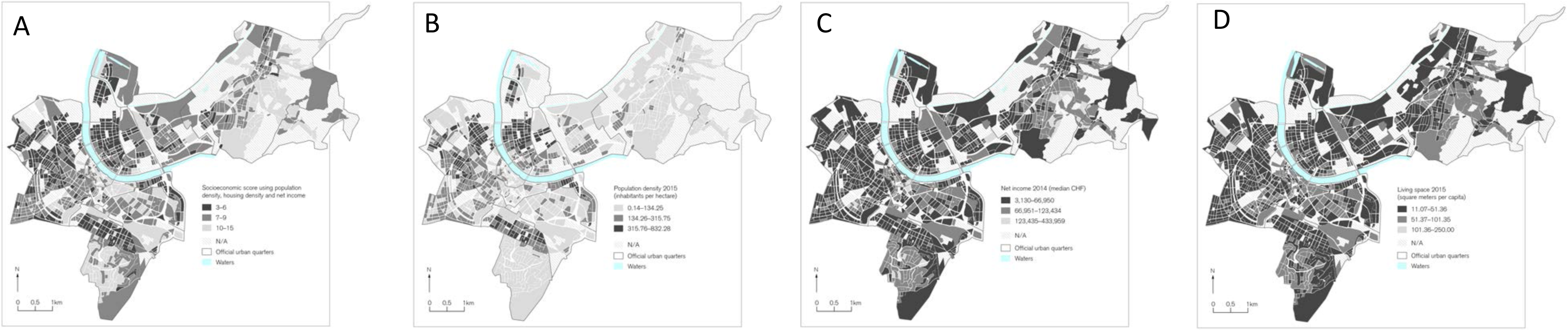
Socioeconomic data of the City of Basel. Combined and individual socioeconomic scores are shown for each statistical block. For each map, the natural breaks of the individual factors is shown. **(A)** Combination of socioeconomic factors. **(B)** Living space in m^2^ per capita. **(C)** Net income in CHF per statistical block. **(D)** Population density as inhabitants per hectare (ha).

**Figure S3:**
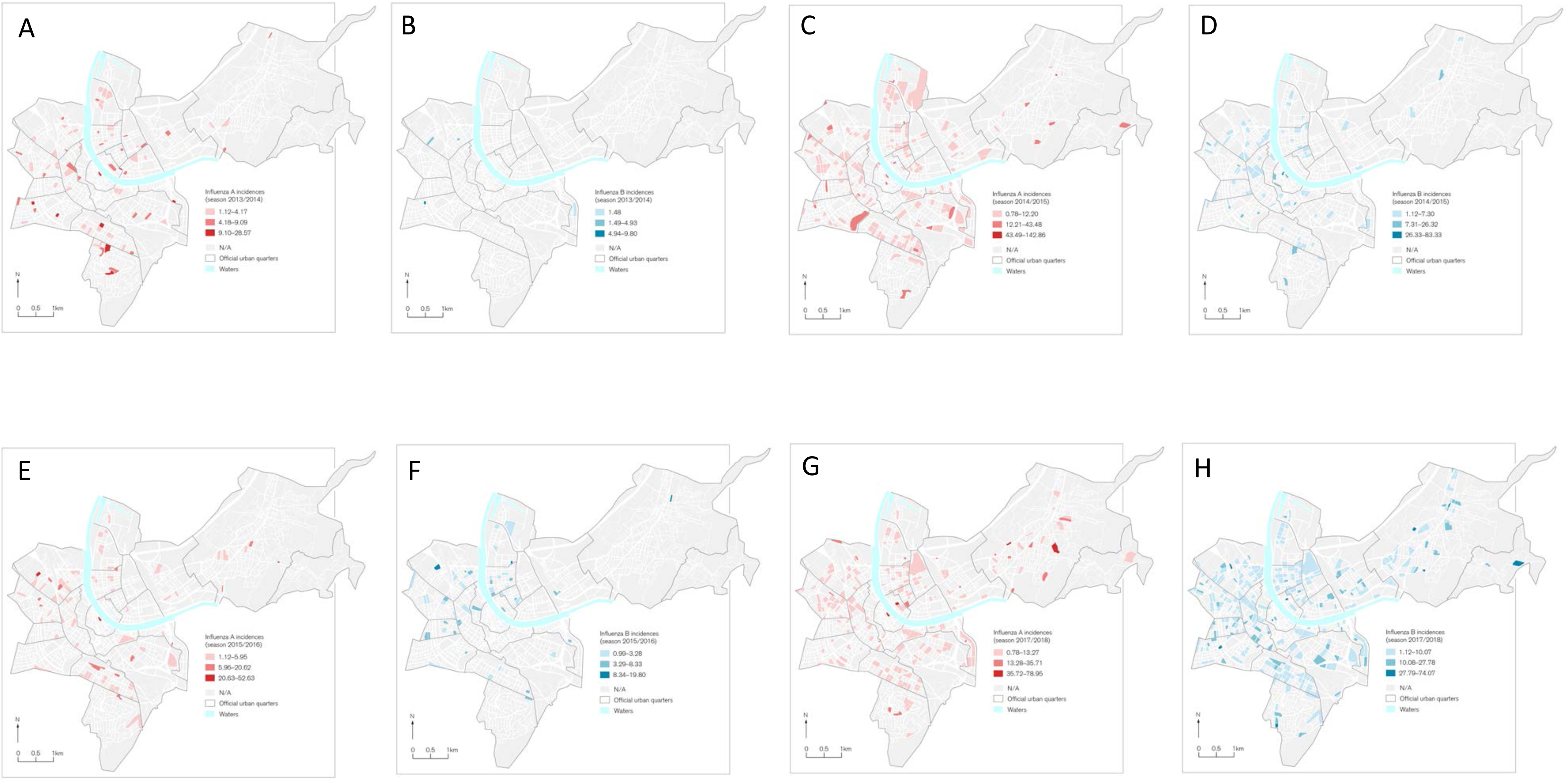
Influenza incidence rates in the City of Basel per season from 2013/2014 to 2017/2018. **(A-H)** PCR-confirmed influenza incidence rates (per 1000 inhabitants) for each season per urban quarter.

**Figure S4:**
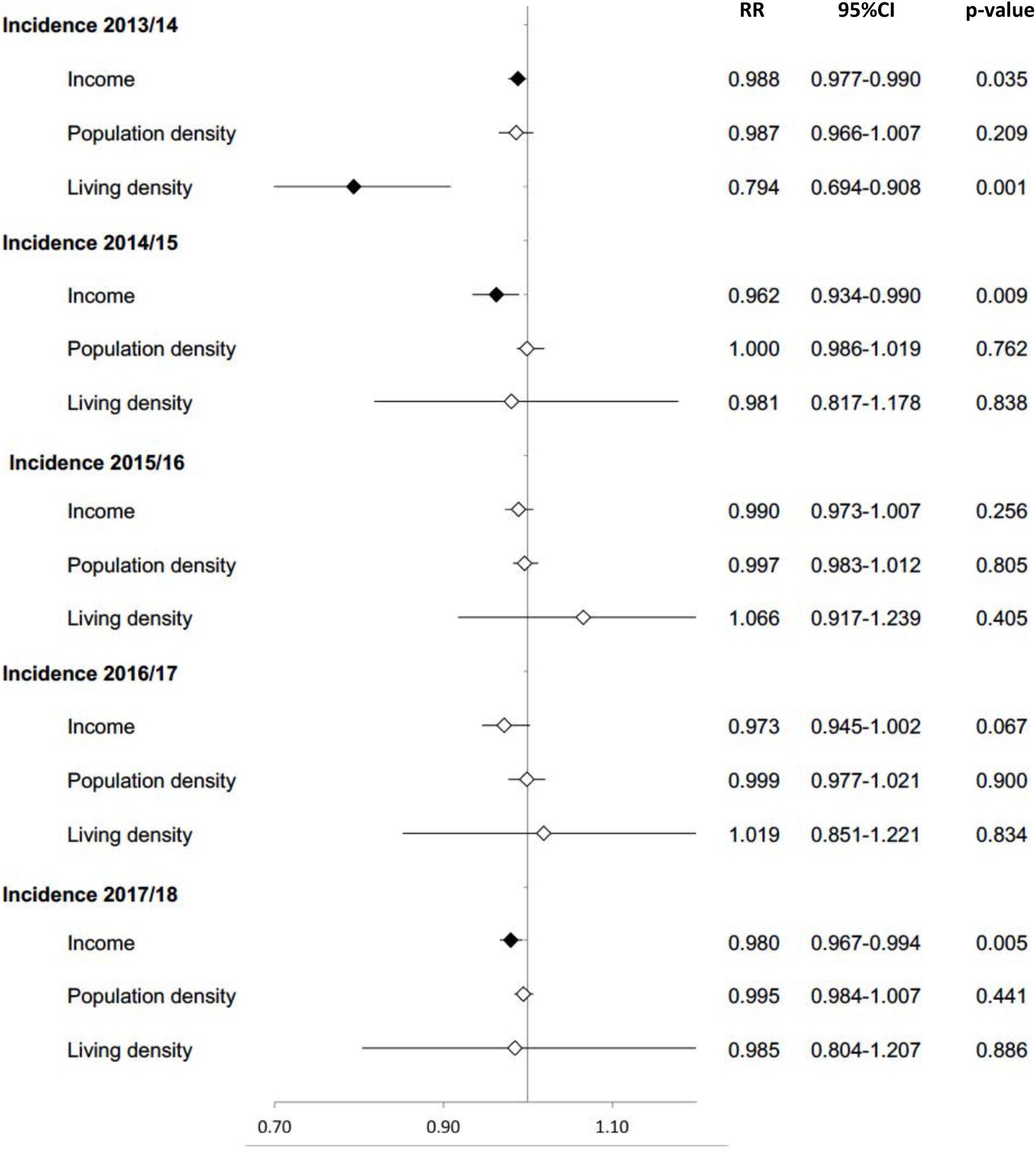
Multivariable Poisson regression for influenza incidence in each influenza season from 2013/2014 to 2017/2018. The correlation of each socioeconomic factors – income, population density, and living density – with influenza incidence is expressed as relative risk (RR).

**Figure S5:**
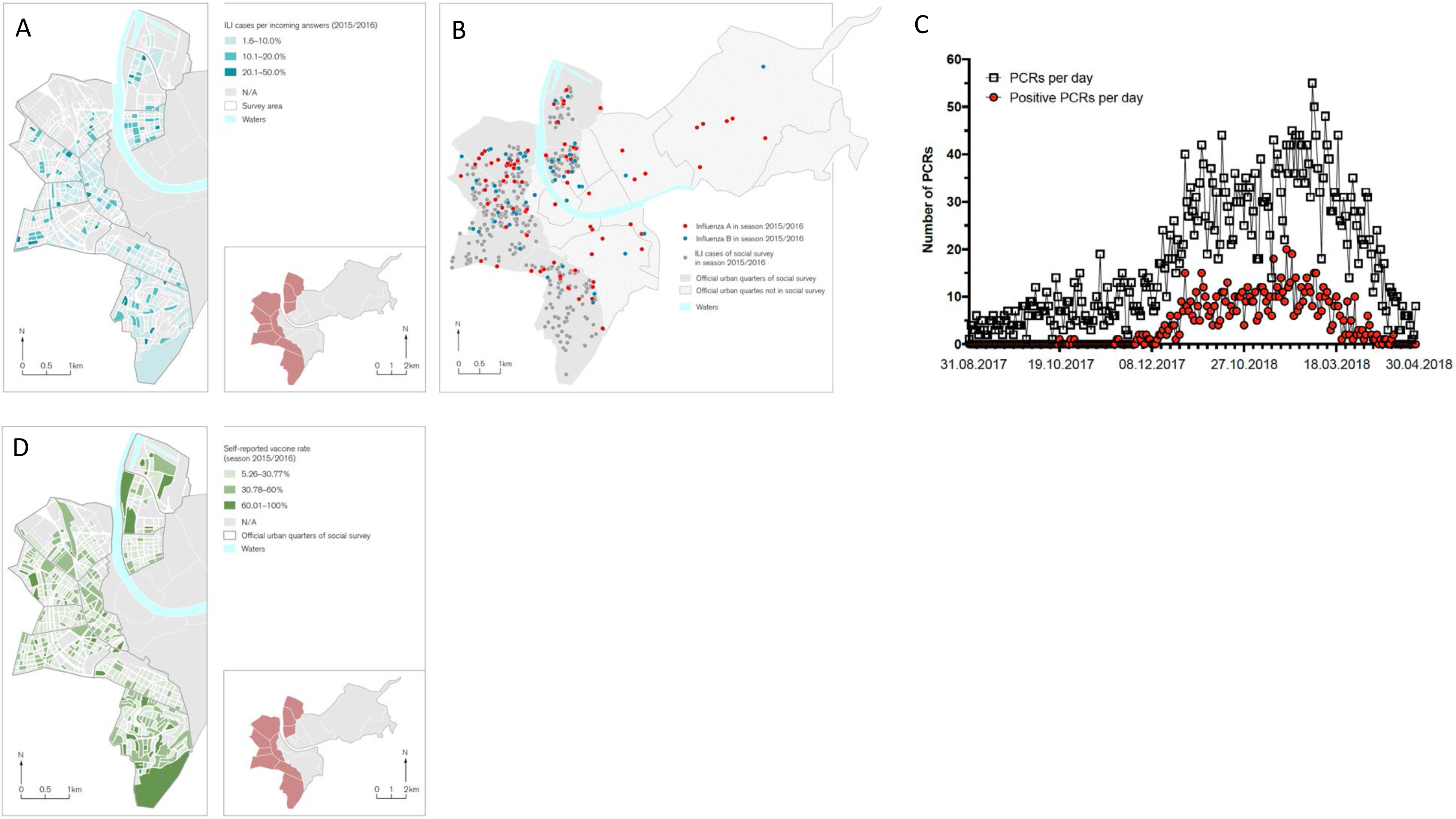
Survey for self-reported ILI and vaccination against influenza. **(A)** Self-reported influenza-like illness cases corrected per returned questionnaire per statistical block. **(B)** Self-reported ILI and PCR-confirmed influenza infections for the 2015/2016 influenza season. ILI cases according to WHO definition are shown in grey for 10 selected urban quarters and PCR-confirmed Influenza A as red dots and Influenza B as blue dots. **(C)** Ratio of tested PCRs and positive results (PCR-confirmed) in a single season. **(D)** Self-reported vaccination corrected per returned questionnaire per statistical block.

**Figure S6:**
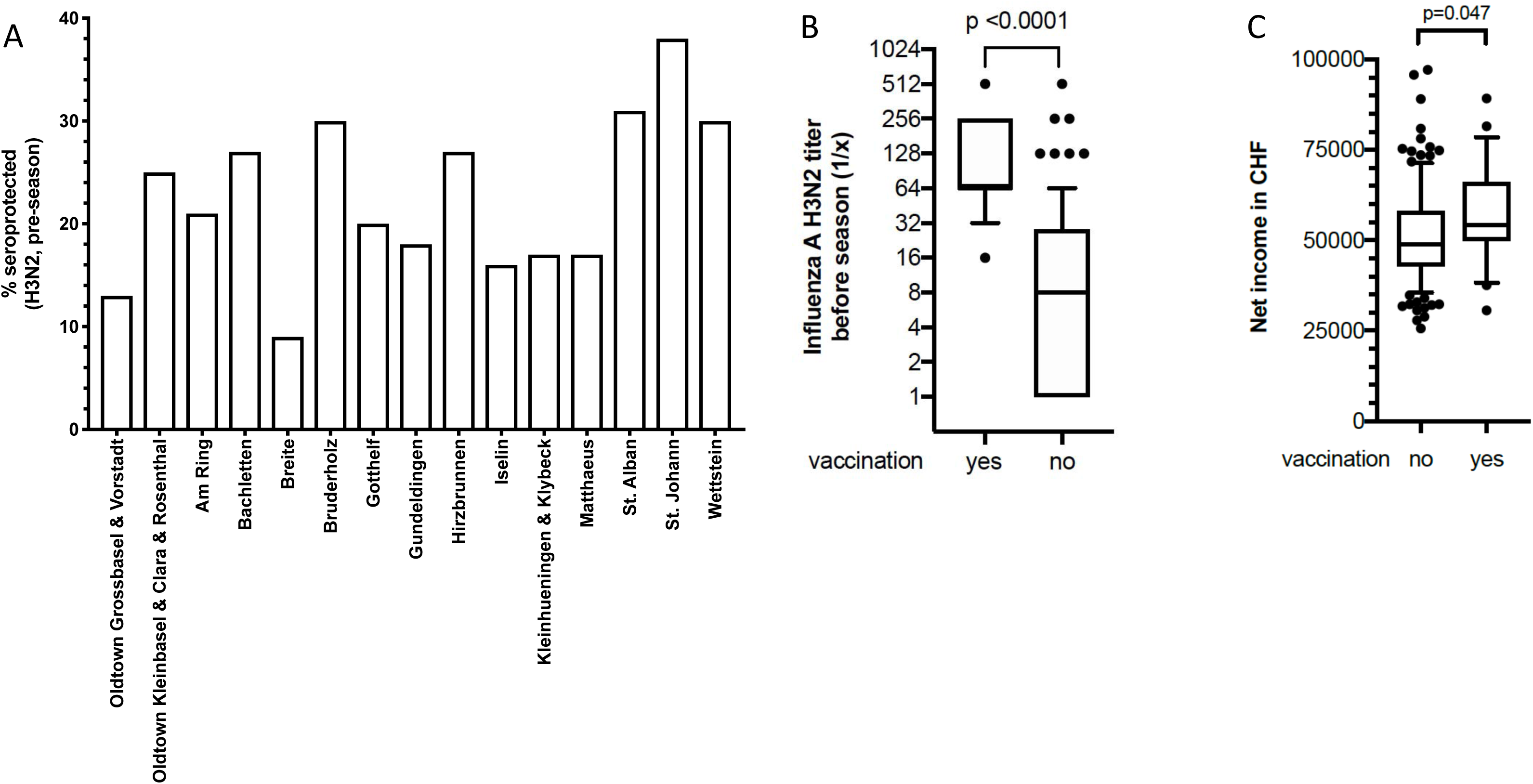
Pre-seasonal influenza antibody titres measured by hemagglutination inhibition assay in season 2016/2017. **(A)** Percentage of seroprotected healthy blood donors (HIA titre of 1:40 or above), prior to the influenza season, by urban quarter. As for smaller urban quarters not enough measurements were available, these were combined based on geographical proximity and socioeconomic similarity. **(B)** Hemagglutination inhibition antibody titres in healthy blood donors before the 2016/2017 season. Influenza A H3N2 titres against the circulating virus of that season is shown in vaccinated and non-vaccinated people. **(C)** Net income in CHF per statistical block shown for vaccinated and non-vaccinated healthy blood donors in the 2016/2017 season.

**Figure S7:**
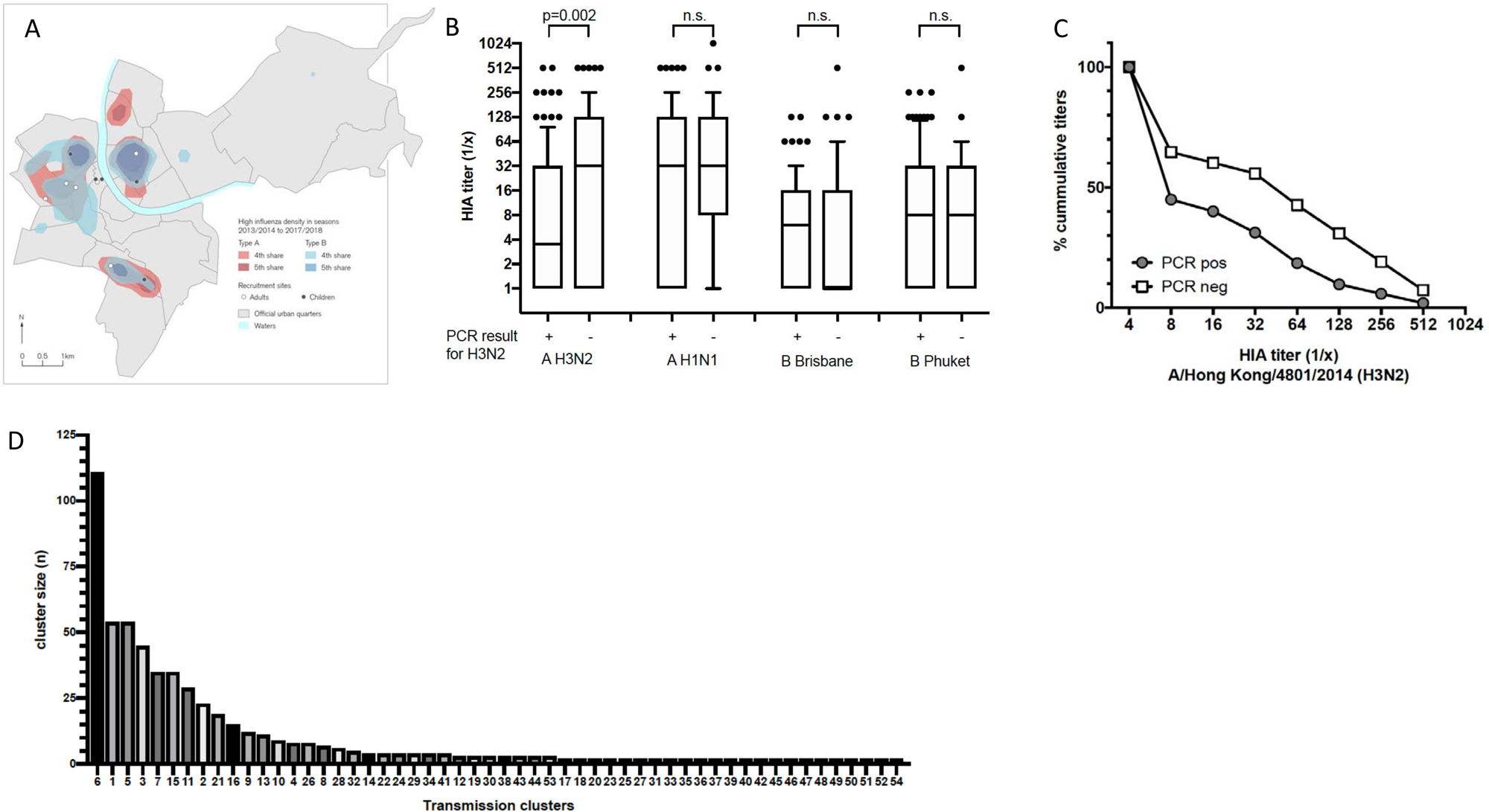
Whole genome sequencing of influenza viruses during the 2016/2017 season. **(A)** The distribution of study recruitment sites for children (dark grey dots) and adults (white dots), and the kernel density estimates of overall influenza burden in the city. **(B)** Hemagglutination inhibition antibody titres against the circulating Influenza A H3N2 virus and all other vaccinated Influenza viruses in each patient enrolled with influenza-like illness during the 2016/2017 season. Patients with PCR-confirmed influenza are compared against patients with negative PCR testing. All box blots show median and interquartile ranges. Whiskers indicate 10-90% percentile with outliers. **(C)** The percentage of cumulative H3N2 specific hemagglutination inhibition antibody titres is shown, again highlighting the difference between PCR positive and negative titres. **(D)** Number of clusters and cluster size are shown, from the whole genome sequencing data. More than 50% of the clusters comprise five or fewer viruses.

**Figure S8:**
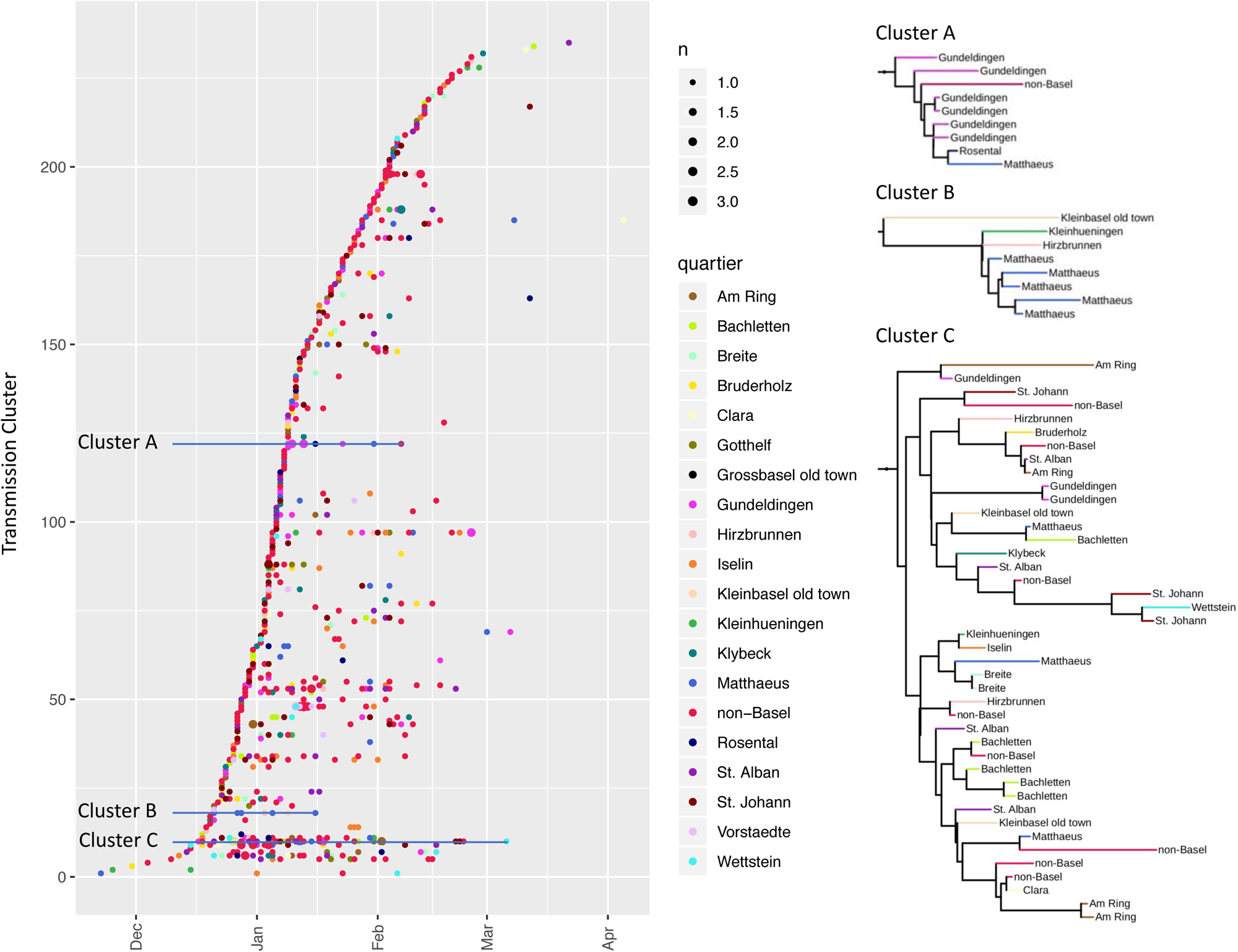
Cluster analysis of each influenza case genome during the 2016/2017 season based on the phylodynamic model. Each dot represents a sequenced influenza virus isolate. Colour reflects the host address by specific urban quarter. Each horizontal line corresponds to a local transmission cluster of influenza viruses. Three representative clusters A, B and C are detailed on the right using maximum likelihood trees indicating the complexity of transmission clusters across time and urban quarters.

**Table S1.** Influenza virus types per season in Basel and Europe.

|  | **City of Basel** | **Switzerland**<sup>54-58</sup> | **Europe**<sup>59-63</sup> |
| --- | --- | --- | --- |
| **2013/2014** | Influenza A (n=136, 97%)<br>Influenza B (n=5, 3%) | Total samples n=580<br>Influenza A (98%): H1N1 41%, H3N2 56%<br>Influenza B (2%): Yamagata 2%, Victoria 0% | Total samples n=34,210<br>Influenza A (94%): H1N1 41%, H3N2 47%<br>Influenza B (6%): Yamagata 1%, Victoria 0% |
| **2014/2015** | Influenza A (n=226, 67%)<br>Influenza B (n=110, 23%) | Total samples n=937<br>Influenza A (71%): H1N1 14%, H3N2 56%<br>Influenza B (29%): Yamagata 27%, Victoria 1% | Total samples n=40,931<br>Influenza A (68%): H1N1 15%, H3N2 49%<br>Influenza B (32%): Yamagata 8%, Victoria 0% |
| **2015/2016** | Influenza A (n=92, 62%)<br>Influenza B (n=56, 38%) | Total samples n=975<br>Influenza A (35%): H1N1 30% H3N2 5%<br>Influenza B (64%): Yamagata 2%, Victoria 62% | Total samples n=56,892<br>Influenza A (59%): H1N1 49%, H3N2 7%<br>Influenza B (41%): Yamagata 1%, Victoria 18% |
| **2016/2017** | Influenza A (n=501, 99%)<br>Influenza B (n=5, 1%) | Total samples n=982<br>Influenza A (96%): H1N1 2% H3N2 94%<br>Influenza B (4%): Yamagata 3%, Victoria 0% | Total samples n=49,410<br>Influenza A (90%): H1N1 1%, H3N2 75%<br>Influenza B (10%): Yamagata 2%, Victoria 2% |
| **2017/2018** | Influenza A (n=268, 38%)<br>Influenza B (n=433, 62%) | Total samples n=1,292<br>Influenza A (29%): H1N1 23%, H3N2 5%<br>Influenza B (71%): Yamagata 66%, Victoria 0% | Total samples n=60,658<br>Influenza A (37%): H1N1 20%, H3N2 11%<br>Influenza B (63%): Yamagata 29%, Victoria 1% |

**Table S2.** Univariable analysis of factors associated with self-reported influenza-like illness.

| Variables | ILI yes |  | ILI no |  | Relative Risk | 95% CI | p-Value |
| --- | --- | --- | --- | --- | --- | --- | --- |
|  | n | in % | n | in % |  |  |  |
| Vaccination (yes) | 37 | 1.8 | 2036 | 98.2 | 0.33 | 0.24-0.46 | <0.001 |
| Risk group based on ATC-level (yes) | 52 | 3.0 | 1678 | 97.0 | 0.59 | 0.37-0.95 | 0.029 |
| Smoker |  |  |  |  | 1.33 | 1.16-1.51 | <0.001 |
| no | 250 | 4.0 | 5971 | 96.0 |  |  |  |
| yes, sometimes | 47 | 5.3 | 837 | 94.7 |  |  |  |
| yes, daily | 61 | 7.1 | 801 | 92.9 |  |  |  |
| Alcohol consumption |  |  |  |  | 0.78 | 0.66-0.92 | 0.003 |
| no | 825 | 4.9 | 1582 | 95.1 |  |  |  |
| yes, sometimes | 259 | 4.7 | 5261 | 95.3 |  |  |  |
| yes, daily | 15 | 2.0 | 742 | 98.0 |  |  |  |
| Implementation of hand washing |  |  |  |  | 1.05 | 0.91-1.21 | 0.510 |
| Very bad | 0 | 0.0 | 19 | 0.0 |  |  |  |
| Bad | 4 | 5.8 | 65 | 94.2 |  |  |  |
| Average | 33 | 4.1 | 770 | 95.9 |  |  |  |
| Good | 126 | 4.6 | 2639 | 95.4 |  |  |  |
| Very good | 193 | 4.7 | 3939 | 95.3 |  |  |  |
| Implementation of health check-ups |  |  |  |  | 0.82 | 0.75-0.89 | <0.001 |
| Very bad | 56 | 6.1 | 856 | 93.9 |  |  |  |
| Bad | 65 | 6.8 | 886 | 93.2 |  |  |  |
| Average | 91 | 4.8 | 1809 | 95.2 |  |  |  |
| Good | 65 | 3.6 | 1731 | 96.4 |  |  |  |
| Very good | 33 | 3.0 | 1061 | 97.0 |  |  |  |
| Implementation of healthy diet |  |  |  |  | 0.82 | 0.73-0.93 | 0.002 |
| Very bad | 1 | 5.6 | 17 | 94.4 |  |  |  |
| Bad | 10 | 8.3 | 110 | 91.7 |  |  |  |
| Average | 63 | 4.8 | 1252 | 95.2 |  |  |  |
| Good | 189 | 5.2 | 3475 | 94.8 |  |  |  |
| Very good | 94 | 3.5 | 2594 | 96.5 |  |  |  |
| Implementation of physical activity |  |  |  |  | 0.83 | 0.75-0.02 | <0.001 |
| Very bad | 3 | 3.4 | 84 | 96.6 |  |  |  |
| Bad | 34 | 7.7 | 409 | 92.3 |  |  |  |
| Average | 94 | 5.3 | 1671 | 94.7 |  |  |  |
| Good | 137 | 4.7 | 2761 | 95.3 |  |  |  |
| Very good | 90 | 3.6 | 2395 | 96.4 |  |  |  |
| Public transport use |  |  |  |  | 1.17 | 1.07-1.29 | 0.001 |
| Never | 2 | 1.4 | 144 | 98.6 |  |  |  |
| Rare | 47 | 3.7 | 1214 | 96.3 |  |  |  |
| Multiple times a month | 69 | 4.1 | 1598 | 95.9 |  |  |  |
| Multiple times a week | 92 | 4.4 | 2007 | 95.6 |  |  |  |
| Daily | 142 | 5.6 | 2375 | 94.4 |  |  |  |
| Contact with more than 50 persons a day | 86 | 6.4 | 1250 | 93.6 | 1.57 | 1.24-1.98 | <0.001 |
| Healthcare worker | 44 | 4.6 | 912 | 95.4 | 0.79 | 0.57-1.08 | 0.135 |
| work with children | 40 | 5.2 | 728 | 94.8 | 0.90 | 0.65-1.25 | 0.541 |
| Work in an open plan office | 135 | 6.2 | 2040 | 93.8 | 1.19 | 0.94-1.49 | 0.147 |
| Gender |  |  |  |  | 0.92 | 0.74-1.13 |  |
| Male | 129 | 4.2 | 2907 | 95.8 |  |  | 0.423 |
| Female | 229 | 4.6 | 4715 | 95.4 |  |  |  |
| old age ($\geq 65$ years) | 36 | 1.5 | 2349 | 98.5 | 0.25 | 0.18-0.36 | <0.001 |
| Foreign population | 93 | 5.8 | 1508 | 94.2 | 1.41 | 1.12-1.77 | 0.004 |
| living space ($\leq 30$ m<sup>2</sup>/capita) | 48 | 7.8 | 566 | 92.2 | 1.71 | 1.26-2.34 | 0.001 |
| $\geq 3$ persons / household | 137 | 7.5 | 1700 | 92.5 | 2.06 | 1.68-2.54 | <0.001 |
| Lower educational level (compulsory) | 13 | 3.2 | 391 | 96.8 | 0.70 | 0.40-1.20 | 0.195 |
| Upper and middle management | 63 | 4.7 | 1270 | 95.3 | 0.80 | 0.61-1.05 | 0.108 |
| Low Income (≤ CHF 6000 gross household income) | 150 | 4.8 | 2985 | 95.2 | 1.00 | 0.80-1.25 | 0.995 |
| Share of green areas ($\leq 20\%$ / residential block) | 85 | 5.1 | 1590 | 94.9 | 0.83 | 0.65-1.05 | 0.121 |
| Share of built up area ($\geq 60\%$ / residential block) | 29 | 3.9 | 721 | 96.1 | 0.87 | 0.60-1.27 | 0.472 |

**Table S3.** Univariable analysis of factors associated with self-reported vaccination.

| Variables | vaccination<br>yes |  | vaccination<br>no |  | Relative<br>Risk | 95% CI | p-Value |
| --- | --- | --- | --- | --- | --- | --- | --- |
|  | n | in % | n | in % |  |  |  |
| Risk group based on ATC-level | 826 | 47.1 | 929 | 52.9 | 1.75 | 1.50-2.03 | <0.001 |
| Smoker |  |  |  |  | 0.68 | 0.63-0.74 | <0.001 |
| no | 1819 | 28.9 | 4473 | 71.1 |  |  |  |
| yes, sometimes | 150 | 16.7 | 749 | 83.3 |  |  |  |
| yes, daily | 130 | 14.9 | 742 | 85.1 |  |  |  |
| Alcohol consumption |  |  |  |  | 1.03 | 0.95-1.100 | 0.477 |
| no | 479 | 28.3 | 1215 | 71.7 |  |  |  |
| yes, sometimes | 1356 | 24.3 | 4222 | 75.7 |  |  |  |
| yes, daily | 254 | 33.1 | 513 | 66.9 |  |  |  |
| Implementation of hand washing |  |  |  |  | 1.11 | 1.05-1.17 | <0.001 |
| Very bad | 4 | 20.0 | 16 | 80.0 |  |  |  |
| Bad | 16 | 23.2 | 53 | 76.8 |  |  |  |
| Average | 187 | 23.0 | 626 | 77.0 |  |  |  |
| Good | 675 | 24.2 | 2113 | 75.8 |  |  |  |
| Very good | 1160 | 27.2 | 3033 | 72.3 |  |  |  |
| Implementation of health check-ups |  |  |  |  | 1.37 | 1.32-1.42 | <0.001 |
| Very bad | 128 | 13.9 | 792 | 86.1 |  |  |  |
| Bad | 163 | 16.9 | 801 | 83.1 |  |  |  |
| Average | 439 | 22.8 | 1487 | 77.2 |  |  |  |
| Good | 600 | 33.2 | 1209 | 66.8 |  |  |  |
| Very good | 502 | 45.1 | 611 | 54.9 |  |  |  |
| Implementation of healthy diet |  |  |  |  | 0.86 | 0.83-0.91 | <0.001 |
| Very bad | 6 | 31.6 | 13 | 68.4 |  |  |  |
| Bad | 38 | 31.9 | 81 | 68.1 |  |  |  |
| Average | 413 | 31.0 | 920 | 69.0 |  |  |  |
| Good | 940 | 25.4 | 2760 | 74.6 |  |  |  |
| Very good | 614 | 22.5 | 2113 | 77.5 |  |  |  |
| Implementation of physical activity |  |  |  |  | 0.91 | 0.87-0.95 | <0.001 |
| Very bad | 24 | 27.0 | 65 | 73.0 |  |  |  |
| Bad | 131 | 29.4 | 314 | 70.6 |  |  |  |
| Average | 515 | 28.7 | 1278 | 71.3 |  |  |  |
| Good | 712 | 24.4 | 2211 | 75.6 |  |  |  |
| Very good | 574 | 22.8 | 1946 | 77.2 |  |  |  |
| Public transport use |  |  |  |  | 1.05 | 1.01-1.08 | 0.005 |
| Never | 40 | 27.2 | 107 | 72.8 |  |  |  |
| Rare | 265 | 20.8 | 1010 | 79.2 |  |  |  |
| Multiple times a month | 423 | 25.0 | 1268 | 75.0 |  |  |  |
| Multiple times a week | 643 | 30.3 | 1480 | 69.7 |  |  |  |
| Daily | 643 | 25.3 | 1903 | 74.4 |  |  |  |
| Contact with more than 50 persons a day | 281 | 20.8 | 1069 | 79.2 | 0.78 | 0.70-0.88 | <0.001 |
| Healthcare worker | 323 | 33.4 | 645 | 66.6 | 2.40 | 2.13-2.70 | <0.001 |
| work with children | 111 | 14.3 | 667 | 85.7 | 0.78 | 0.65-0.94 | 0.008 |
| Work in an open plan office | 396 | 18.0 | 1810 | 82.0 | 1.02 | 0.91-1.16 | 0.708 |
| Gender |  |  |  |  | 1.13 | 1.05-1.22 | 0.001 |
| Male | 857 | 27.9 | 2217 | 72.1 |  |  |  |
| Female | 1235 | 24.7 | 3767 | 75.3 |  |  |  |
| old age ($\geq 65$ years) | 1116 | 46.1 | 1303 | 53.9 | 2.72 | 2.53-2.93 | <0.001 |
| Foreign population | 377 | 23.2 | 1249 | 76.8 | 0.87 | 0.79-0.96 | 0.005 |
| living space ($\leq 30\text{m}^2/\text{capita}$) | 89 | 14.4 | 527 | 85.6 | 0.56 | 0.46-0.69 | <0.001 |
| ≥ 3 persons / household | 359 | 19.3 | 1500 | 80.7 | 0.69 | 0.63-0.77 | <0.001 |
| Lower educational level (compulsory) | 168 | 40.3 | 249 | 59.7 | 1.62 | 1.43-1.83 | <0.001 |
| Upper and middle management | 297 | 22.0 | 1051 | 78.0 | 1.37 | 1.20-1.55 | <0.001 |
| Low Income (≤ CHF 6000.- gross household income) | 673 | 21.3 | 2491 | 78.7 | 0.79 | 0.72-0.86 | <0.001 |
| Share of green areas (≤ 20% / residential block) | 375 | 22.1 | 1324 | 77.9 | 1.22 | 1.10-1.34 | <0.001 |
| Share of built up area (≥ 60% / residential block) | 161 | 21.2 | 597 | 78.8 | 0.81- | 0.70-0.93 | 0.003 |

**Table S4.**
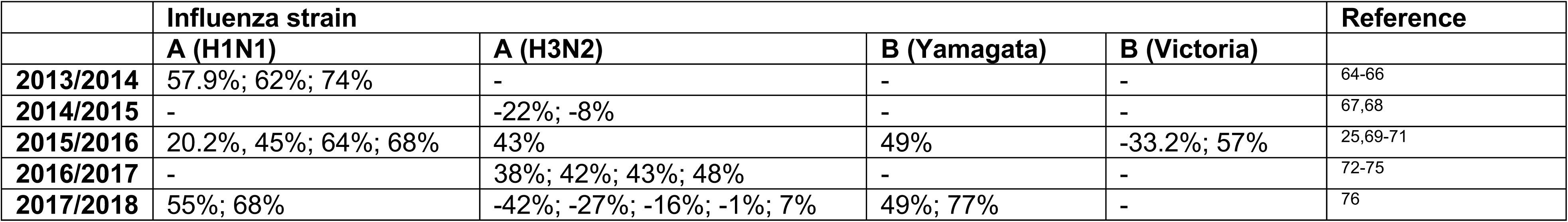
Vaccine effectiveness from 2013 to 2018.

**Table S5.** List of all link to Nextstrain for visualization of phylogenetic trees.

| **Viral segment** | **Links** |
| --- | --- |
| all | https://nextstrain-dev.herokuapp.com/community/appliedmicrobiologyresearch/Influenza-2016-2017/h3n2/full |
| HA | https://nextstrain-dev.herokuapp.com/community/appliedmicrobiologyresearch/Influenza-2016-2017/h3n2/ha |
| NA | https://nextstrain-dev.herokuapp.com/community/appliedmicrobiologyresearch/Influenza-2016-2017/h3n2/na |
| NP | https://nextstrain-dev.herokuapp.com/community/appliedmicrobiologyresearch/Influenza-2016-2017/h3n2/np |
| PA | https://nextstrain-dev.herokuapp.com/community/appliedmicrobiologyresearch/Influenza-2016-2017/h3n2/pa |
| PB1 | https://nextstrain-dev.herokuapp.com/community/appliedmicrobiologyresearch/Influenza-2016-2017/h3n2/pb1 |
| PB2 | https://nextstrain-dev.herokuapp.com/community/appliedmicrobiologyresearch/Influenza-2016-2017/h3n2/pb2 |

## References

1 Geoghegan, J. L. et al. Continental synchronicity of human influenza virus epidemics despite climatic variation. PLoS Pathog. 14, e1006780, doi: 10.1371/journal.ppat.1006780 (2018).

2 Peci, A. et al. Effects of Absolute Humidity, Relative Humidity, Temperature, and Wind Speed on Influenza Activity in Toronto, Ontario, Canada. Appl. Environ. Microbiol. 85, doi: 10.1128/AEM.02426-18 (2019).

3 Roosa, K. & Chowell, G. Assessing parameter identifiability in compartmental dynamic models using a computational approach: application to infectious disease transmission models. Theor. Biol. Med. Model. 16, 1, doi: 10.1186/s12976-018-0097-6 (2019).

4 Langat, P. et al. Genome-wide evolutionary dynamics of influenza B viruses on a global scale. PLoS Pathog. 13, e1006749, doi: 10.1371/journal.ppat.1006749 (2017).

5 Lewis, N. S. et al. The global antigenic diversity of swine influenza A viruses. Elife 5, e12217, doi: 10.7554/eLife.12217 (2016).

6 Neher, R. A., Bedford, T., Daniels, R. S., Russell, C. A. & Shraiman, B. I. Prediction, dynamics, and visualization of antigenic phenotypes of seasonal influenza viruses. Proc. Natl. Acad. Sci. U. S. A. 113, E1701–1709, doi: 10.1073/pnas.1525578113 (2016).

7 Vijaykrishna, D. et al. The contrasting phylodynamics of human influenza B viruses. Elife 4, e05055, doi: 10.7554/eLife.05055 (2015).

8 Virk, R. K. et al. Molecular Evidence of Transmission of Influenza A/H1N1 2009 on a University Campus. PLoS One 12, e0168596, doi: 10.1371/journal.pone.0168596 (2017).

9 McCrone, J. T. et al. Stochastic processes constrain the within and between host evolution of influenza virus. Elife 7, doi: 10.7554/eLife.35962 (2018).

10 Poon, L. L. et al. Viral genetic sequence variations in pandemic H1N1/2009 and seasonal H3N2 influenza viruses within an individual, a household and a community. J. Clin. Virol. 52, 146–150, doi: 10.1016/j.jcv.2011.06.022 (2011).

11 Thai, P. Q. et al. Pandemic H1N1 virus transmission and shedding dynamics in index case households of a prospective Vietnamese cohort. J. Infect. 68, 581–590, doi: 10.1016/j.jinf.2014.01.008 (2014).

12 Valley-Omar, Z. et al. Intra-host and intra-household diversity of influenza A viruses during household transmissions in the 2013 season in 2 peri-urban communities of South Africa. PLoS One 13, e0198101, doi: 10.1371/journal.pone.0198101 (2018).

13 Adeola, O. A., Olugasa, B. O., Emikpe, B. O. & Folitse, R. D. Syndromic survey and molecular analysis of influenza viruses at the human-swine interface in two West African cosmopolitan cities suggest the possibility of bidirectional interspecies transmission. Zoonoses Public Health 66, 232–247, doi: 10.1111/zph.12559 (2019).

14 Dalziel, B. D. et al. Urbanization and humidity shape the intensity of influenza epidemics in U.S. cities. Science 362, 75–79, doi: 10.1126/science.aat6030 (2018).

15 Todd, S. et al. Primary care influenza-like illness surveillance in Ho Chi Minh City, Vietnam 2013-2015. Influenza Other Respir Viruses, doi: 10.1111/irv.12574 (2018).

16 Chen, T. M. et al. The transmissibility estimation of influenza with early stage data of small-scale outbreaks in Changsha, China, 2005-2013. Epidemiol. Infect. 145, 424–433, doi: 10.1017/S0950268816002508 (2017).

17 Uchida, M. et al. High vaccination coverage is associated with low epidemic level of seasonal influenza in elementary schools: an observational study in Matsumoto City, Japan. BMC Infect. Dis. 18, 128, doi: 10.1186/s12879-018-3025-9 (2018).

18 Goncalves, L. et al. Urban Planning and Health Inequities: Looking in a Small-Scale in a City of Cape Verde. PLoS One 10, e0142955, doi: 10.1371/journal.pone.0142955 (2015).

19 Wilking, H., Hohle, M., Velasco, E., Suckau, M. & Eckmanns, T. Ecological analysis of social risk factors for Rotavirus infections in Berlin, Germany, 2007-2009. Int J Health Geogr 11, 37, doi: 10.1186/1476-072X-11-37 (2012).

20 Egli, A. et al. Identification of influenza urban transmission patterns by geographical, epidemiological and whole genome sequencing data: protocol for an observational study. BMJ Open 9, e030913, doi: 10.1136/bmjopen-2019-030913 (2019).

21 Park, J. E. & Ryu, Y. Transmissibility and severity of influenza virus by subtype. Infect. Genet. Evol. 65, 288–292, doi: 10.1016/j.meegid.2018.08.007 (2018).

22 Huang, Q. S. et al. Risk Factors and Attack Rates of Seasonal Influenza Infection: Results of the Southern Hemisphere Influenza and Vaccine Effectiveness Research and Surveillance (SHIVERS) Seroepidemiologic Cohort Study. J. Infect. Dis. 219, 347–357, doi: 10.1093/infdis/jiy443 (2019).

23 Somes, M. P., Turner, R. M., Dwyer, L. J. & Newall, A. T. Estimating the annual attack rate of seasonal influenza among unvaccinated individuals: A systematic review and meta-analysis. Vaccine 36, 3199–3207, doi: 10.1016/j.vaccine.2018.04.063 (2018).

24 Zurcher, K., Zwahlen, M., Berlin, C., Egger, M. & Fenner, L. Trends in influenza vaccination uptake in Switzerland: Swiss Health Survey 2007 and 2012. Swiss Med. Wkly. 149, w14705, doi: 10.4414/smw.2019.14705 (2019).

25 Kimiya, T. et al. Effectiveness of inactivated quadrivalent influenza vaccine in the 2015/2016 season as assessed in both a test-negative case-control study design and a traditional case-control study design. Eur. J. Pediatr. 177, 1009–1017, doi: 10.1007/s00431-018-3145-7 (2018).

26 Puig-Barbera, J. et al. Low influenza vaccine effectiveness and the effect of previous vaccination in preventing admission with A(H1N1)pdm09 or B/Victoria-Lineage in patients 60 years old or older during the 2015/2016 influenza season. Vaccine 35, 7331–7338, doi: 10.1016/j.vaccine.2017.10.100 (2017).

27 Fine, P., Eames, K. & Heymann, D. L. “Herd immunity”: a rough guide. Clin. Infect. Dis. 52, 911–916, doi: 10.1093/cid/cir007 (2011).

28 Kim, T. H., Johnstone, J. & Loeb, M. Vaccine herd effect. Scand. J. Infect. Dis. 43, 683–689, doi: 10.3109/00365548.2011.582247 (2011).

29 Plant, E. P. et al. Different Repeat Annual Influenza Vaccinations Improve the Antibody Response to Drifted Influenza Strains. Sci. Rep. 7, 5258, doi: 10.1038/s41598-017-05579-4 (2017).

30 Olafsdottir, T. A. et al. Age and Influenza-Specific Pre-Vaccination Antibodies Strongly Affect Influenza Vaccine Responses in the Icelandic Population whereas Disease and Medication Have Small Effects. Front. Immunol. 8, 1872, doi: 10.3389/fimmu.2017.01872 (2017).

31 Trombetta, C. M., Perini, D., Mather, S., Temperton, N. & Montomoli, E. Overview of Serological Techniques for Influenza Vaccine Evaluation: Past, Present and Future. Vaccines (Basel) 2, 707–734, doi: 10.3390/vaccines2040707 (2014).

32 Egli, A. et al. Identification of influenza urban transmission patterns by geographical, epidemiological and whole genome sequencing: Protocol for an observational study. BMJopen, accepted, doi: http://dx.doi.org/10.1136/bmjopen-2019-030913 (2019).

33 Wuthrich, D. et al. Evaluation of two workflows for whole genome sequencing-based typing of influenza A viruses. J. Virol. Methods 266, 30–33, doi: 10.1016/j.jviromet.2019.01.009 (2019).

34 Kim, J. I. et al. Distinct molecular evolution of influenza H3N2 strains in the 2016/17 season and its implications for vaccine effectiveness. Mol. Phylogenet. Evol. 131, 29–34, doi: 10.1016/j.ympev.2018.10.042 (2019).

35 Xue, K. S., Moncla, L. H., Bedford, T. & Bloom, J. D. Within-Host Evolution of Human Influenza Virus. Trends Microbiol. 26, 781–793, doi: 10.1016/j.tim.2018.02.007 (2018).

36 Iuliano, A. D. et al. Estimates of global seasonal influenza-associated respiratory mortality: a modelling study. Lancet 391, 1285–1300, doi: 10.1016/S0140-6736(17)33293-2 (2018).

37 Organization, W. H. Global Influenza Strategy 2019-2030. 34 (WHO, 2019).

38 Chunara, R., Goldstein, E., Patterson-Lomba, O. & Brownstein, J. S. Estimating influenza attack rates in the United States using a participatory cohort. Sci. Rep. 5, 9540, doi: 10.1038/srep09540 (2015).

39 Lee, R. U., Phillips, C. J. & Faix, D. J. Seasonal Influenza Vaccine Impact on Pandemic H1N1 Vaccine Efficacy. Clin. Infect. Dis. 68, 1839–1846, doi: 10.1093/cid/ciy812 (2019).

40 Basel-Stadt, S. O. o. t. C. Statistischer Block, <https://www.statistik.bs.ch/zahlen/raumdaten/raumeinheiten/block.html> (2019).

41 Fitzner, J. et al. Revision of clinical case definitions: influenza-like illness and severe acute respiratory infection. Bull. World Health Organ. 96, 122–128, doi: 10.2471/BLT.17.194514 (2018).

42 Organization, W. H. WHO Global Epidemiological Surveillance Standards for Influenza. 84 (WHO, 2014).

43 Bolger, A. M., Lohse, M. & Usadel, B. Trimmomatic: a flexible trimmer for Illumina sequence data. Bioinformatics 30, 2114–2120, doi: 10.1093/bioinformatics/btu170 (2014).

44 Langmead, B. & Salzberg, S. L. Fast gapped-read alignment with Bowtie 2. Nature methods 9, 357–359, doi: 10.1038/nmeth.1923 (2012).

45 Li, H. et al. The Sequence Alignment/Map format and SAMtools. Bioinformatics 25, 2078–2079, doi: 10.1093/bioinformatics/btp352 (2009).

46 Wilm, A. et al. LoFreq: a sequence-quality aware, ultra-sensitive variant caller for uncovering cell-population heterogeneity from high-throughput sequencing datasets. Nucleic Acids Res. 40, 11189–11201, doi: 10.1093/nar/gks918 (2012).

47 Edgar, R. C. MUSCLE: multiple sequence alignment with high accuracy and high throughput. Nucleic Acids Res. 32, 1792–1797, doi: 10.1093/nar/gkh340 (2004).

48 Munir, M. Bioinformatics analysis of large-scale viral sequences: from construction of data sets to annotation of a phylogenetic tree. Virulence 4, 97–106, doi: 10.4161/viru.23161 (2013).

49 Stamatakis, A. RAxML version 8: a tool for phylogenetic analysis and post-analysis of large phylogenies. Bioinformatics 30, 1312–1313, doi: 10.1093/bioinformatics/btu033 (2014).

50 Klein, E. Y., Serohijos, A. W., Choi, J. M., Shakhnovich, E. I. & Pekosz, A. Influenza A H1N1 pandemic strain evolution--divergence and the potential for antigenic drift variants. PLoS One 9, e93632, doi: 10.1371/journal.pone.0093632 (2014).

51 Wickham, H. ggplot2: Elegant Graphics for Data Analysis (UseR!). 2nd edn, 276 (Springer, New York, 2016).

52 Letunic, I. & Bork, P. Interactive Tree Of Life (iTOL): an online tool for phylogenetic tree display and annotation. Bioinformatics 23, 127–128, doi: 10.1093/bioinformatics/btl529 (2007).

53 Krzywinski, M. et al. Circos: an information aesthetic for comparative genomics. Genome Res. 19, 1639–1645, doi: 10.1101/gr.092759.109 (2009).

54 Hadfield, J. et al. Nextstrain: real-time tracking of pathogen evolution. Bioinformatics 34, 4121–4123, doi: 10.1093/bioinformatics/bty407 (2018).

55 Elbe, S. & Buckland-Merrett, G. Data, disease and diplomacy: GISAID’s innovative contribution to global health. Global challenges 1, 33–46, doi: 10.1002/gch2.1018 (2017).

56 Katoh, K., Misawa, K., Kuma, K. & Miyata, T. MAFFT: a novel method for rapid multiple sequence alignment based on fast Fourier transform. Nucleic Acids Res. 30, 3059–3066, doi: 10.1093/nar/gkf436 (2002).

57 Nguyen, L. T., Schmidt, H. A., von Haeseler, A. & Minh, B. Q. IQ-TREE: a fast and effective stochastic algorithm for estimating maximum-likelihood phylogenies. Mol. Biol. Evol. 32, 268–274, doi: 10.1093/molbev/msu300 (2015).

58 Sagulenko, P., Puller, V. & Neher, R. A. TreeTime: Maximum-likelihood phylodynamic analysis. Virus evolution 4, vex042, doi: 10.1093/ve/vex042 (2018).

59 Zou, G. A modified poisson regression approach to prospective studies with binary data. Am J Epidemiol 159, 702–706, doi: 10.1093/aje/kwh090 (2004).

60 Shapiro, B., Rambaut, A. & Drummond, A. J. Choosing appropriate substitution models for the phylogenetic analysis of protein-coding sequences. Mol. Biol. Evol. 23, 7–9, doi: 10.1093/molbev/msj021 (2006).

61 Bouckaert, R. et al. BEAST 2.5: An advanced software platform for Bayesian evolutionary analysis. PLoS Comput. Biol. 15, e1006650, doi: 10.1371/journal.pcbi.1006650 (2019).

62 Vaughan, T. G., Kuhnert, D., Popinga, A., Welch, D. & Drummond, A. J. Efficient Bayesian inference under the structured coalescent. Bioinformatics 30, 2272–2279, doi: 10.1093/bioinformatics/btu201 (2014).

