## Supplementary files 2 and 3 for "High-resolution influenza mapping of a city reveals socioeconomic determinants of transmission within and between urban quarters"

Universität Basel | Departement Umweltwissenschaften – Humangeographie Stadt- und Regionalforschung (DUW) | Universitätsspital Basel | Departement Biomedizin Basel | ETH Zürich

### Questionnaire on Influenza in the City of Basel

Within the scope of the **research project on influenza in the city of Basel**, we are conducting a **survey in selected quarters of Basel**.

The participation in the survey takes approximately 15 minutes and is voluntary. **Your anonymity will be ensured**. We ask you to return this brochure with the completed questionnaire until 1st May 2016 with the postage-paid envelope.

Participants can partake in a lottery (**Apple Laptop worth CHF 2500.- and two "Pro Innerstadt"-coupons each worth CHF 500.-).** See end of questionnaire. Many thanks for your participation.

#### A. Influenza and common cold

**A.1. Did you suffer from a strong cold this winter?**

☐1 no *(if no, please continue with question A6)* ☐2 yes, multiple times ☐3 yes, once

If **yes**, during which month(s): ____________________

If **yes**, how many days did it considerably impact your day-to-day life? __________

**A.2. Which grievances did you have?** (*multiple answers possible*)

☐1 temperature (over 38 degrees) ☐3 muscle and rheumatic pains ☐5 cough ☐7 sore throat ☐9 runny nose

☐2 diarrhoea ☐4 strong fatigue ☐6 headache ☐8 strong sense of illness

**A.3. Has anyone else in your proximity been suffering from the same grievances (influenza/cold) and fallen ill?**

☐1 no ☐2 yes ☐3 don´t know

If **yes,** people (*multiple answers possible*):

☐1 in the family ☐2 at work ☐3 in the neighbourhood ☐4 in the circle of friends ☐5 in the "Verein" (club)

**A.4. Did you contact an expert regarding the cold symptoms?**

☐1 no ☐2 yes

If **yes**, where?

☐1 medical practice ☐2 hospital ☐3 pharmacy ☐4 other, namely: ____________________

**A.5. Did you take medication to cure of the cold?**

☐1 no ☐2 yes

If **yes,** did you receive Tamiflu?

☐1 yes ☐2 no

**A.6. Did you get vaccinated in autumn/winter 2015/16 against Influenza?**

☐1 no ☐2 yes

If **yes**, where: ☐1 medical practice ☐2 pharmacy ☐3 hospital ☐4 other, namely: ______

If yes, during which month? ____________________

If **no**: Why did you not get a vaccination? (please mark all relevant points)

☐1 I don´t know why I should get vaccinated.
☐2 I wanted to, but then I ended up not doing it.
☐3 I don´t believe in the effect of the vaccination.
☐4 A real flu strengthens my immune system and the protection lasts longer.
☐5 It´s too expensive for me.
☐6 I strengthen my immune system with other means.
☐7 Friends/family have had negative experiences with it.
☐8 I´m afraid of the side effects.
☐9 I find needles unpleasant.
☐10 The vaccination was not recommended to me by experts (doctor/pharmacist).
☐11 other reason, namely: ____________________

If **yes**: Why did you get vaccinated? (please mark all relevant points)

☐1 I don´t want to get influenza.
☐2 I don´t want to be missing at work.
☐3 I have friends/family who I want to protect from influenza.
☐4 The vaccination was recommended to me by family/friends.
☐5 The vaccination was recommended to me at work.
☐6 The vaccination was recommended to me by experts (doctor/pharmacist) .
☐7 other reason, namely: ____________________

**A.7. If you got vaccinated against influenza this year, state your experiences on a scale of 1 (very negative) to 10 (very positive):**

☐1 1 ☐2 2 ☐3 3 ☐4 4 ☐5 5 ☐6 6 ☐7 7 ☐8 8 ☐9 9 ☐10 10

very negative — very positive

If **positive** experiences, which: ______________________________

If **negative** experiences, which: ______________________________

**A.8. Do you get vaccinated against influenza on a regular basis?**

☐1 no ☐2 yes

If **yes**, since how many years: __________

**A.9. In the past, have you refused other vaccinations which were recommended to you or your children by a doctor?**

☐1 no, never ☐2 yes, but only certain vaccinations ☐3 yes, all vaccinations

**A.10. How do you evaluate your general state of health on a scale from 1 (very negative) to 10 (very positive)?**

☐1 1 ☐2 2 ☐3 3 ☐4 4 ☐5 5 ☐6 6 ☐7 7 ☐8 8 ☐9 9 ☐10 10

very negative — very positive

**A.11. Do you have a general practitioner?**

☐1 no ☐2 yes

**A.12. Do you have any chronic diseases due to which you have to take drugs on a daily basis (since at least 6 months)?**

☐1 no ☐2 yes

If **yes,** which drugs: ______________________________

**A.13. Do you smoke?**

☐1 no ☐2 yes, sometimes ☐3 yes, daily (number of packets per day): _________ ☐4 not specified

**A.14. Do you drink alcohol?**

☐1 no ☐2 yes, sometimes ☐3 yes, daily ☐4 not specified

**A.15. How important do you regard the following measures to prevent influenza?**

| | important | rather important | neutral | rather unimportant | unimportant | don´t know | no specification |
|---|---|---|---|---|---|---|---|
| Vaccinating | ☐1 | ☐2 | ☐3 | ☐4 | ☐5 | ☐6 | ☐7 |
| Washing hands | ☐1 | ☐2 | ☐3 | ☐4 | ☐5 | ☐6 | ☐7 |
| Health check-up at the doctor | ☐1 | ☐2 | ☐3 | ☐4 | ☐5 | ☐6 | ☐7 |
| Healthy diet | ☐1 | ☐2 | ☐3 | ☐4 | ☐5 | ☐6 | ☐7 |
| Regular physical activity | ☐1 | ☐2 | ☐3 | ☐4 | ☐5 | ☐6 | ☐7 |
| Other | ☐1 | ☐2 | ☐3 | ☐4 | ☐5 | ☐6 | ☐7 |

other, namely: ______________________________

**A.16. How good do you implement the following measures into your day-to-day life?**

| | very good | good | average | bad | very bad | don´t know | no specification |
|---|---|---|---|---|---|---|---|
| Vaccinating | ☐1 | ☐2 | ☐3 | ☐4 | ☐5 | ☐6 | ☐7 |
| Washing hands | ☐1 | ☐2 | ☐3 | ☐4 | ☐5 | ☐6 | ☐7 |
| Health check-up at the doctor | ☐1 | ☐2 | ☐3 | ☐4 | ☐5 | ☐6 | ☐7 |
| Healthy diet | ☐1 | ☐2 | ☐3 | ☐4 | ☐5 | ☐6 | ☐7 |
| Regular physical activity | ☐1 | ☐2 | ☐3 | ☐4 | ☐5 | ☐6 | ☐7 |
| Other | ☐1 | ☐2 | ☐3 | ☐4 | ☐5 | ☐6 | ☐7 |

#### B. Aspects of city environment

##### B.1. How often do you use the following means of transport?

| | daily | several times per week | several times per month | rarely | never | don´t know | no specification |
|---|---|---|---|---|---|---|---|
| Car, motor cycle, motor scooter | ☐1 | ☐2 | ☐3 | ☐4 | ☐5 | ☐6 | ☐7 |
| Public transport (bus, tram, train) | ☐1 | ☐2 | ☐3 | ☐4 | ☐5 | ☐6 | ☐7 |
| Bike | ☐1 | ☐2 | ☐3 | ☐4 | ☐5 | ☐6 | ☐7 |
| On foot | ☐1 | ☐2 | ☐3 | ☐4 | ☐5 | ☐6 | ☐7 |
| Other | ☐1 | ☐2 | ☐3 | ☐4 | ☐5 | ☐6 | ☐7 |

other, namely: ________________________________________________

##### B.2. How often do you undertake in the following activities?

| | daily | several times per week | several times per month | rarely | never | don´t know | no specification |
|---|---|---|---|---|---|---|---|
| Shopping in supermarkets (Coop, Migros, etc.) | ☐1 | ☐2 | ☐3 | ☐4 | ☐5 | ☐6 | ☐7 |
| Cinema | ☐1 | ☐2 | ☐3 | ☐4 | ☐5 | ☐6 | ☐7 |
| Restaurant/ café/ bar | ☐1 | ☐2 | ☐3 | ☐4 | ☐5 | ☐6 | ☐7 |
| Cultural events | ☐1 | ☐2 | ☐3 | ☐4 | ☐5 | ☐6 | ☐7 |
| Sporting events / games | ☐1 | ☐2 | ☐3 | ☐4 | ☐5 | ☐6 | ☐7 |

**B.3. Approximately, with how many people do you have contact on a regular work day (work environment, family, friends, clubs)?**
☐1 0-10 ☐2 10-50 ☐3 50-100 ☐4 more than 100

**B.4. If you are working, do you have contact with other people?**
☐1 no ☐2 yes ☐3 no specification

**B.5. Do you work in the health sector with contact to patients (medical personnel or nurse)?**
☐1 no ☐2 yes ☐3 no specification

**B.6. Do you work with children (kindergarten, play group, day care, school etc.)?**
☐1 no ☐2 yes ☐3 no specification

**B.7. Is your workplace in an open-plan office or in a room with many people?**
☐1 no ☐2 yes ☐3 no specification

**B.8. Do you feel exposed to damaging environmental influences in your living environment? (e.g. emissions, fine dust, electric smog, noise etc.)**
☐1 no ☐2 yes ☐3 no specification
If **yes**, which:___________________

**B.9. Have you visited people in hospital or an old people´s home this autumn/winter 2015/16?**
☐1 no ☐2 yes
If **yes**, how often:
☐1 1-4 times ☐2 more than 5 times ☐3 I regularly visit people in healthcare facilities.

**B.10. Do you regularly look after care-dependent family members at home?**
☐1 no ☐2 yes

**B.11. Do you live with people who suffer from a chronic disease?**
☐1 no ☐2 yes

### C. Procurement of Information on Health Questions

**C.1. From where do you procure your information on health questions?** (*multiple answers possible*)

☐1 doctor
☐2 pharmacy
☐3 social circle (friends, acquaintances, family etc.)
☐4 religio-cultural context
☐5 other source of information, namely:____________________
☐6 TV
☐7 radio
☐8 newspaper and magazines
☐9 social networks
☐10 internet, namely:
  ☐11 from experience reports of other people
  ☐12 from official websites (Federal Office of Public Health, hospital)
  ☐13 from other sites: ____________________

☐14 I don´t inform myself.

**C.2. How helpful do you find the information offered?**

| | very helpful | helpful | average | less helpful | not at all helpful | don´t know | no specification |
|---|---|---|---|---|---|---|---|
| **Doctor** | ☐1 | ☐2 | ☐3 | ☐4 | ☐5 | ☐6 | ☐7 |
| **Pharmacy** | ☐1 | ☐2 | ☐3 | ☐4 | ☐5 | ☐6 | ☐7 |
| **Social circle (friends, acquaintances, family etc.)** | ☐1 | ☐2 | ☐3 | ☐4 | ☐5 | ☐6 | ☐7 |
| **Religio-cultural context** | ☐1 | ☐2 | ☐3 | ☐4 | ☐5 | ☐6 | ☐7 |
| **TV** | ☐1 | ☐2 | ☐3 | ☐4 | ☐5 | ☐6 | ☐7 |
| **Radio** | ☐1 | ☐2 | ☐3 | ☐4 | ☐5 | ☐6 | ☐7 |
| **Newspaper and magazines** | ☐1 | ☐2 | ☐3 | ☐4 | ☐5 | ☐6 | ☐7 |
| **Social networks** | ☐1 | ☐2 | ☐3 | ☐4 | ☐5 | ☐6 | ☐7 |
| **Experience reports of other people on the internet** | ☐1 | ☐2 | ☐3 | ☐4 | ☐5 | ☐6 | ☐7 |
| **Official websites on the internet** | ☐1 | ☐2 | ☐3 | ☐4 | ☐5 | ☐6 | ☐7 |
| **Other websites on the internet** | ☐1 | ☐2 | ☐3 | ☐4 | ☐5 | ☐6 | ☐7 |
| **Other sources of information** | ☐1 | ☐2 | ☐3 | ☐4 | ☐5 | ☐6 | ☐7 |

#### D. Personal Data

**D.1. Gender**

☐1 male ☐2 female

**D.2. Year of birth:**

______________________________

**D.3. Nationality**

☐1 Swiss ☐2 other nationality: ________________________________ ☐3 no specification

**D.4. How long have you been living at your current location?**

☐1 up to 1 year ☐3 more than 2 to 5 years ☐5 more than 10 to 15 years ☐7 no specification
☐2 more than 1 to 2 years ☐4 more than 5 to 10 years ☐6 more than 15 years

**D.5. Please look at the map of your quarter on the last page. In which segment do you live?** Please state the coordinates (a combination of a letter and a number on the horizontal respectively vertical axis).
Segment:____________________

**D.6. Residential status of the household you are living in:**

☐1 owner-occupied flat ☐3 rental apartment ☐5 co-operative flat ☐7 other:_______________
☐2 owner-occupied house ☐4 rental house ☐6 old people´s home ☐8 no specification
Number of square metres of the apartment/house: _________________

**D.7. How many people live in your household / flat share including yourself?**

_______ people

**D.8. How many children live in your household?**

| | 0 | 1 | 2 | 3 | More than 3 |
|---|---|---|---|---|---|
| Children under 7 years old | ☐1 | ☐2 | ☐3 | ☐4 | ☐5 |
| Children over 7 years old | ☐1 | ☐2 | ☐3 | ☐4 | ☐5 |

**D.9. If you have children under 7 years old, are they being looked after externally together with other children?**

☐1 no ☐2 yes

**D.10. What is the structure of your household?**

☐1 one-person household ☐3 (married) couple with child ☐5 single parent with child ☐7 no specification
☐2 flat share ☐4 (married) couple without child ☐6 other:___________

**D.11. Where have you attained your educational qualification?**

☐1 compulsory school
☐2 vocational education / -training, trade school
☐3 gymnasium
☐4 higher vocational education (commercial college, higher professional school, foreman, technician)
☐5 institution of higher education (ETH, university, college, teacher training college)
☐6 other school qualification _____________
☐7 no specification

**D.12. Are you currently employed? (*multiple answers possible*)**

☐1 full-time (min. 90%) ☐3 pupil/apprentice/student ☐5 pensioner ☐7 voluntary work
☐2 part-time/side job (<90%) ☐4 housewife/househusband ☐6 currently not employed ☐8 no specification

**D.13. If you are employed, what position do you hold?**

☐1 employee with management function ☐2 employee without management function ☐3 self-employed

**D.14. Category of income (monthly gross household income)**

☐1 up to CHF 2000 ☐3 4001-6000 CHF ☐5 8001-10‘000 CHF ☐7 > 15‘000 CHF
☐2 2001-4000 CHF ☐4 6001-8000 CHF ☐6 10‘001-15‘000 CHF ☐8 no specification

**D.15. Where do you work? (place and postcode):** ____________________________________________

**Map for question D5**
Please look at the map of your quarter and mark the approximate location of your residence. Please state the coordinates (a combination of a letter and a number on the horizontal respectively vertical axis).

**Example (right)**
You live at the corner of Bläsiring / Müllheimerstrasse.
Answer: N12

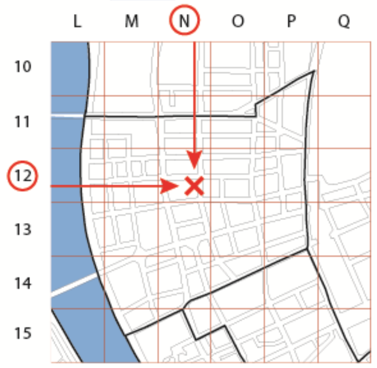


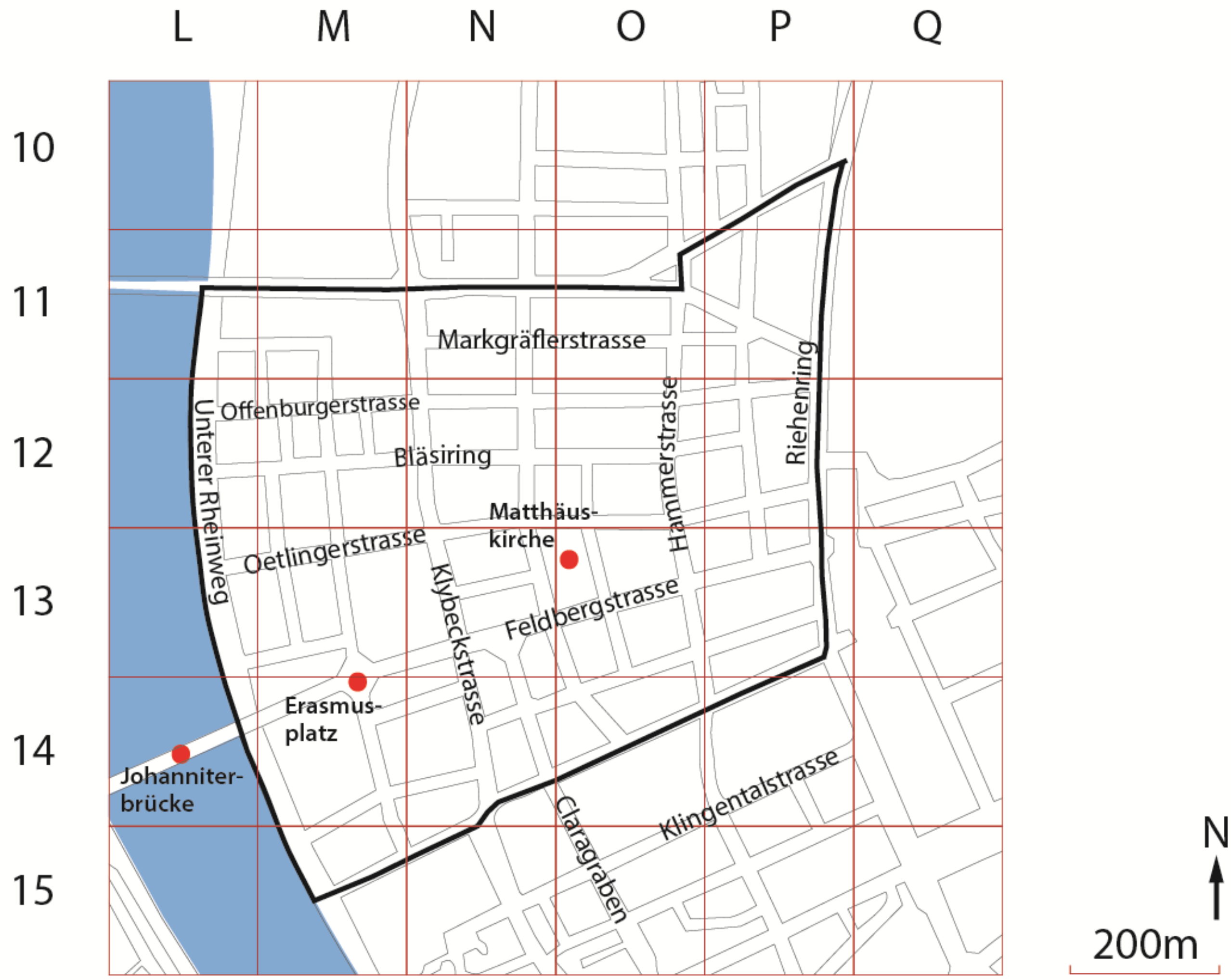


#### Thank you for your participation.

**If you would like to participate in the lottery for an Apple computer worth CHF 2500.- and two "Pro Innerstadt"-coupons each worth CHF 500.-, you can leave your name and address here:**

MA

### Characterising the epidemic spread of Influenza A/H3N2 within a city through phylogenetics

Nicola F. Müller[a,b,+], Daniel Wüthrich[b,c,d], Nina Goldman[e], Nadine Sailer[e], Claudia Saalfrank[e], Myrta Brunner[e], Noémi Augustin[e], Helena MB Seth-Smith[b,c,d], Yvonne Hollenstein[c,d], Mohameedyaseen Syedbasha[c,d], Daniela Lang[c,d], Richard A. Neher[b,f], Olivier Dubuis[g], Michael Naegele[g], Andreas Buser[h], Christian H. Nickel[i], Nicole Ritz[j], Andreas Zeller[k], Brian M. Lang[a,b], James Hadfield[l], Trevor Bedford[l], Manuel Battegay[m], Rita Schneider-Sliwa[e], Adrian Egli[c,d,*] and, Tanja Stadler[a,b,*,+].

[a] ETH Zürich, Department of Biosystems Science and Engineering, 4058 Basel, Switzerland
[b] Swiss Institute of Bioinformatics (SIB), Lausanne, Switzerland
[c] Clinical Bacteriology and Mycology, University Hospital Basel, Basel, Switzerland
[d] Applied Microbiology Research, Department of Biomedicine, University of Basel, Basel, Switzerland
[e] Human Geography, Department of Environmental Sciences, University of Basel
[f] Biozentrum, University of Basel, Basel, Switzerland
[g] Viollier AG, Allschwil, Switzerland
[h] Blood donation center, Swiss red cross, Basel, Switzerland
[i] Emergency Department, University Hospital Basel, Basel, Switzerland
[j] Pediatric Infectious Diseases and Vaccinology, University Children's Hospital Basel and University of Basel, Basel Switzerland
[k] Institute for Family Medicine, University of Basel, Basel, Switzerland
[l] Fred Hutchinson Cancer Research Center, Seattle, USA
[m] Infectious Diseases and Hospital Epidemiology, University Hospital Basel, Basel, Switzerland

[*]Equal contribution
[+]Corresponding authors

**Abstract** Infecting large portions of the global population, seasonal influenza is a major burden on societies around the globe. In order to improve our understanding of how influenza is transmitted on a city scale, we collected an extremely densely sampled set of influenza sequences alongside patient metadata. We find that repeated introductions into the city drove the 2016/2017 influenza season and that downstream transmission dynamics correlated positively with temperature. Zooming into the local transmission outbreaks suggests that the elderly were to a large extent infected within their own transmission network, while school children drove the spread within the remaining transmission network. These patterns will be valuable to plan interventions combating the spread of influenza within cities given that similar patterns are observed for other influenza seasons and cities.

### Main

With a large fraction of the population being infected annually and up to 650,000 deaths per year, seasonal influenza causes a major burden on societies around the globe (https://www.who.int/news-room/fact-sheets/detail/influenza-(seasonal)). Through rapid evolution, influenza strains evade host immunity, allowing them to reinfect large fractions of a population every year. In order to prevent infections, limited public health resources have to be streamlined as efficiently as possible [1]. The planning of interventions is dependent upon knowledge of the dynamics of epidemic spread of influenza viruses in a city environment, which includes understanding the drivers of the spread of seasonal influenza between individuals. Incidence and prevalence data can be used to some extent to infer such dynamics. However, they lack the information about how individual cases are epidemiologically related.

Phylogenetics allows us to see how individual cases are epidemiologically connected. This is done by reconstructing the evolutionary relationship between temporally spaced samples of genetic sequence data, isolated from different infected individuals. The resulting phylogenetic tree displays how samples are related to each other, and branch lengths in calendar time display the elapsed time. The phylogenetic tree can therefore be interpreted as an approximation of the transmission chain of the sampled cases. Such a view on part of the influenza transmission chain allows to further quantify the epidemiological dynamics which gave rise to the observed phylogenetic tree using phylodynamic methods [2]. Phylogenetics and phylodynamics thus allows us to elucidate past epidemiological dynamics [3, 4] or to infer migration patterns [5, 6].

Several studies have used phylogenetic approaches to study how influenza and its subtypes spread globally [7–11]. On an intermediate scale, college campuses have been studied by using phylogenetics, revealing extensive mixing of influenza strains [12]. On the smallest scale, studies have been performed to investigate person-to-person transmission of influenza in households [13]. There is however a gap in studies that seek to describe transmission of influenza on a city scale. In contrast to college campuses, cities constitute highly heterogeneous societies with various different living arrangements and vastly different social and age groups. This means that lessons learnt about influenza transmission from college campuses are not necessarily transferable to cities. Children have, for example, been described to make up a larger portion of the infected population at the beginning of a season compared to the end over 3 seasons in Houston [14].

In an effort to fill that gap, we studied the local spread of influenza and the factors contributing to it in the city of Basel, Switzerland during the 2016/2017 Influenza season which was dominated by Influenza A/H3N2. To do so, influenza samples together with the age and residential address were collected from around 669 patients from the University Hospital (USB), the Children's Hospital of Basel (UKBB) and patients of private practices from around the city. Around 200 of these patients also provided additional information through filling out a survey. The survey asked questions about family status, financial status, and demographics. Details on the data collection is provided in [15]. We assess here the importance of introductions of influenza into a city for seeding a seasonal epidemic, the overall dynamics of transmission throughout the season, and the extent to which different age groups and family statuses drive the spread of seasonal influenza in a city.

#### Introduction of new lineages into the city drive the local epidemic

We first assessed how the 669 sampled cases in Basel compare to 11,000 sequences sampled from around the world between January 2016 and December 2017, by inferring a phylogenetic tree using the hemagglutinin (HA) sequences. The Basel sequences span the existing global diversity (see figure 1a), suggesting strong exchange of viruses with other areas around the globe. We however did not find isolates in Basel that were part of the 3c2 and 3c3 clades; notably a strain of the 3c2 clade was the vaccine strain in that year. The number of sampled sequences in Basel peaked at the end of 2016, with a smaller peak at the beginning of February of 2017 (see figure 1b). The sequenced cases were dominated by the 3c2 sub-clades A1, A1a and A3, and we observe that a peak in A1a cases mainly contributed to the peak in February of 2017.

Basel sequences cluster into local transmission clusters within the global diversity (see Materials and Methods). We obtained around 240 local clusters (see figure 1c and https://nextstrain.org/community/jameshadfield/basel-flu/1), suggesting that the sampled sequences were the result of around 240 influenza introductions from areas outside of Basel. In order to investigate if this number is a strict lower bound for the number of introductions, we use random subsets of the 669 sequences to re-estimate the number of introductions. We find that the number of estimated introductions grows approximately linearly with the number of sequences in a subset (see figure 1c). This suggests that with additional sampling effort, we would have captured more introductions. With its own international airport, the airport of Zürich nearby, and a major rail hub, Basel is well connected to the rest of Europe and the world. As such, people working in Basel often do not live in the city and commute daily from elsewhere in Switzerland, Germany and France. Basel is a tourist destination and often hosts international conferences, attracting people from all over the world. This connectedness likely drives these introductions of influenza into the city.

#### Quantification of the overall local epidemic following the introduction into the city

After introduction into the city, we next study how influenza is transmitted locally. Clusters with more than one isolate have an average tree height of around 30 days (see figure 1e), which we interpret as a lower bound of how long transmission chains persisted within the city. In order to quantify the amount of local transmission, we estimated the effective reproduction number ($R_{eff}$) to be between 1 and 1.5 for most of the season, which agrees with previous estimates of the effective reproduction number for seasonal influenza [16]. The $R_{eff}$ peaks in December and in February (see figure 1f) with the 95% credible interval excluding and $R_{eff}$ of 1 in December. In January, we inferred a drop in the effective reproduction number with the estimated median being below 1, which is consistent with the trend in the number of sampled cases (see figure 1f). Further, these estimates are comparable to the overall trend of influenza cases during the 2016/2017 season in Europe (https://ecdc.europa.eu/en/publications-data/summary-influenza-2016-2017-season-europe).

We next investigated potential factors determining the changes in $R_{eff}$. The number of influenza cases over the years show a strong seasonality, with the majority of cases occurring in the winter months in both the northern and southern hemisphere [7]. Relative humidity and temperature have been described to drive influenza transmission [17]. Additionally, the effect of school closures on the spread of pandemic influenza has been discussed [18, 19]. Thus we investigated potential correlations of $R_{eff}$ with temperature,relative humidity and school days (i.e. days when children go to school). As we only studied one season, these correlations have to be interpreted with caution, and analyses of other seasons are needed to confirm the potential correlations. Neither humidity nor school days showed a significant correlation: relative humidity stayed fairly constant over the season, and both low and high $R_{eff}$ are found during times when schools were open (figure 1g and figures S7, S8 and S9). To account for autocorrelation, we performed the correlation analysis, averaging the temperature, relative humidity and mean number of school days over 4, 6, 8 and 10 days instead of just 2 days. We find the mean temperature to be significantly correlated with the mean Ref in both scenarios (see figures S6 and S8). While previous studies in animal models observed higher effective reproduction numbers at lower temperatures,

Viral shedding of viruses has been shown previously to be increased at lower temperatures in animal models [17] and higher absolute humidity has been shown to favour transmission on a population level23. We here observe lower effective reproduction numbers at lower temperatures [20]. The correlation of the effective reproduction number with the temperature however is not necessarily causal, as it for example could be due to social behaviour being different at lower temperatures. Additionally, the computed p-values could be inflated due to unaccounted autocorrelation.

Along with the $R_{eff}$, we co-estimated the sampling probability, that is the probability of an infected individual being sampled. Since we followed the same procedure for inclusion of patients throughout the epidemic season [15], we assumed that this probability is constant throughout the influenza season. We estimated the sampling probability to be between 3% and 5% (see figures 1d and S5). In contrast to the $R_{eff}$ estimates, this value is more sensitive to the procedure of clustering of sequences into sets of locally transmitted sequences (see figure S5). Additionally, the prior probability on the effective reproduction number, as well as the assumed becoming un-infectious rate can influence this estimate [21]. With 676 samples from different patients included in this analysis, this would suggest that between 13'500 and 22'500 people in Basel or between 8% and 13% of its population of about 171'000 were infected with influenza H3N2 during the 2016-17 season. The city limits of Basel are however in reality not fixed and the metropolitan area of Basel is much larger. Furthermore we have sampled patients who went to a doctor or Hospital in the city of Basel, but live in the surrounding areas or other parts of the world. We therefore expect the estimate of between 8% and 13% to be an estimate for the upper bound of the number of infected people, rather than the true percentage of infected individuals.

Overall, our analyses suggest that transmission occurred with an effective reproduction number varying between 1-1.5 throughout the season, overall infecting at most 8-13% of the population.

#### Importance of age groups and family status in the local epidemic spread

After having determined the importance of introductions of influenza into the city and the overall rate at which influenza is spread in the city, we next studied the effect that age and family status has on the overall spread. Figures 2a and b show the patient age distribution within our samples. In order to study the role of age and family status in spreading influenza, we next subdivided our Basel patients into four different age groups, preschoolers (<7 years old), school-aged children (7 to 17 years), adults (18 to 65 years) and the elderly (>65 years old). We further categorized adults into three subgroups corresponding to family status: adults for whom we know that they live in the same household as children, adults for whom we know that they do not and adults for whom we do not have this information. We thus have overall six different categories of patient groups.

For each individual infected with influenza in each of these categorical groups, we determined the number of patients with influenza viruses isolated below a certain phylogenetic distance. This number, we then define as the number of connections a patient has. A connection exists if the pairwise phylogenetic distance between viruses isolated from two patients is at most 0.1 years. We then evaluate the mean number of connections of a negative binomial distribution for all individuals from each of the six groups. We later repeated the analyses using different cutoff values. his way of defining two viral isolates to be connected is therefore done independently of the above used definition of local transmission clusters. Using pairwise distances allows us to use distance in the transmission chain whereas from the same cluster only says if two sequences originated from the same introduction.

We find that school-aged children are on average connected to more individuals, than preschoolers (see figure 2c). This difference is statistically significant after multiple hypothesis testing at a cutoff value of 0.1 years, but not other cutoff values. We further, but not significantly, find that adults that reported to live in the same household as children are on average connected to more patients than those that do not live in the same household as children. For the elderly, we find that they have significantly more connections compared to adults with unknown household status, adults without children, and preschoolers. They do not have more connections on average compared to adults living in the same household as children and school aged children. In summary, there is signal for school children and elderly having more connections to other individuals compared to the three groups unknown household status, adults without children, and preschoolers. The adult living in the same household as children group show tendencies to be connected more in average than the latter three groups, though the data is not informative enough, respectively we do not have enough data, to provide strong evidence for that.

That school aged children and elderly are connected to more individuals than the other groups can have different explanations. The most obvious one is that individuals of these groups are more likely to participate in transmission events, either as a donor or recipient; alternatively, strong mixing within a group and a higher probability of visiting a doctor upon infection and therefore a higher sampling probability could act as an explanation (if members from any group are equally likely to transmit or receive influenza to and from members of any other group, higher sampling probability would increase to number of connections of every group and not just one).

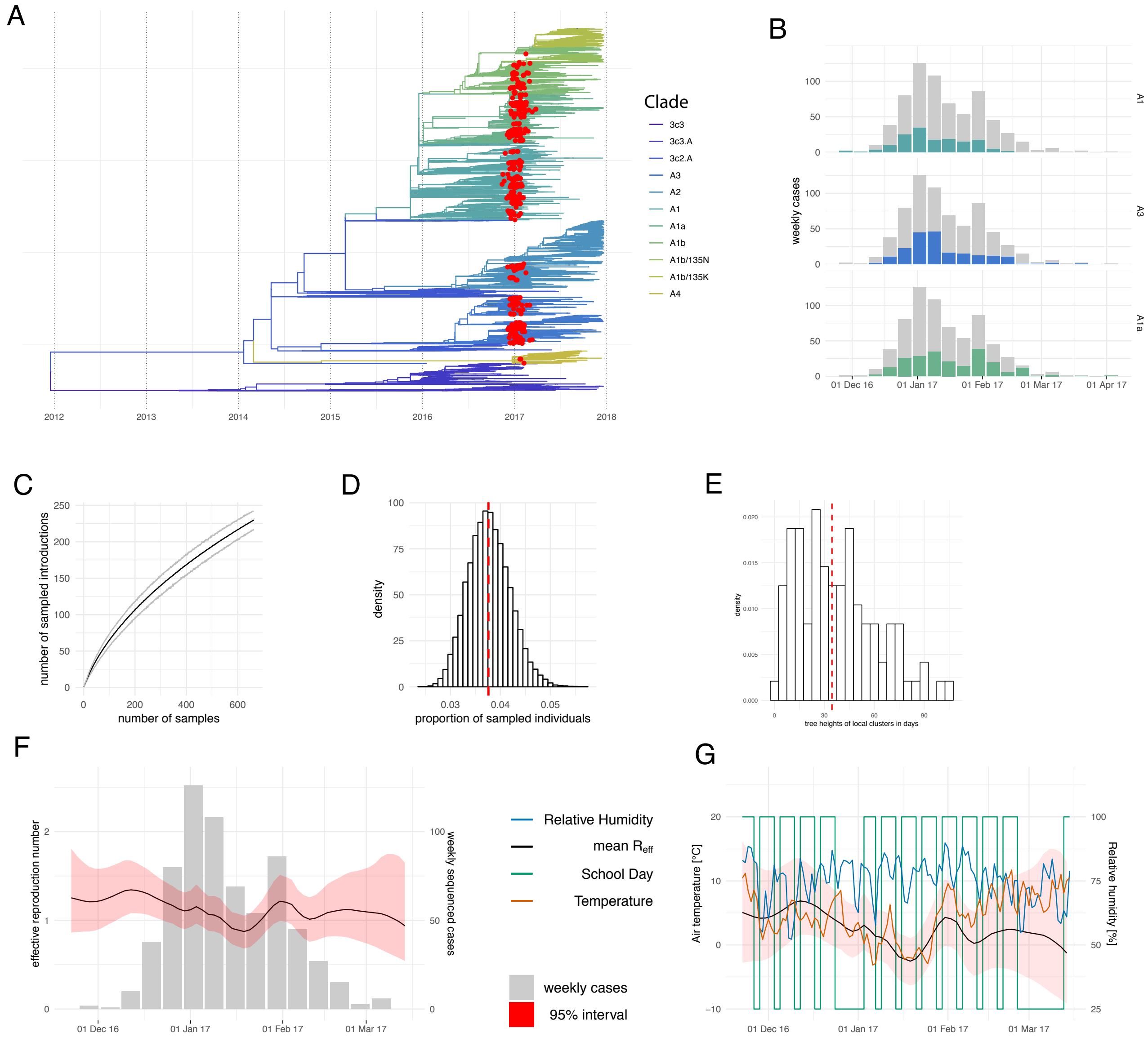


Figure 1: **The local spread of Influenza H3N2 in Basel in the 2016/2017 season. A** Time tree of HA segment reconstructed from sequences from this study and sequences from around the world. The HA sequences sampled in this study (red) are dispersed across almost all clades present between 2016-2018. This indicates that the diversity of samples in Basel is similar to the diversity of samples around the globe. **B** Number of sequenced weekly cases for the three most abundant HA clades. The number of cases for the corresponding clade is shown in color and the overall number of cases is shown in grey. **C** Number of introductions depending on how many random sequences from Basel are used. **D** Estimated proportion of sampled individuals averaged over ten different bdsky runs using different classifications of local sequences into local clusters. Estimates for individual classifications are shown in figure S4. **E** Estimates of tree heights of local clusters, which can be used as a lower bound to how long lineages persisted in the city. **F** Estimates of the effective reproduction number through time inferred from all local cluster jointly by using BDSKY. The black line is the mean estimate and the red area denotes the 95% interval. The grey bars denote the number of cases per week that were sequenced and used in the BDSKY analysis. **G** Comparison of estimates of the effective reproduction number with temperature and relative humidity and if a day is a school day or not. The effective reproduction number curves are averaged over the 10 different classifications of sequences into local clusters. Comparisons for the individual random classifications are shown in figure S5.

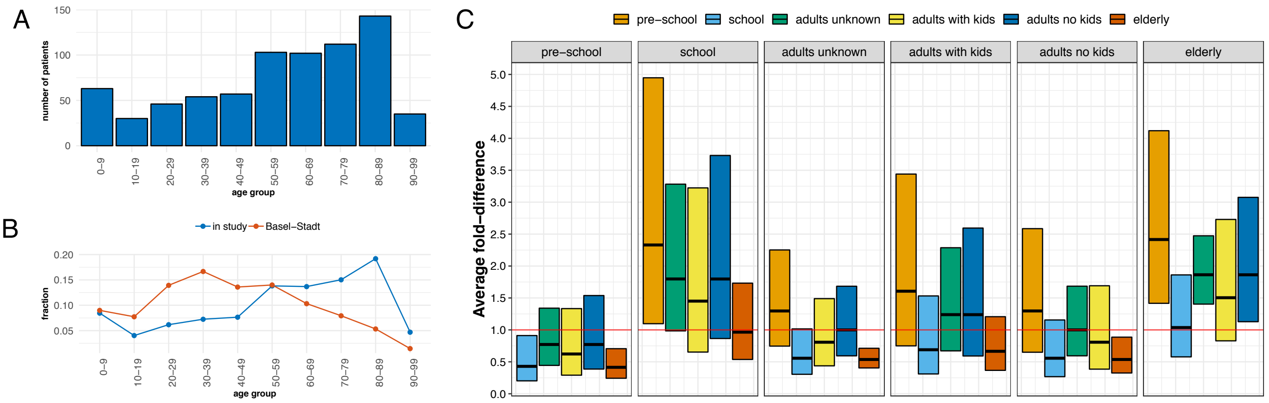


Figure 2: **Age Distribution and differences in the number of connections to other patients that members of the different categorical patient groups have. A** Distribution of patient ages in this study in age group intervals of 10 years. The intervals are different from the age groups used in the analysis due to the availability of age data for the population of Basel. **B** Proportion of different age groups in this study compared to in the city of Basel. **C** Model-based confidence intervals for the difference in average number of connections between each pair of age groups from a negative binomial model. Upper and lower bounds represent 95% confidence intervals for the average fold-difference in connection number between two groups corrected for multiple hypothesis testing using the Tukey method. This means that confidence intervals that do not include 1 are statistically significant. We see that the average connection number for elderly patients is twice the average connection number for preschoolers, adults without children, and adults with unknown status. These values are estimated for a cutoff values of 0.1 years. Estimates for different cutoff values are shown in figure S10. Also, the estimates of the mean number of connections in each group are shown in figure S11.

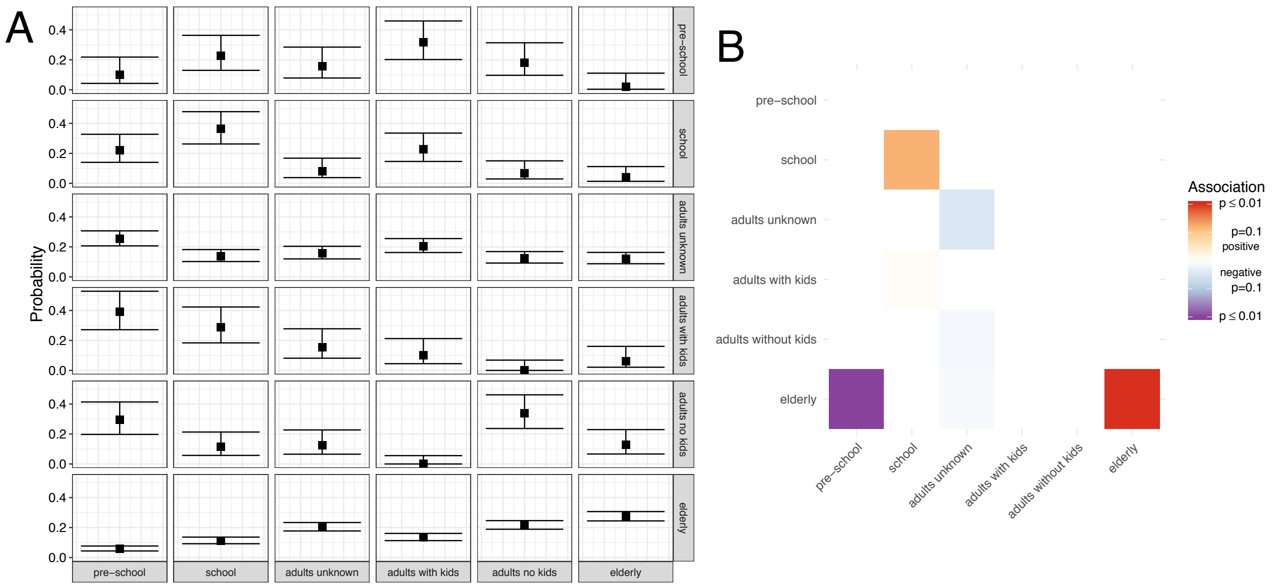


Figure 3: **Mixing of different age groups that are in proximity to each other.** Here we show the mixing patterns between the different categorial patient groups. We define pairs of patients to be connected if their pairwise phylogenetic distance was below 0.1 years. Results for other thresholds are shown in Figures S12 and S13. **A** Probability that an individual from the group in each row is connected to an individual from the group in a column. These probabilities were calculated by using the inverse number of samples from each group as weights. Upper and lower bounds correspond to 95% confidence intervals around the estimated probability. **B** The color of each tile in the heatmap corresponds to the p-value for either positive (red) or negative (blue) associations. These p-values are bonferroni corrected for the number of comparisons (42). We estimate these p-values by randomly permuting the group to patient labels and then comparing the number of pairs of interactions we observe in the data vs. when randomly permuting.

In order to assess the potential reasons, we investigated how strongly or weakly different age groups are connected with each other. We did so using two different approaches. First, we estimated the probability that an individual from a group is connected to a member from the same or a different group by using multinomial logistic regression with the inverse number of samples from each group as weights. Second, we use permutation testing to estimate the probability that the number of connections between different groups is significantly higher or lower than expected if all groups would be equally connected. To do so, we again use the definition of a connection between pairs of patients from the last section.

For both approaches, we find that mostly school-aged children are associated with other school-aged children, and the elderly are associated with other elderly people. We additionally find higher association between children of any age and adults living in the same household as children than children of any age and adults without children (see figure 3). Adults living in the same household as children, on the other hand, are estimated to have low association to other adults without children and much higher association to children, whereas adults without children are mostly associated to other adults without children (see figure 3a).

Increased sampling of the elderly relative to the other age groups is likely to occur, since the elderly are more likely to visit a doctor when in case of infections with influenza [22]. Thus, strong mixing within group and high sampling, might explain the increased connectivity of the elderly.

The second group that we found to have many connections to other patients where school-aged children. When looking to which groups these were connected, we found them to be associated with other school-aged children. However, they are unlikely to be sampled more than preschoolers [23], and we do not see evidence for oversampling of this group compared to preschoolers (see figure 2b). We therefore interpret our results as school-aged children being involved in more transmission events compared to the other patient groups, including preschoolers. Furthermore, adults living in the same household as children might get mainly infected by the children and not by other adults, which does however not mean that adults do not play a crucial role in introducing novel lineages into the city. Indeed, children have been previously reported to be a strong driver of influenza transmission [14].

These interpretations however are based on the analysis of influenza isolates from one season and city and will therefore need to be repeated in different seasons and cities to get a more complete understanding of the transmission patterns of influenza across age groups.

#### Conclusions

Seasonal influenza annually infects a large portion of the global population and while its global spread has been studied extensively, its local spread remains largely unstudied. Our results are based on one of the most densely sampled genetic datasets of influenza sequences to date. Additionally, we connected the genetic information to patient information such a residential address and age for all and more personal information for a lot of the patients, providing unparalleled resolution to study how influenza spreads locally. The 2016/17 season for which we collected data was dominated by influenza A/H3N2. Based on this data, we observe that hundreds of introductions initiate the seasonal influenza epidemic in the studied city of Basel, that the overall spread varies throughout the season, and that children seem to drive the local outbreaks, while elderly have their own transmission chains.

It will be interesting to see how these results transfer to other cities and seasons with potentially other social structure or other geographical location. In particular, the subtypes that circulate and their ability to escape host immunity and seasons dominated by different influenza types may influence mixing patterns. For the future, it will be particularly interesting to see if seasons where other subtypes such as influenza A/H1N1 or influenza B dominate show the same or differing patterns that we observed. While such studies on a population level requires great effort in recruiting patients as well as in sequencing viruses, they can greatly improve our understanding of how influenza spreads locally. This will hopefully allow us to streamline public health interventions in the most efficient way possible, and thus, help to reduce the great burden on societies caused by the seasonal flu.

### Methods and Material

#### Data collection and sequencing

We collected all data in the 2016/17 influenza season as described in [15]. Sequencing was performed as described in Wüthrich *et al.* [24]. Raw Illumina reads were trimmed with Trimmomatic 0.36 [25]. Alignment of paired-end reads was done by using bowtie 2.2.3 [26], using strain A/New York/18/2014 as a reference. The aligned reads were sorted by using samtools 1.2 [27]. Variants were called and filtered by using lofreq 2.1.2 [28]. Variant calling was done for sites with a coverage of at least 100. Sites with a coverage of less than 100 were assumed to be unknown and were denoted as N, that is any possible nucleotide (Details on the exclusion of sequences are described in the supplement). Exact input specification can be found at https://github.com/nicfel/FluBaselPhylo/Sequences. The consensus sequences from this study were deposited in GenBank (numbers MN299375-MN304713).

#### Timed phylogenetic tree based on the HA segment

We combined the Basel sequences with all sequences (as of the 17th of Juli 2018) from https://www.gisaid.org sampled between January 1st 2016 and December 31st 2017 for which at least the segments HA, NA and MP were available. We first aligned all consensus sequences using muscle v3.8.3129. We then built an initial phylogeny from the HA segment alone by using RaxML 8.2.1 [29] and obtained branch lengths in calendar time via timetree [30] using the nextstrain pipeline [31].

#### Initial clustering based on nucleotide differences

We then calculated the average nucleotide difference between any of the sequences and sequences from Basel. In order to split the dataset into manageable pieces, we first grouped any two sequences from Basel together if they were within an average nucleotide difference of 0.0025 per position. If the full genome for two sequences was available, this would correspond to about 32 different positions on the full genome. For an average clock rate of 2.9*10-3 per site and year, this would correspond to a pairwise phylogenetic distance of just below 1 year. Sets of sequences from Basel are only split into two groups if the two closest related sequences of each group exceeds this distance. Based on this initial grouping, we added sequences that were not from Basel to each cluster if they were at maximum 0.0025/2 mutations per position away from any of the sequences from Basel. The factor of 2 is only to reduce the number of non-Basel sequences in each of these initial clusters.

#### Phylogenetic trees of initial clusters

We next estimated rates of evolution for each genomic segment using the SRD06 model [32] and a strict clock model from 200 full genome sequences sampled in California, New York and Europe between 2010 and 2015 in Beast 2.5 [33]. These sequences were downloaded from fludb.org, were not used otherwise and are an independent dataset. We allowed each segment to have its own phylogeny in order to avoid reassortment to bias the estimates of evolutionary rates. Each of the segments, as well as the first two and third codon position was allowed to have its own rate scaler. We ran 10 independent MCMC chains each for $10^8$ iterations and then combined them after a burn-in of 10%. These estimated evolutionary rates are long-term rates of influenza A/H3N2. Since the effects of selection over short time periods are smaller compared to longer time periods. The evolutionary rates can be expected to be faster for shorter time windows [34]. We therefore expect the pairwise distances estimated for our data from the 2016/17 outbreak using these rates to be an overestimate of the actual divergence times. The xml and log files for the analysis can be found here https://github.com/nicfel/FluBaselPhylo/tree/master/EvolutionaryRates.

We next reconstructed the phylogenetic trees of all initial clusters by using the full genomes of all samples in the initial clusters. We fixed the evolutionary rates to be equal to the mean evolutionary rates as estimated previously, with the mean evolutionary rate being $2.9 * 10^-3$ per site and year. As a population prior, we used a constant coalescent model with an effective population size being shared among all initial clusters. We then estimated a distribution of phylogenies for each initial cluster, assuming that all segments share the same phylogeny. If reassortment happened, it would increase the distance between samples. Due to using fixed evolutionary rates as estimated in the previous analysis, reassortment will not bias evolutionary rates. Hence, reassortment events will increase the pairwise distance between isolates separated by reassortment, but will not bias the distance between isolates that are not.

#### Local cluster identification

To identify sets of sequences from Basel that were likely transmitted locally, we used the phylogenetic tree distributions for each initial cluster and reconstructed the ancestral states using parsimony. We made some modifications to the standard algorithm for ancestral state reconstruction. To reflect our prior belief that Basel is unlikely to act as a relevant source of influenza on a global scale, we classified internal nodes that are not exclusively classified to be in Basel as not in Basel. Since the flu season is only a few months long, we additionally assumed that lineages are unlikely to persist in Basel for more than 0.1 years without being sampled. To reflect that assumption, we classified internal nodes that are more than 0.1 years from a sample from Basel to be either in a location other than Basel, or to be in an unknown location. We then defined sequences to be in the same local cluster if all their ancestors are inferred to be in Basel. We get these local clusters for each iteration of the MCMC. The exact workflow, including BEAST2 input files can be found at https://github.com/nicfel/FluBaselPhylo/LocalClusters. While alternative model based approaches exist to reconstruct locations of internal nodes (e.g. Vaughan *et al.* [35]), these approaches themselves make strict assumptions that are violated when studying the spread of diseases on a city scale. Also, it is further unknown how well they perform when migration between individual locations is very strong.

From the grouping of sequences into local clusters as described above, sequences can be classified into different local clusters over the course of the MCMC. For the estimation of effective reproduction rates we however require each sequence to be in a distinct local clusters. To do so, we randomly picked an iteration of the MCMC and then chose the local clusters present in that iteration. In order to avoid sensitivity of the results to the iteration we chose, we repeated each analysis 10 times with randomly chosen iterations.

#### Estimation of the effective reproduction number and sampling probability

We then estimated the effective reproduction number through time as well as the sampling proportions and phylogenies from all these local clusters jointly using BDSKY [36]. We assumed the effective reproduction number to be piecewise constant in intervals of 2 days and allowed it to change every 2 days. We then assumed the difference between the log effective reproduction number in interval t (log $R_{eff}$(t)) and in interval t-1 (log $R_{eff}$(t-1)) to be distributed around N(0,$\sigma$), with $\sigma$ being estimated in the MCMC [37]. Additionally, we assume the log $R_{eff}$ at the most recent time interval and the one at the very last time interval to be normally distributed in log space around N(-0.6931,0.1). This adapted version of BDSKY is available on https://github.com/nicfel/bdsky. We assumed the rate at which an infected individual transitions to being non-infectious to be 0.25 per day. Since we followed the same procedure for inclusion of patients throughout the epidemic season, we assumed constant sampling over time. The weather data used for the correlation analysis was obtained from www.meteoblue.com. This data is based on measurements of weather stations which are then used in simulations to estimate local weather variables (see https://content.meteoblue.com/nl/specifications/data-sources).

#### Defining connectedness between individuals

We define two individuals to be connected if their pairwise phylogenetic distance is less than 0.1 years. If we assume two individuals to be isolated at the same time, this cutoff would correspond to a common ancestor that was at most 18.25 days ago. Considering that the evolutionary rates we used to perform these inferences are long term rates and therefore lower than the actual short term rates [34], we expect that the cutoff values are effectively lower in reality. This means that if we use a cutoff of 0.1 years, even individuals that are at an inferred pairwise distance of 0.1 years are very likely more closely related than that. To avoid biases originating from these cutoff values, we repeated all analyses that are based on cutoffs with of 0.05, 0.15, 0.2 and 0.3 years as the cutoff value.

#### Connectedness across age and family status groups

We estimate the average number of connections members from each of the six categorial age/family status groups have, according to the above definition of connectedness. To do so, we model the number of connections an individual from a group has as a negative binomial distribution. This allows us to model the number of connections an individual from a group has, while taking the variance of the relationship between the group label and the number of connections into account. This is in contrast to, for example, the poisson or geometric models. We assess overall model fit with an ANOVA and then perform Tukey contrasts, comparing all pairs of age groupings [38]. We correct for multiple testing by using Schaffer's method, which is similarly conservative to bonferroni, but takes into account the dependencies enforced in a linear modelling framework [39].

#### Age mixing patterns

To identify mixing patterns between the six categorial age/family status groups, we again use the definition of connection of two patients.

We use two different approaches to estimate how different groups are connected to each other. First, we use multinomial logistic regression to estimate the probability that a member from one group is connected to a member from another group. As weights, we use the inverse number of samples from each group. This implicitly assumes that individuals from each group have the same probability of being infected. Children however might have higher rates of infection, and we therefore expect this weighting to underestimate the role of children and to overestimate the role of adults

Second, we use a permutation approach. Between any two groups a and b, we compute the probability of them being associated with one another as follows: For each combination of groups a and b, we count the number of pairs that are associated with one another. We then randomly permute the age labels $10^6$ times. For each permutation, we calculate if the number of pairs between these groups is greater or smaller than what we observed. From these values, we then compute the probability that age groups a and b are positively ($P^+_{ab}$) or negatively ($P^-_{ab}$) associated with one another as:

$$\begin{aligned} P^+_{ab} &= \sum_{r=1}^{nrperm} Pairs^{observed}_{ab} > Pairs^{random}_{ab} \\ P^-_{ab} &= \sum_{r=1}^{nrperm} Pairs^{observed}_{ab} < Pairs^{random}_{ab} \end{aligned}$$

With $Pairs^{observed}_{ab}$ the number of pairs we observe in the data and $Pairs^{random}_{ab}$ the number of pairs we observe after we permute the group to patient labels. Because this test does not have an underlying model for how many connections there are between individual groups, we here use a Bonferroni correction instead of the Schaffer's method to correct for multiple testing. In order to test the sensitivity of these estimates, we repeated this analysis using cut-off values of 0.05, 0.1, 0.15, 0.2 and 0.3 years.

#### Acknowledgement

This project was funded by the Swiss National Science Foundation (SNF; grant number CR32I3_166258), the Freiwillige Akademische Gesellschaft Basel and the Swiss Red Cross. We thank the family doctors and pediatricians helping to recruit the patients for this study: Praxisgemeinschaft Dornacherstrasse (Dr. Burger, Dr. Eggenschwiler, Dr. Wyss, Dr. Gessler, Dr. Nonnenmacher), Praxis Bündnerhof (Dr. Müller, Dr. Peters, Dr. Hantke), Praxisgemeinschaft Banderet-Malè (Dr. Banderet-Uglioni, Dr. Malè), Hammerpraxis (Prof. Zeller), Praxis Schneider/von Hornstein (Dr. Schneider, Dr. von Hornstein), Davidsbodenpraxis (Dr. Amacher, Dr. Hug, Dr. Voelin, Isay, Dr. Pizzagalli, Dr. Navarini), Praxis Türkoglu/Bär (Dr. Türkoglu, Dr. Bär), and Praxis Gordon / Walker (Dr. Gorden, Dr. Walker). We also thank the study nurse team of the Clinical trial unit (Silke Purschke and Karin Wild) for their excellent organization and coordination of the patient recruitment. We thank Magdalena Schneider, Rosamaria Vesco, Christine Kiessling, Elisabeth Schultheiss, and Clarisse Straub for excellent technical assistance of genome sequencing. We thank Louise Moncla for help with the sequences assembly pipeline. We would also like to thank the laboratories which made the influenza A/H3N2 sequences used in this study publicly available on gisaid.org. The full list of the sequences used here as well as the author who made them publicly available can be found here https://github.com/nicfel/FluBaselPhylo/tree/master/Sequences/gisaid.


#### Author Contributions

TS, AE, RSS, and MB designed the overall study. NFM, TB and TS designed the phylogenetic analysis. NFM performed the phylogenetic analysis. NFM and BML performed the statistical analysis. JH made the nextrain.org implementation. ???, ???, ??? and ??? collected the data. DW, HSS and DL performed the sequencing. NFM, TS, AE, RSS and MB interpreted the results. NFM and TS wrote a first draft of the manuscript. All co-authors reviewed the manuscript draft and agreed with the final version.

### References


1. Fitzner, K. A., Shortridge, K., McGhee, S. & Hedley, A. Cost-effectiveness study on influenza prevention in Hong Kong. *Health Policy* **56,** 215–234 (2001).
2. Grenfell, B. T. *et al.* Unifying the epidemiological and evolutionary dynamics of pathogens. *science* **303,** 327–332 (2004).
3. Pybus, O. G. *et al.* The epidemic behavior of the hepatitis C virus. *Science* **292,** 2323–2325 (2001).
4. Drummond, A. J., Rambaut, A., Shapiro, B. & Pybus, O. G. Bayesian coalescent inference of past population dynamics from molecular sequences. *Molecular biology and evolution* **22,** 1185–1192 (2005).
5. Kühnert, D., Stadler, T., Vaughan, T. G. & Drummond, A. J. Phylodynamics with migration: a computational framework to quantify population structure from genomic data. *Molecular biology and evolution* **33,** 2102–2116 (2016).
6. Müller, N. F., Rasmussen, D. A. & Stadler, T. The structured coalescent and its approximations. *Molecular biology and evolution* **34,** 2970–2981 (2017).
7. Russell, C. A. *et al.* The global circulation of seasonal influenza A (H3N2) viruses. *Science* **320,** 340–346 (2008).
8. Bedford, T., Cobey, S., Beerli, P. & Pascual, M. Global migration dynamics underlie evolution and persistence of human influenza A (H3N2). *PLoS pathogens* **6,** e1000918 (2010).
9. Bahl, J. *et al.* Temporally structured metapopulation dynamics and persistence of influenza A H3N2 virus in humans. *Proceedings of the National Academy of Sciences* **108,** 19359–19364 (2011).

10. Lemey, P. *et al.* Unifying viral genetics and human transportation data to predict the global transmission dynamics of human influenza H3N2. *PLoS pathogens* **10,** e1003932 (2014).

11. Bedford, T. *et al.* Global circulation patterns of seasonal influenza viruses vary with antigenic drift. *Nature* **523,** 217 (2015).

12. Holmes, E. C. *et al.* Extensive geographical mixing of 2009 human H1N1 influenza A virus in a single university community. *Journal of virology* **85,** 6923–6929 (2011).

13. McCrone, J. T. *et al.* Stochastic processes constrain the within and between host evolution of influenza virus. *Elife* **7,** e35962 (2018).

14. Glezen, W. P. *et al.* Interpandemic influenza in the Houston area, 1974–76. *New England Journal of Medicine* **298,** 587–592 (1978).

15. Egli, A. *et al.* Identification of influenza urban transmission patterns by geographical, epidemiological and whole genome sequencing data: protocol for an observational study. *BMJ open* **9,** e030913 (2019).

16. Biggerstaff, M., Cauchemez, S., Reed, C., Gambhir, M. & Finelli, L. Estimates of the reproduction number for seasonal, pandemic, and zoonotic influenza: a systematic review of the literature. *BMC infectious diseases* **14,** 480 (2014).

17. Lowen, A. C., Mubareka, S., Steel, J. & Palese, P. Influenza virus transmission is dependent on relative humidity and temperature. *PLoS pathogens* **3,** e151 (2007).

18. Ferguson, N. M. *et al.* Strategies for mitigating an influenza pandemic. *Nature* **442,** 448 (2006).

19. Cauchemez, S., Valleron, A.-J., Boelle, P.-Y., Flahault, A. & Ferguson, N. M. Estimating the impact of school closure on influenza transmission from Sentinel data. *Nature* **452,** 750 (2008).

20. Deyle, E. R., Maher, M. C., Hernandez, R. D., Basu, S. & Sugihara, G. Global environmental drivers of influenza. *Proceedings of the National Academy of Sciences* **113,** 13081–13086 (2016).

21. Stadler, T. *et al.* Estimating the basic reproductive number from viral sequence data. *Molecular biology and evolution* **29,** 347–357 (2011).

22. Thompson, W. W. *et al.* Influenza-associated hospitalizations in the United States. *Jama* **292,** 1333–1340 (2004).

23. Troeger, C. E. *et al.* Mortality, morbidity, and hospitalisations due to influenza lower respiratory tract infections, 2017: an analysis for the Global Burden of Disease Study 2017. *The Lancet Respiratory Medicine* **7,** 69–89 (2019).

24. Wüthrich, D. *et al.* Evaluation of two workflows for whole genome sequencing-based typing of influenza A viruses. *Journal of virological methods* **266,** 30–33 (2019).

25. Bolger, A. M., Lohse, M. & Usadel, B. Trimmomatic: a flexible trimmer for Illumina sequence data. *Bioinformatics* **30,** 2114–2120 (2014).

26. Langmead, B. & Salzberg, S. L. Fast gapped-read alignment with Bowtie 2. *Nature methods* **9,** 357 (2012).

27. Li, H. *et al.* The sequence alignment/map format and SAMtools. *Bioinformatics* **25,** 2078–2079 (2009).

28. Wilm, A. *et al.* LoFreq: a sequence-quality aware, ultra-sensitive variant caller for uncovering cell-population heterogeneity from high-throughput sequencing datasets. *Nucleic acids research* **40,** 11189–11201 (2012).

29. Stamatakis, A. RAxML version 8: a tool for phylogenetic analysis and post-analysis of large phylogenies. *Bioinformatics* **30,** 1312–1313 (2014).

30. Sagulenko, P., Puller, V. & Neher, R. A. TreeTime: Maximum-likelihood phylodynamic analysis. *Virus evolution* **4,** vex042 (2018).

31. Hadfield, J. *et al.* Nextstrain: real-time tracking of pathogen evolution. *Bioinformatics* **34,** 4121–4123 (2018).

32. Shapiro, B., Rambaut, A. & Drummond, A. J. Choosing appropriate substitution models for the phylogenetic analysis of protein-coding sequences. *Molecular biology and evolution* **23,** 7–9 (2005).

33. Bouckaert, R. *et al.* BEAST 2: a software platform for Bayesian evolutionary analysis. *PLoS computational biology* **10,** e1003537 (2014).

34. Holmes, E. C., Dudas, G., Rambaut, A. & Andersen, K. G. The evolution of Ebola virus: Insights from the 2013–2016 epidemic. *Nature* **538,** 193 (2016).

35. Vaughan, T. G., Kühnert, D., Popinga, A., Welch, D. & Drummond, A. J. Efficient Bayesian inference under the structured coalescent. *Bioinformatics* **30,** 2272–2279 (2014).

36. Stadler, T., Kühnert, D., Bonhoeffer, S. & Drummond, A. J. Birth–death skyline plot reveals temporal changes of epidemic spread in HIV and hepatitis C virus (HCV). *Proceedings of the National Academy of Sciences* **110,** 228–233 (2013).

37. Gill, M. S. *et al.* Improving Bayesian population dynamics inference: a coalescent-based model for multiple loci. *Molecular biology and evolution* **30,** 713–724 (2012).

38. Montgomery, D. C. *Design and analysis of experiments* (John wiley & sons, 2017).

39. Donoghue, J. R. *et al.* in *Recent Developments in Multiple Comparison Procedures* 1–23 (Institute of Mathematical Statistics, 2004).

### Supplementary Material

#### Quality criteria for sequences used in this study

Overall, the dataset included sequencing runs from 842 unique isolates. For some of these isolated, sequencing was performed more than once. In this case, the isolate that fulfilled the quality criteria detailed below better was used. These isolates, in some cases, were isolated from the same patient, but at different time points. Bases were called to be neutral if the coverage for a site was less than 100. If a segment of a sequence had more than 20% neutral bases on more than 5 segments, the sequence was not used for further analysis. If the HA segment was not part of the 3 segments that were had less than 20% neutral bases, the sequence was discarded as well. Using the same criteria but requiring at least 4 segments, would have resulted in 736 sequences to be used. For 5 segments, 728 would have passed, 719 for 6 segments, 713 for 7 and 664 for 8. This quality criteria was most likely violated by the NA segment, explaining the drop of isolates that would have been used when requiring all 8 segments instead of 7. For patients with more than 1 isolate that passed the quality control, the sequence that was isolated earlier was used.

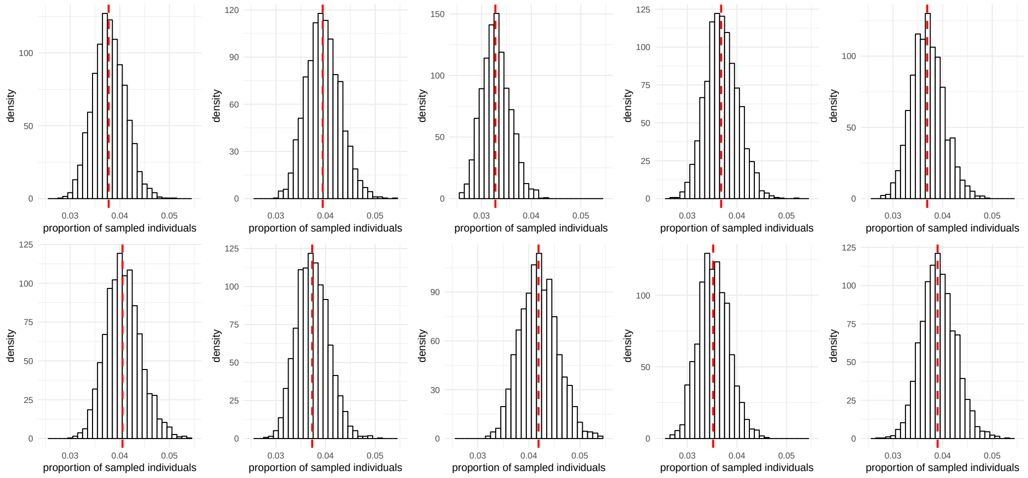


Figure S4: **Estimates of the sampling proportion for different classifications of sequences into local clusters.** Each histogram shows an inferred sampling proportion based on a classification of sequences into local clusters. Since these classifications are dependent on which iteration of the MCMC was used for the classification into local clusters, we repeated the analysis using 10 different random iterations. Each subplot shows the estimated sampling proportion when using one of these classifications. The dotted red line shows the median estimate of the sampling proportion over all iterations.

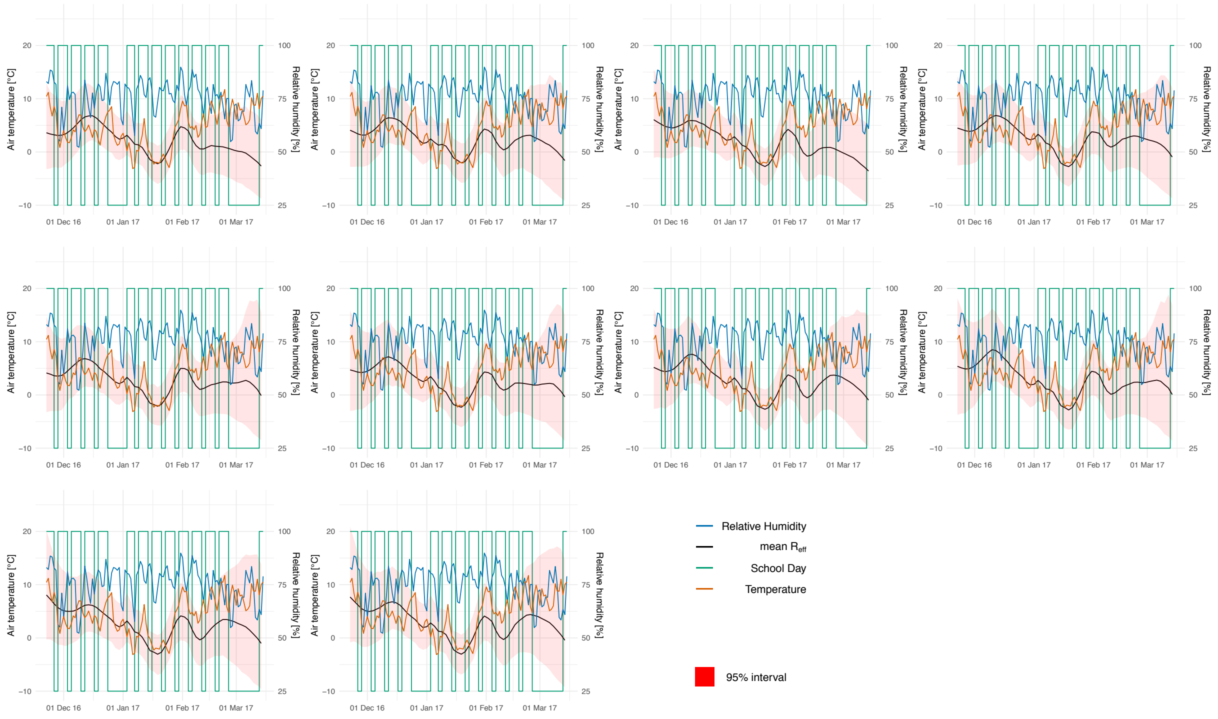


Figure S5: **Estimates of the effective reproduction number over time using different classifications of sequences into local clusters.** Each subplot shows the inferred effective reproduction number when using a different iteration of the MCMC for the assignment of Basel sequences into local clusters.

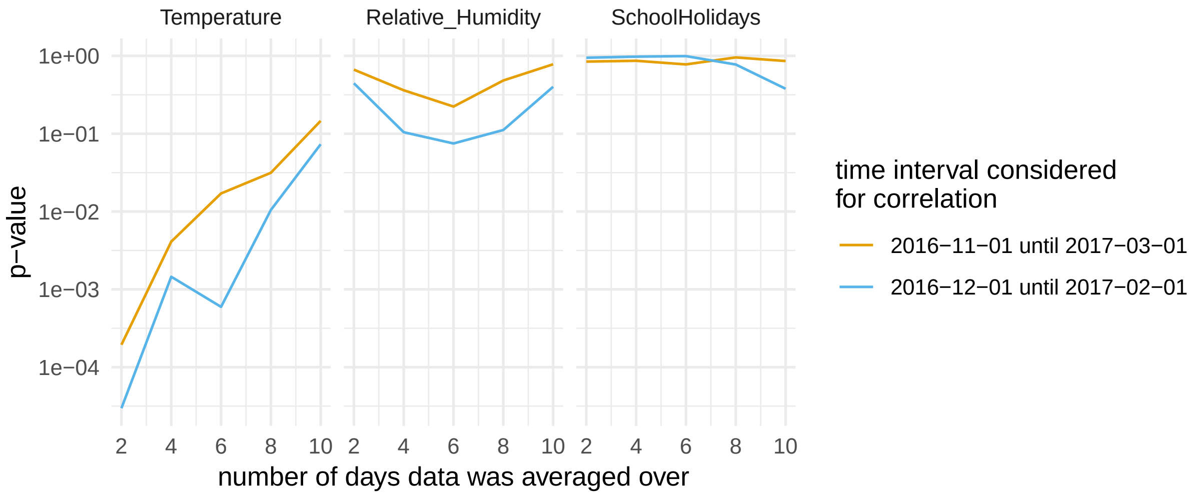


Figure S6: **Estimated p-values for correlation between the effective reproduction number and temperature, relative humidity and school days.** Here we show the estimated p-values for the correlation between the effective reproduction number and temperature, relative humidity and school days estimated when the data was averaged over different number of days (x-axis). The estimate p-values are shown for two different time intervals (in different colors). For the orange line, estimates for 1 November 2016 until 1 March 2017 were used and for the blue line, estimated from Dezember until February were used. These plots were generated using the effective reproduction number averaged over 10 different classifications of sequences into local clusters. The equivalent plots for generated using the effective reproduction number estimates of each individual subset is shown in figure S8.

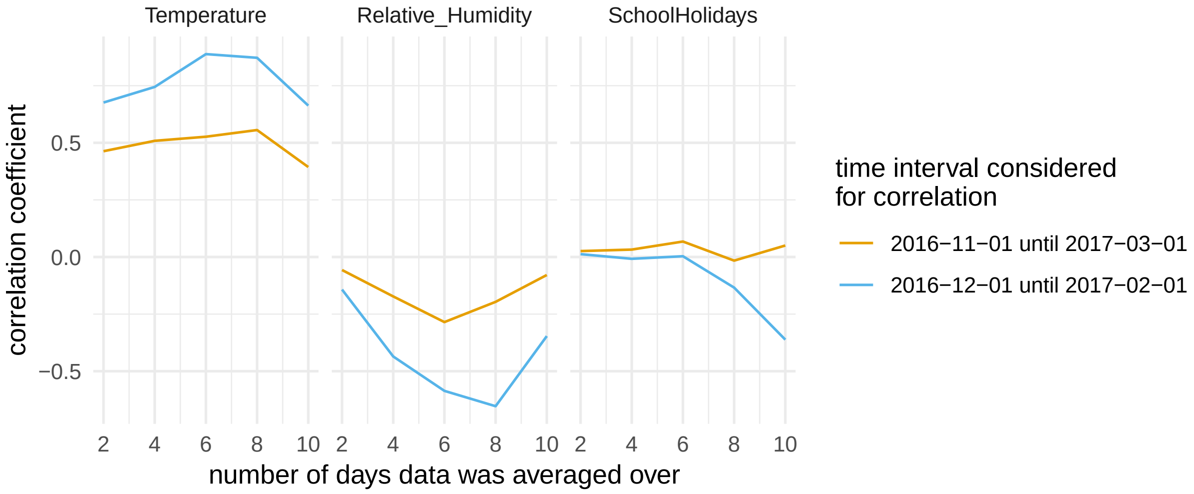


Figure S7: **Estimated correlation coefficients between the effective reproduction number and temperature, relative humidity and school days.** Here we show the estimated correlation coefficients for the correlation between the effective reproduction number and temperature, relative humidity and school days estimated when the data was averaged over different number of days (x-axis). The estimate p-values are shown for two different time intervals (in different colors). For the orange line, estimates for 1 November 2016 until 1 March 2017 were used and for the blue line, estimated from Dezember until February were used. These plots were generated using the effective reproduction number averaged over 10 different classifications of sequences into local clusters. The equivalent plots for generated using the effective reproduction number estimates of each individual subset is shown in figure S9.

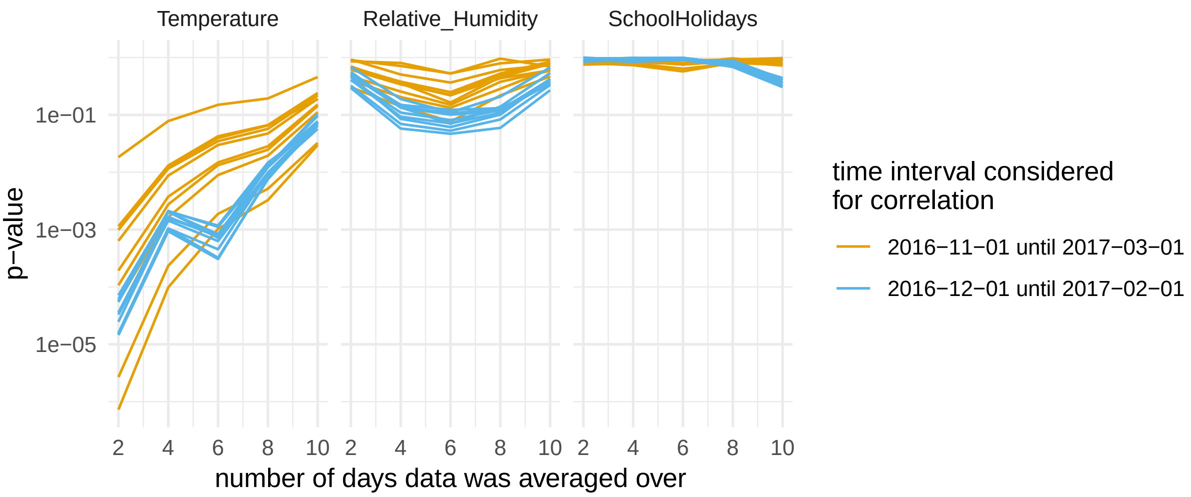


Figure S8: **Estimated p-values for correlation between the effective reproduction number and temperature, relative humidity and school days.** Here we show the estimated p-values for the correlation between the effective reproduction number and temperature, relative humidity and school days estimated when the data was averaged over different number of days (x-axis). The estimate p-values are shown for two different time intervals (in different colors). For the orange line, estimates for 1 November 2016 until 1 March 2017 were used and for the blue line, estimated from Dezember until February were used. The different lines of the same color show the p-values estimates using the effective reproduction number estimates of individual classifications of sequences into local clusters.

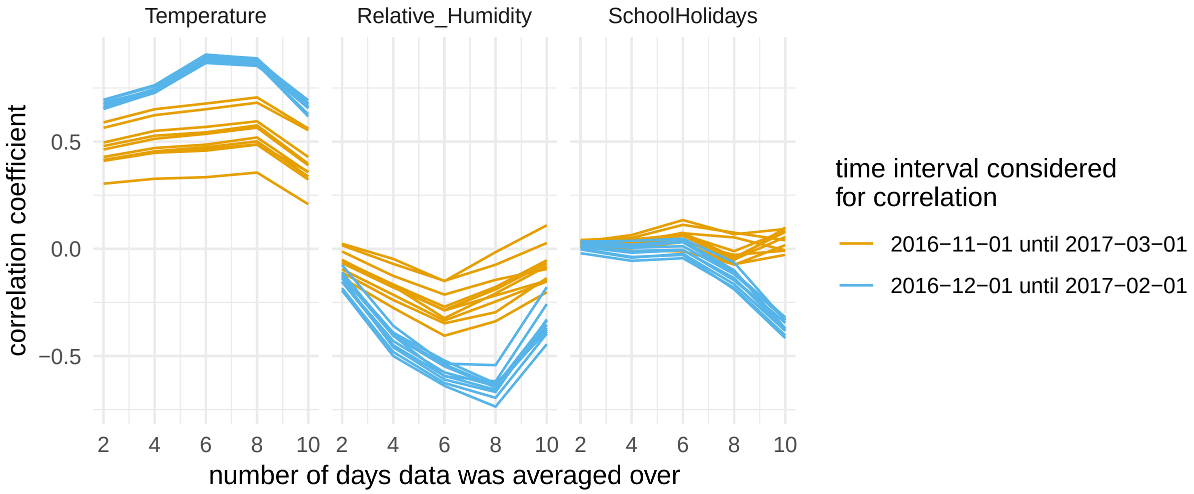


Figure S9: **Estimated correlation coefficients between the effective reproduction number and temperature, relative humidity and school days.** Here we show the estimated correlation coefficients for the correlation between the effective reproduction number and temperature, relative humidity and school days estimated when the data was averaged over different number of days (x-axis). The estimate p-values are shown for two different time intervals (in different colors). For the orange line, estimates for 1 November 2016 until 1 March 2017 were used and for the blue line, estimated from December until February were used. The different lines of the same color show the p-values estimates using the effective reproduction number estimates of individual classifications of sequences into local clusters.

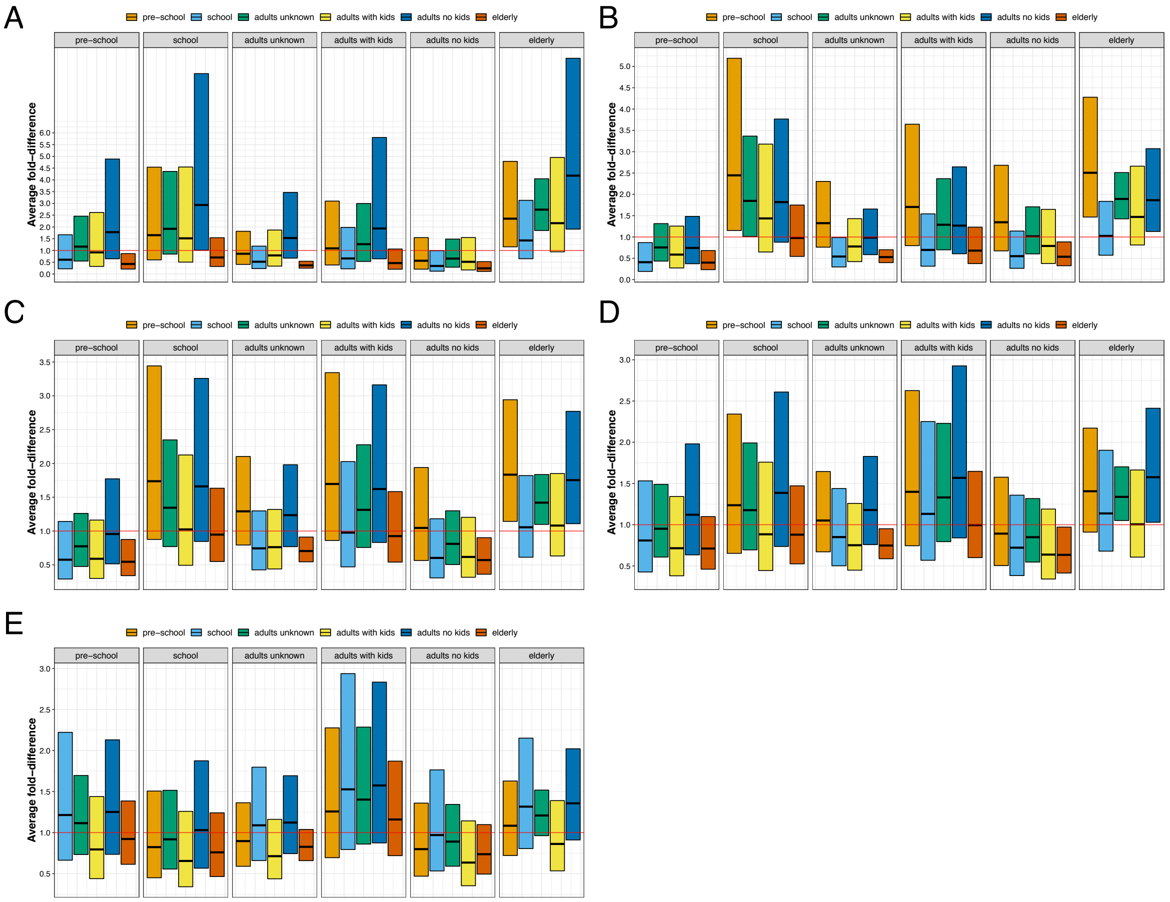


Figure S10: **Fold difference between the average number of connection of individuals from the different groups.** Plots are analogue to Fig. 2c, but for different thresholds: 0.05 years in plot **A**, 0.1 years in plot **B** (analogue to figure 2c), 0.15 years in plot **C**, 0.2 years in plot **D** and 0.3 years in plot **E**.

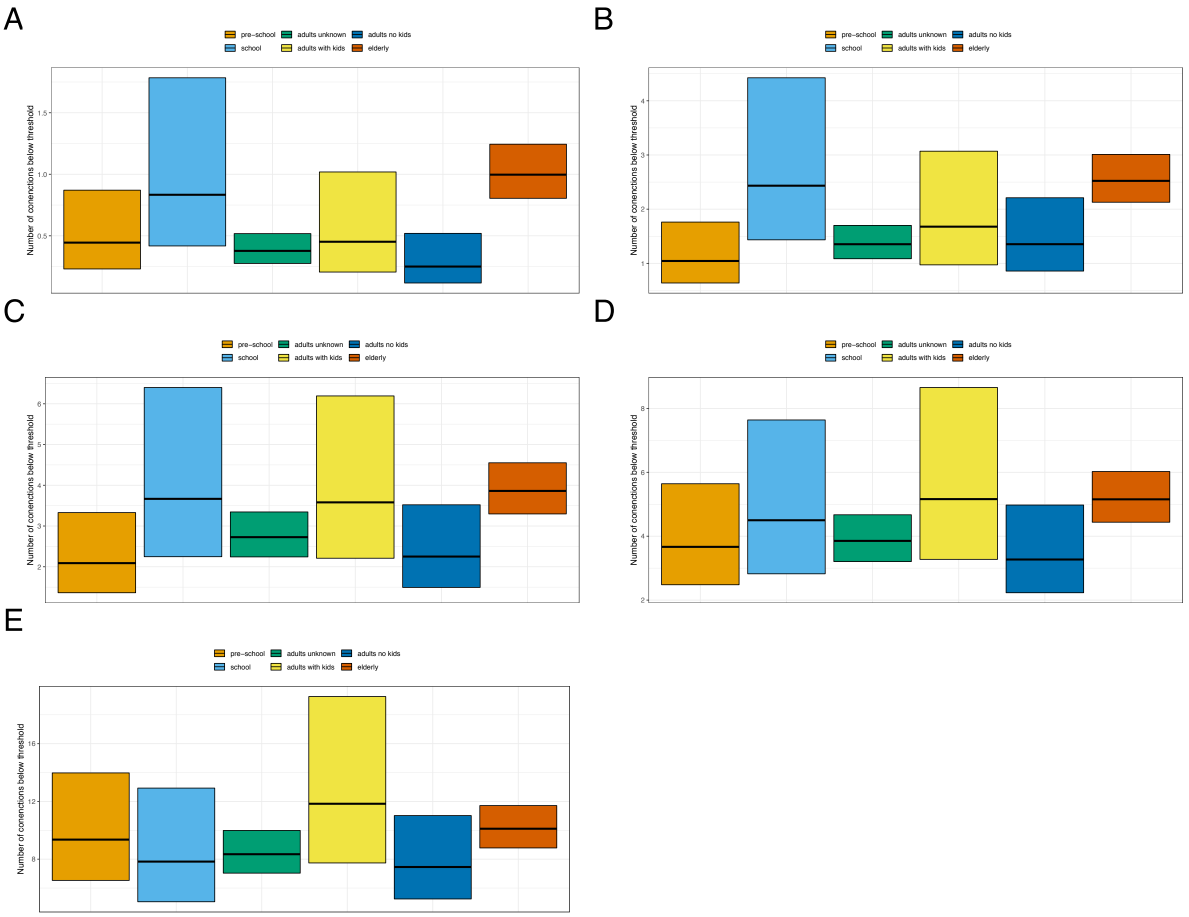


Figure S11: **Average number of patients and individual from a group is connected to.** Each subplot shows the average number of connected individuals a patient from the group shown by the color is connected to. Upper and lower bounds correspond to 95% confidence intervals around the average. We consider two patients to be connected if the pairwise phylogenetic distance between the influenza viruses sequenced from them is below a certain threshold. These thresholds are 0.05 years in plot **A**, 0.1 years in plot **B**, 0.15 years in plot **C**, 0.2 years in plot **D** and 0.3 years in plot **E**.

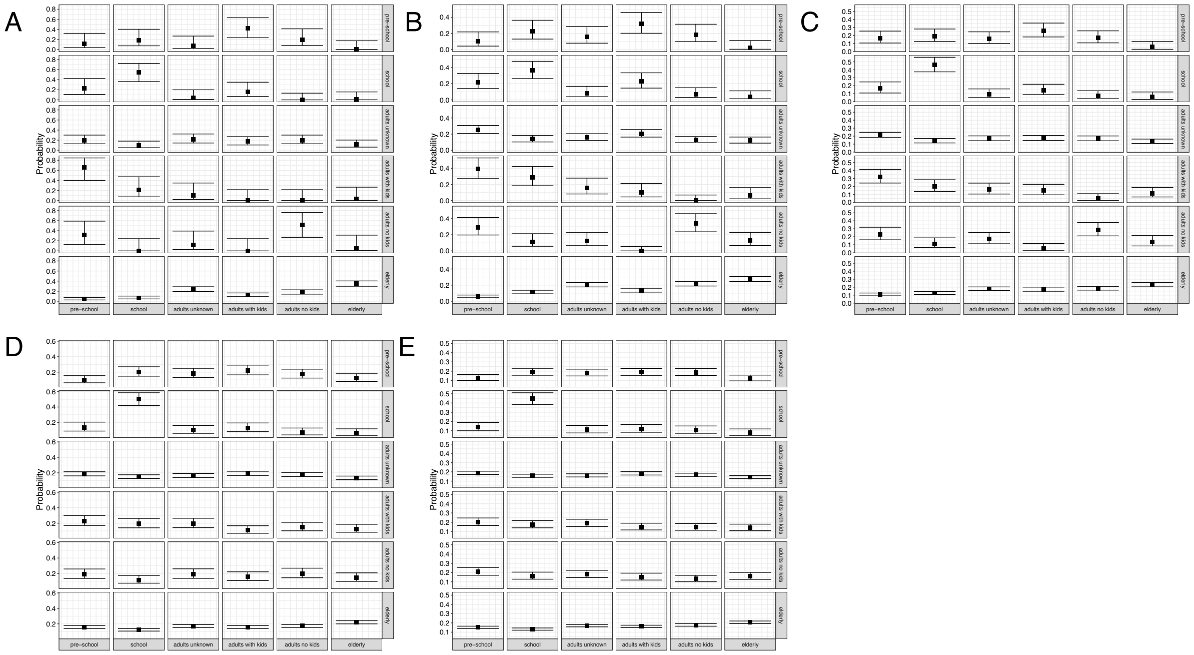

Figure S12: **Probability that an individual from the group in each row is connected to an individual from the group in a column.** Plots are analogue to figure 3a, but for different thresholds: 0.05 years in plot **A**, 0.1 years in plot **B**, 0.15 years in plot **C**, 0.2 years in plot **D** and 0.3 years in plot **E**.

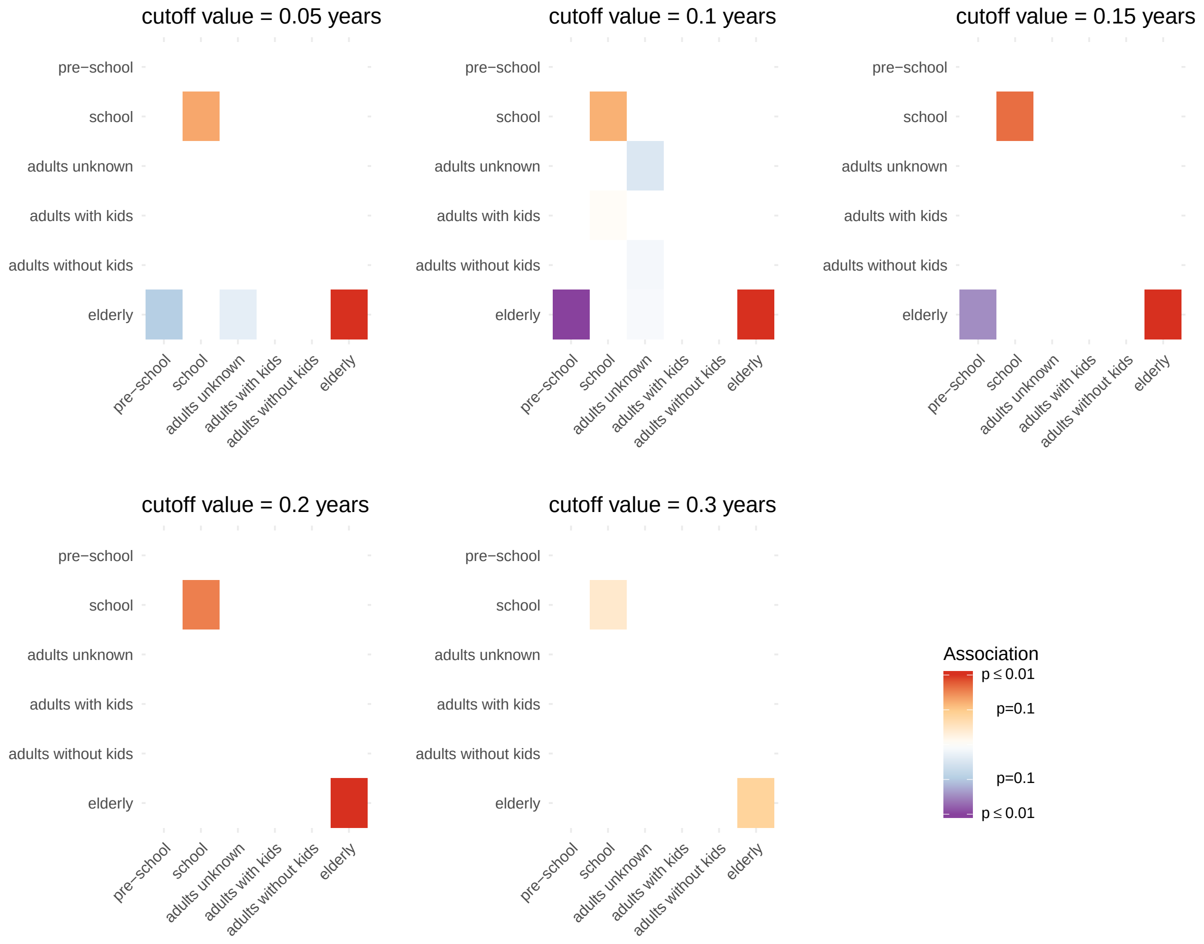


Figure S13: **Mixing of different age groups when using different cutoff values.** Plots are analogue to figure 3b, but for different thresholds. The thresholds in years are given on top of each subfigure. Unless very large thresholds are used, the elderly are estimated to be positively associated with other members from the same group. School aged children are associated with other school aged children except for very low thresholds where the number of pairs is very low. Elderly being negatively associated with pre-school aged children shows up only at lower thresholds.
